# Local interaction networks reconstructed from global biodiversity data improve pollinator restoration decision making

**DOI:** 10.64898/2026.03.30.715389

**Authors:** Teagan Baiotto, Christopher T. Cosma, Yan Yin Jenny Cheung, Desiree Narango, Jessica Woodard, Paige McCarville, Alejandra Echeverri, Grace Marguerite Horne, Eric Wood, Neal M. Williams, Katja C. Seltmann, Jesse R. Fleri, April Owens, Manuel Lequerica Tamara, Alexandra Boren, Simon Doneski, Robert P. Guralnick, Daijiang Li, Laura Melissa Guzman

## Abstract

Global pollinator declines threaten the health of ecosystems and food systems, underscoring the urgency of conservation actions such as habitat restoration. However, data gaps on plant use among pollinators continue to limit reliable design of restoration plant mixes. To address this, we present NECTAR (Network-Enhanced Conservation Tool for Analysis and Recommendation), a new modular framework that integrates multiple data modalities–including species distributions, phenological metrics, and phylogenetic data–to infer flower visitation and host plant interactions from spatial, temporal, and phylogenetic overlap, generating spatially explicit plant-insect interaction networks that guide planting recommendations for pollinator habitat restoration. We demonstrate the utility of NECTAR by generating a large plant-insect metaweb across California, comprising 2,473,729 spatially explicit interactions that included 3,792 pollinator species and 4,363 native plant species. NECTAR achieved high interaction recall across withheld interactions and independent datasets, substantially outperforming null models and matching or exceeding values reported in comparable studies. NECTAR’s data-driven plant mix recommendations are predicted to support up to 2.4 times more pollinator species compared to existing resources and random selection of plants. This optimization facilitates the inclusion of multiple goals and constraints, and provides complementary decision-making information to existing resources. NECTAR offers a scalable, evidence-based framework for translating increasingly available global biodiversity data into locally actionable restoration guidance, with broad potential to improve pollinator habitat restoration worldwide.

## Main

Global insect declines^1,2^ are compromising ecosystem functions and services that sustain human health, economic security, and ecosystem stability^3–7^. Foremost among these is pollination, a service performed largely by insects which underpins reproduction in up to 90% of flowering plant species^8^ and is integral to global food security^9–11^. With more than one in five pollinators at risk of extinction^12^, scientists have made urgent calls for policymakers, conservation practitioners, and the public to prioritize pollinator conservation^13–15^. Since the loss and degradation of native habitat is one of the primary threats to insects^16,17^, habitat restoration or enhancement (hereafter restoration) has become a central pollinator conservation strategy worldwide–one to which even small-scale actions by individuals can meaningfully contribute. Efforts to increase the quantity and quality of habitat have been shown to provide substantial benefits to pollinator communities^18–21^, including in anthropogenic landscapes^22^. The growing public awareness of insect declines and interest in pollinator gardening^23^ provides an unprecedented opportunity to engage diverse private and public land stewards in an “all hands on deck” approach to counter pollinator declines through habitat restoration.

The value of habitat restoration for pollinators, however, depends on the strategic selection of plants that meet their specific resource requirements^24^. Many insects have evolved to be dependent on narrow sets of plants^25,26^. For example, an average of 69% of caterpillar species feed only on a single plant family^25^ and 50% of bees provision offspring on pollen from only one or two plant families, with some restricted to only a single plant species^26^. While native plants are beneficial to pollinators^27,28^, individual plant taxa differ widely in the number and identity of insects they support^29^, so optimal plant mixes should reflect specific restoration goals. For instance, plantings designed to conserve individual threatened or highly specialized pollinator species may prioritize specific host plants, whereas those intended to maximize ecosystem services such as crop pollination may favor plant species that support the greatest number of pollinator species^30–32, 12^. These goals do not always align, and random selection from the regional plant species pool cannot be assumed to achieve either^33,34^. Detailed knowledge of which plants support which pollinators would greatly facilitate pollinator habitat optimization, enabling practitioners to tailor plant mixes to specific goals rather than relying on generalized assumptions about nativeness alone.

A growing array of plant lists, guides, and restoration tools have greatly improved support for pollinator-friendly planting by providing generalized recommendations for broad audiences. While these resources often reflect substantial ecological expertise, they are designed for general audiences across diverse landscape contexts, and therefore relinquish the detail and flexibility required to meet the variety of constraints and goals that practitioners and gardeners face in individual projects^35^. Translating these lists into effective site-specific plantings typically requires additional specialist knowledge, limiting the ability of the many gardeners, farmers, and other land stewards without that expertise to maximize the conservation impact of their efforts^36–38^. For instance, existing resources rarely specify which pollinator species are supported by each plant recommended or how this varies geographically, limiting their utility for projects with explicit small-scale community- or species-level restoration goals^35,39^. Due to this lack of comprehensive ecological data, such lists have been shown to provide incomplete and inconsistent planting recommendations that omit many effective species, include suboptimal ones, and favor common, charismatic, and easily identifiable pollinators (e.g., butterflies and bumble bees)^40–42^. Together, these limitations highlight the need for complementary tools that translate ecological data into context-specific, evidence-based guidance—customizable to the goals (e.g., target species conservation or ecosystem service optimization) and practical constraints (e.g., soil type, irrigation requirements, and nursery availability) that characterize every restoration, landscaping, and conservation project^39,43^.

Data-driven, ecological network-based approaches offer a promising approach to identifying plant species that support high pollinator diversity and promote community stability^44–47^, while also providing quantitative criteria to support multi-objective decision making^33,47–51^. Ecological network-based approaches have been used to optimize pollinator plant mixes at local scales^34,52^, but remain as conceptual frameworks in broader applications due to a lack of comprehensive data on biotic interactions (the Eltonian Shortfall of biodiversity data)^53,54^. Implementing ecological network-based approaches at broader scales thus requires deriving reliable interaction networks from large-scale biodiversity data that are increasingly available from community science contributions^55–59^, archived datasets, and digitized museum specimens^60–63^. However, these data can be biased toward charismatic and easily identifiable species and toward more accessible locations^64–69^, such as roads^70^. There is thus an urgent need for robust analytical frameworks that can derive interaction networks from heterogeneous biodiversity data, account for inherent biases, and translate the results into actionable, spatially explicit guidance for pollinator habitat restoration.

In response to this need, we present NECTAR (Network-Enhanced Conservation Tool for Analysis and Recommendation), a new modular ecoinformatic and statistical framework that integrates multiple data modalities–including species distributions, phenological metrics of onset and offset, and phylogenetic data–to generate spatially explicit plant-pollinator interaction networks and plant recommendations given planting constraints and objectives. To test whether NECTAR improves the pollinator conservation value of planting recommendations, we applied the framework in California, one of the world’s floral and entomological hotspots, generating one of the largest spatially-explicit plant-insect metawebs to date with over 2.2 million spatially explicit interactions for over 630,000 plant-insect pairs (including both flower-visitation and host plant interactions) across 35 Jepson eFlora districts^71^ (hereafter referred to as ecoregions). Under cross-validation, predicted networks recovered more than twice as many withheld and fully independent interactions than null models, demonstrating NECTAR’s ability to identify interactions completely absent from its training data. The resulting data-driven plant mix recommendations align with the top host and nectar plant genera recognized from independent studies. Both NECTAR recommendations and plants selected solely from observed interactions are predicted to support higher pollinator richness than existing restoration resources, supporting the value of quantitative interaction data. Our results suggest that NECTAR captures and extends this ecological signal by deliberately adding validated interactions to fill known data gaps. NECTAR also facilitates the optimization of additional goals in realistic gardening and restoration scenarios by allowing the integration of other information alongside the quantitative values of pollinator support. To enable public access to the recommendations provided by NECTAR in California, we integrated the results in the development of the Calscape Pollinator Companion tool (Box 1), a pollinator plant selection tool for habitat gardening. Because of its modularity, the framework can be extended to other regions, planting contexts, goals, and interaction types, such as at-risk-species conservation, companion planting in agriculture, or supporting natural enemies.

## Results

### Overview of NECTAR

NECTAR is a modular framework that integrates large-scale biodiversity data and ecological network modeling to identify spatially-explicit native plant recommendations that optimize ecosystem restoration goals subject to planting constraints (Fig. 1a). NECTAR is available as a pipeline (github.com/EntoDataSciCornell/NECTAR) as well as implemented for the general public in California (Box 1). We applied NECTAR to predict pollinator flower visitation and larval Lepidopteran host-plant use networks across 35 ecoregions in California. Using occurrence and interaction data, we identified likely plant–pollinator interaction pairs by filtering the set of all potential pairs using a phylogenetic constraint based on previously observed interactions, and by predicting new pairs using graph embedding with transfer learning^72^. We then computed the expected spatial co-occurrence (for both host-plant and flower visitation interactions) and overlap of key life stages (only for flower visitation interactions using flowering phenology for plants and adult flight periods for pollinators) in each ecoregion to compute an interaction plausibility score and classified likely unobserved interactions using a threshold based on the plausibility scores of previously observed spatially-explicit interactions (Fig. 1b). Finally, we used optimization approaches including genetic algorithms to maximize pollinator richness and beta diversity with a small set of plant choices. Regional interactions produced with NECTAR are predictions of plant-pollinator pairs likely to interact in at least part of a given ecoregion, and thus the top regional plant recommendations can be interpreted as plants which, on average within a focal region, support a high diversity of pollinators.

**Figure 1.**
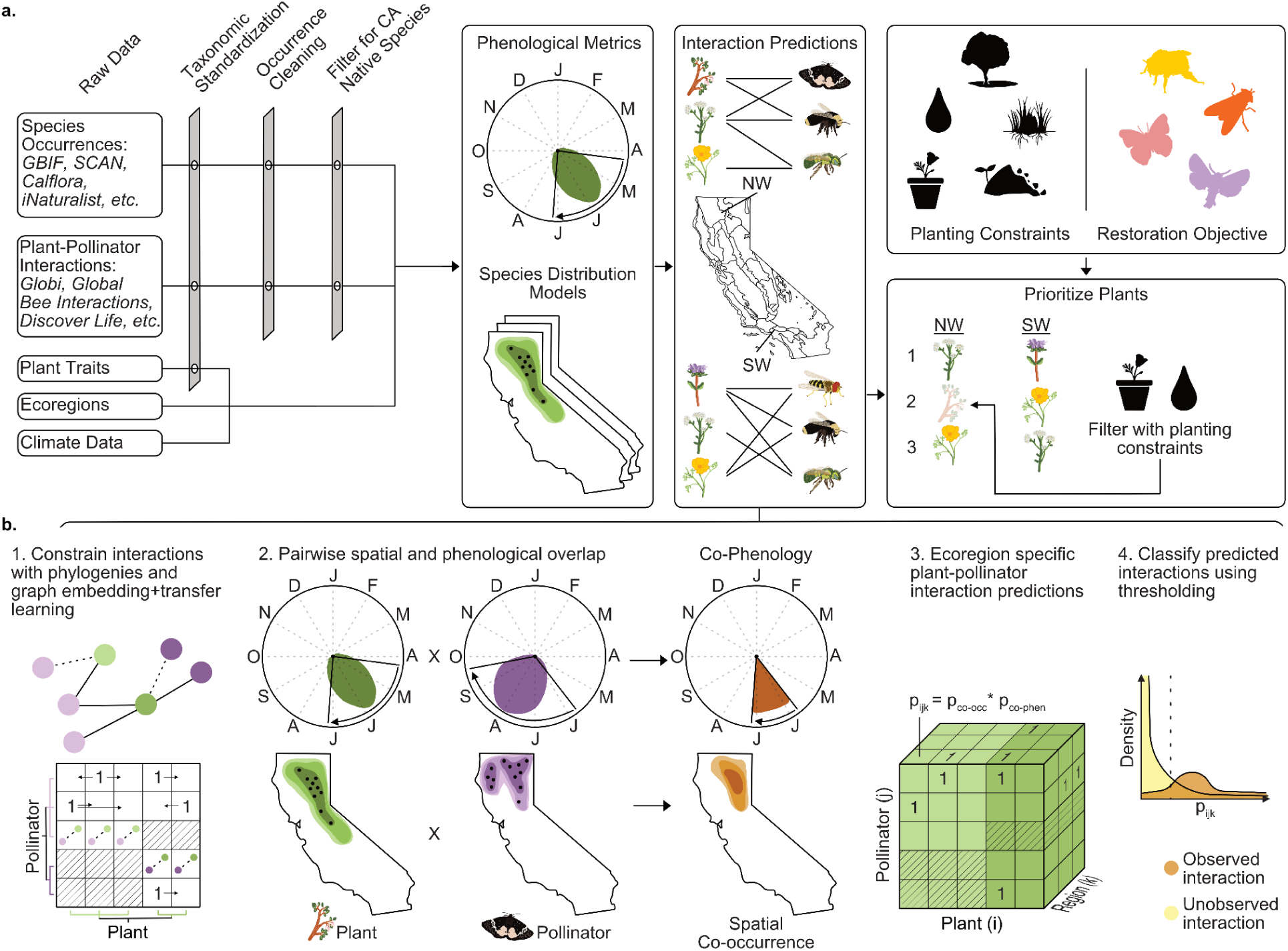
Application of NECTAR to provide regional planting recommendations in California. NECTAR’s modular design includes a data aggregation and interoperability module, a network prediction module, and an optimization module. **a.** Conceptual workflow for integrating species occurrence data, plant-pollinator interaction records, and environmental variables to predict spatially-explicit plant-pollinator interaction networks across California’s 35 ecoregions (Jepson eFlora districts). **b.** We predicted spatially-explicit networks by combining estimates of spatial co-occurrence (derived from ensemble species distribution models) with phenological overlap between plant flowering and pollinator flight periods (estimated using circular statistics). We identified potential plant-pollinator interactions using phylogenetic information (genus-level matching for taxa with existing interaction records) and/or graph embedding with transfer learning (for taxa lacking interaction data), with the primary goal of this filtering step to remove biologically implausible pairings based on observed interaction networks (b1). For all potential interaction pairs, we then calculated the spatial overlap (for both flower visitation and Lepidopteran host interactions) and phenological overlap (for flower visitation only) across all ecoregions (b2) and calculated the plausibility of a plant-pollinator interaction occurring in a given region as a function of spatial and phenological overlap (b3). Finally, we classified interactions as highly plausible using a threshold based on the 25th percentile of confirmed spatially-explicit interaction probabilities (b4; see also Supplementary Material 2 Fig. S1). These interaction networks are then integrated with restoration/planting constraints (as we exemplify below) to identify spatially-explicit plant recommendations for pollinator restoration.

### Constructing spatially-explicit plant-pollinator networks with NECTAR

We aggregated occurrence and interaction data for tens of thousands of species, resulting in 5,391 plant species and 6,242 pollinator species native to California (1,610 bees, 214 hoverflies, 281 butterflies, and 4,137 moths, Table 1). Among these species, 5,130 plants and 5,131 pollinators had at least one georeferenced and dated record in California to estimate both species distributions and phenological metrics using our tiered approach (and thus included in our interaction prediction framework, see Methods; Supplementary Material 1 Table S1 and Supplementary Material 2 Figs. S2-8). For common species modeled with ensemble Species Distribution Models (SDMs), spatial block cross-validation showed consistently strong predictive performance across taxa, with median continuous Boyce index (CBI) values of 0.644–0.813, with 97.9–99.6% of species exceeding the conservative CBI threshold of 0.25 used in previous studies^73^ (Supplementary Material 2 Fig. S9, Table S1). Ensembles of small models (ESMs) for rarer species also generally performed well (median CBI = 0.492–0.798; 67.8–96.6% ≥ 0.25), although performance was more variable, particularly for moths (Supplementary Material 2 Fig. S9, Table S1). These performance values are generally in line with or exceed those reported in other SDM studies, including large-scale insect SDMs and ESMs for rare species^74,75^.

**Table 1.**
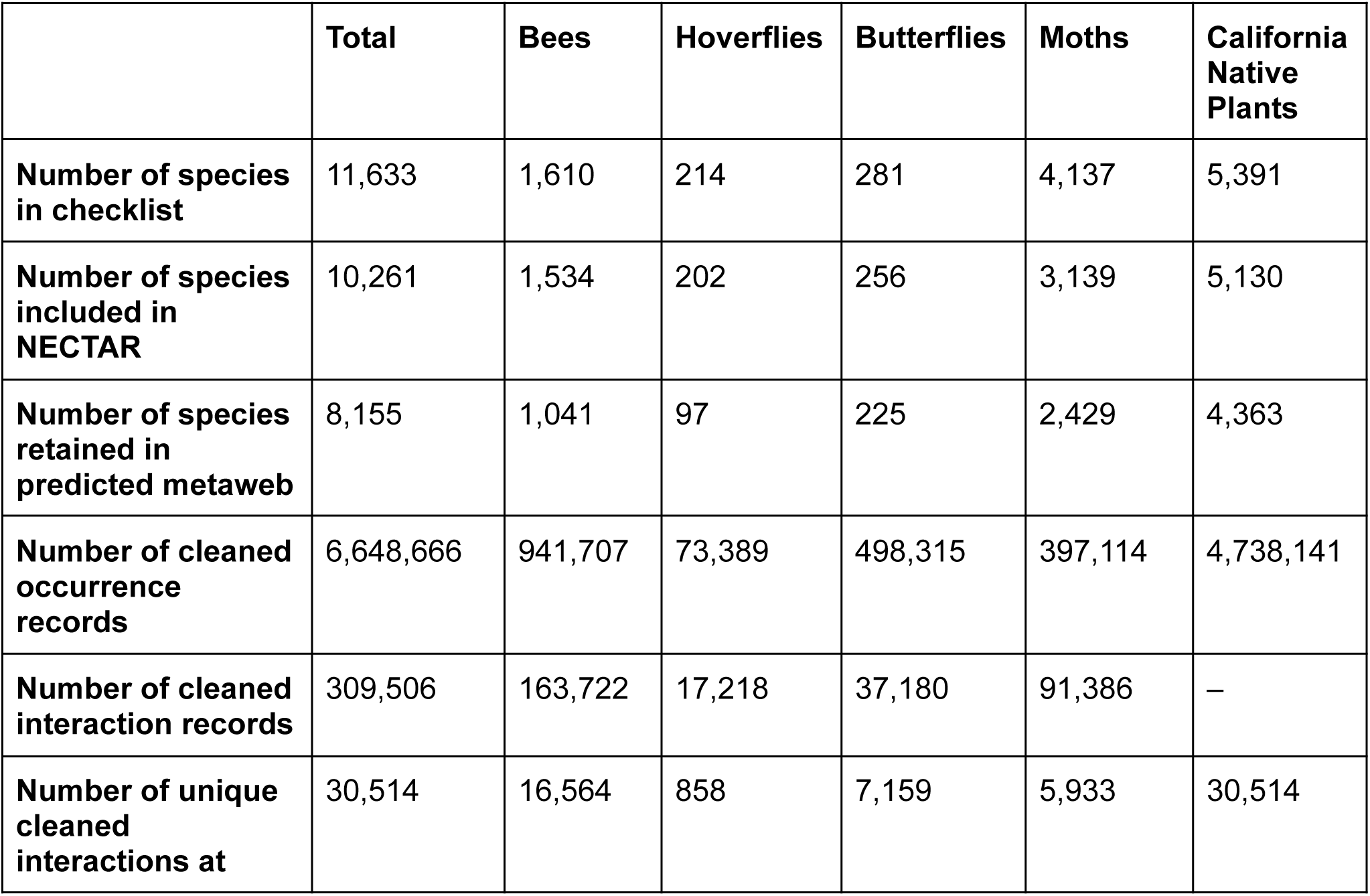
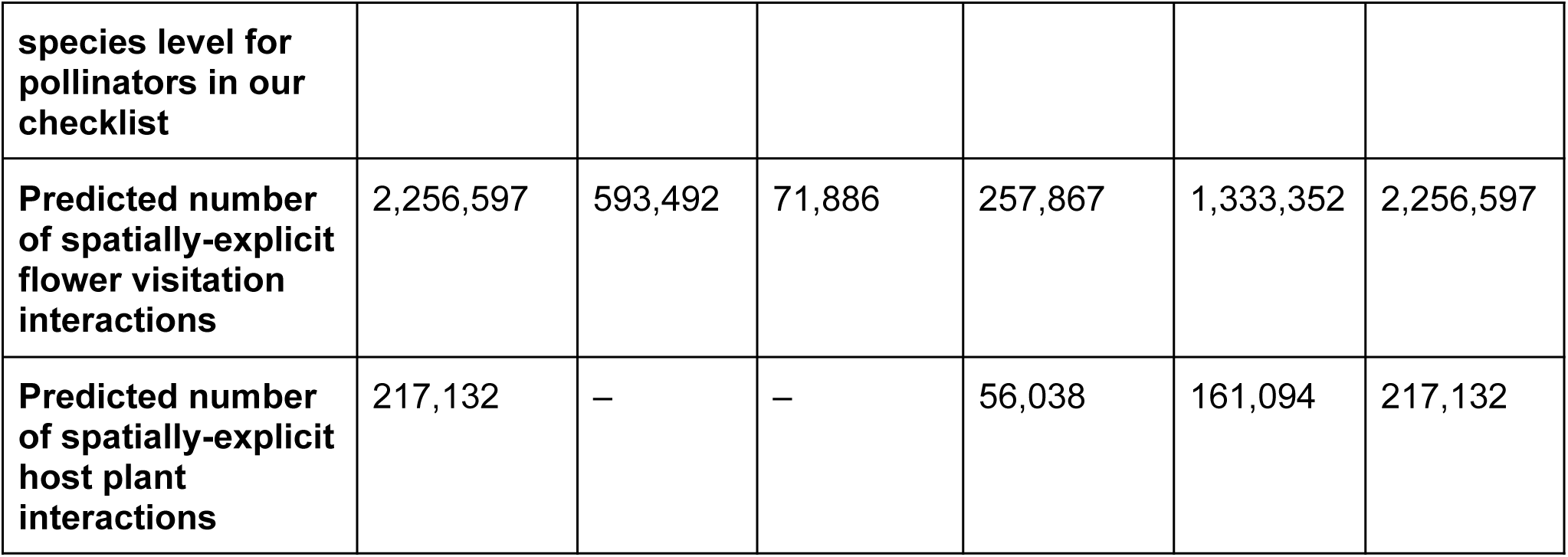
Summary of the occurrence and interaction data used in NECTAR for our analysis in California and the breakdown of interactions predicted by taxonomic group.

We compiled 309,506 unique interaction records from global biodiversity databases (e.g., Globi^60^) and through data scraping (see Methods). These records included repeated plant–pollinator pairs, but varied in the quality of geographic information, and differed in taxonomic resolution. These 309,506 records consisted of flower visitation by 2,107 pollinator species on 1,902 plant species (812 genera) and larval host plant use by 262 butterfly and 2,313 moth species on 1,539 plant species (513 genera) native to California (Supplementary Material 2 Fig. S10). In total, there were 111,416 records of unique plant-pollinator interaction pairs, of which 30,154 identified both plant and pollinator at the species level with both species native to California. The remaining interaction pairs were either at a lower taxonomic resolution (e.g., plant identified to genus level, but anything lower than genus was not kept) or either the plant or pollinator was not a known California native (Supplementary Material 2 Fig. S11).

We used the interaction prediction workflow detailed in the Methods to predict 218,175 possible unique plant-pollinator interactions at the species level for flower visitation and host plant interaction pairs between 1,530 pollinators with prior interaction data and 4,001 plants (607 genera). Predicted interaction pairs occurred on average in 4.12 ecoregions among potential interaction pairs identified from existing interaction data alone (using phylogenetic constraints, see Methods). Together with graph embedding and transfer learning (see Methods), we also predicted an additional 414,453 unique flower visitation interactions for 2,059 species of pollinators that have no existing interaction data (with 3,247 plants belonging to 438 genera). These steps alone increased the number of species-specific plant-pollinator pairs by a minimum of 20X (from 30,514 unique plant-pollinator pairs at the species level in the raw data, to 632,628 predicted). Of the initial 309,506 interaction records, 152,635 had geographic information, and of those, 5,742 were recorded at the species level for both plant and pollinator, were for species native to California, and had a georeferenced locality in California. Because our interactions are spatially explicit, our prediction space included up to 921 million potential spatially explicit interactions (5,130*5,131*35 = 921,271,050). Of these, we predicted 2,256,597 plant-pollinator-ecoregion combinations (0.24% of all possibilities) to occur, thereby increasing the number of species-specific and spatially explicit interactions by over 400X.

### Comparison of NECTAR and existing planting recommendations

We evaluated whether plants identified by our workflow improve expected pollinator support by comparing our recommendations and existing pollinator restoration resources to randomly selected regionally occurring native plants using an iterative stochastic subsampling process. In short, for every region we (i) drew a random selection of plants native to California and historically occurring in the region to provide a baseline of the minimum percentage of the pollinator community supported; (ii) drew a random mix of 10 plants from expert-based plant lists^76,77^, which were available for most but not all of California and were chosen because they are the best available resources that most closely match the goals of our framework (i.e., optimizing the pollinator value of plantings); (iii) drew a random mix of 10 plants from the recommended plants based on a metaweb of the raw interactions; (iv) drew a random mix of 10 plants from the top recommended plants by NECTAR that on their own support a high richness of pollinators; and (v) identified an optimal mix of 10 complementary plants (i.e. that together maximize pollinator richness) using a genetic algorithm within NECTAR (see Methods). On average across California, 22.9% (95% CrI [20.5%, 25.3%]) of the pollinator community would be supported when selecting plants at random (Fig. 2a - circle), 25.2 (95% CrI [13.5%, 43.9%]) when selecting plants from external resources (Fig. 2a - triangle), 39.0% (95% CrI [35.7%, 42.2%]) when selecting plants using a metaweb of raw interaction data (Fig. 2a - star), 50.4% (95% CrI [46.8%, 53.7%]) when selecting from high-degree plants at random from lists generated by NECTAR (Fig. 2a - square), and 59.9% (95% CrI [55.2%, 64.3%]) when selecting plants with the optimized approach generated by NECTAR (Fig. 2a - diamond; also see Model Summaries in Supplementary Material 2 Table S2). Because the proportion of pollinators supported based on the NECTAR networks can bias the results to support NECTAR, we re-ran the comparisons where the proportion of the pollinator community was obtained from the raw interaction networks (i.e. from only existing data). Based on raw interaction networks alone, we found that plant recommendations identified with NECTAR and the external expert-based plant lists support a similar diversity of pollinators, and are both significantly better than random (Supplementary Material 2 Fig. S12, Table S3). However, plant lists identified directly from the raw metaweb perform substantially better than either. This suggests that neither existing lists nor NECTAR’s lists are fully optimized when using only available raw data. We also found that the pollinator support quality of plants identified by NECTAR is robust to regions with low density of interaction records (Fig. 2b,c,d, mean slope of expected pollinator support w.r.t. log of interaction data density -0.16 with CrI [-0.49, 0.14] for NECTAR complimentary selection to optimize support for all pollinators and mean slope -0.11 with CrI [-0.34, 0.11] for NECTAR rank draws selection to optimize support for all pollinators), while existing resources are not (mean slope of expected pollinator support w.r.t. log of interaction data density was -0.13 with CrI [-0.15, -0.12]). This discrepancy reinforces the need to account for spatial gaps in interaction data. Further, our data-driven approach to selecting plant lists for pollinator restoration is expected to increase support across all pollinator groups included in our analysis, whether using the raw data alone or complemented with NECTAR’s predicted interactions (Fig. 2a, Supplementary Material 2 Fig. S12).

**Figure 2.**
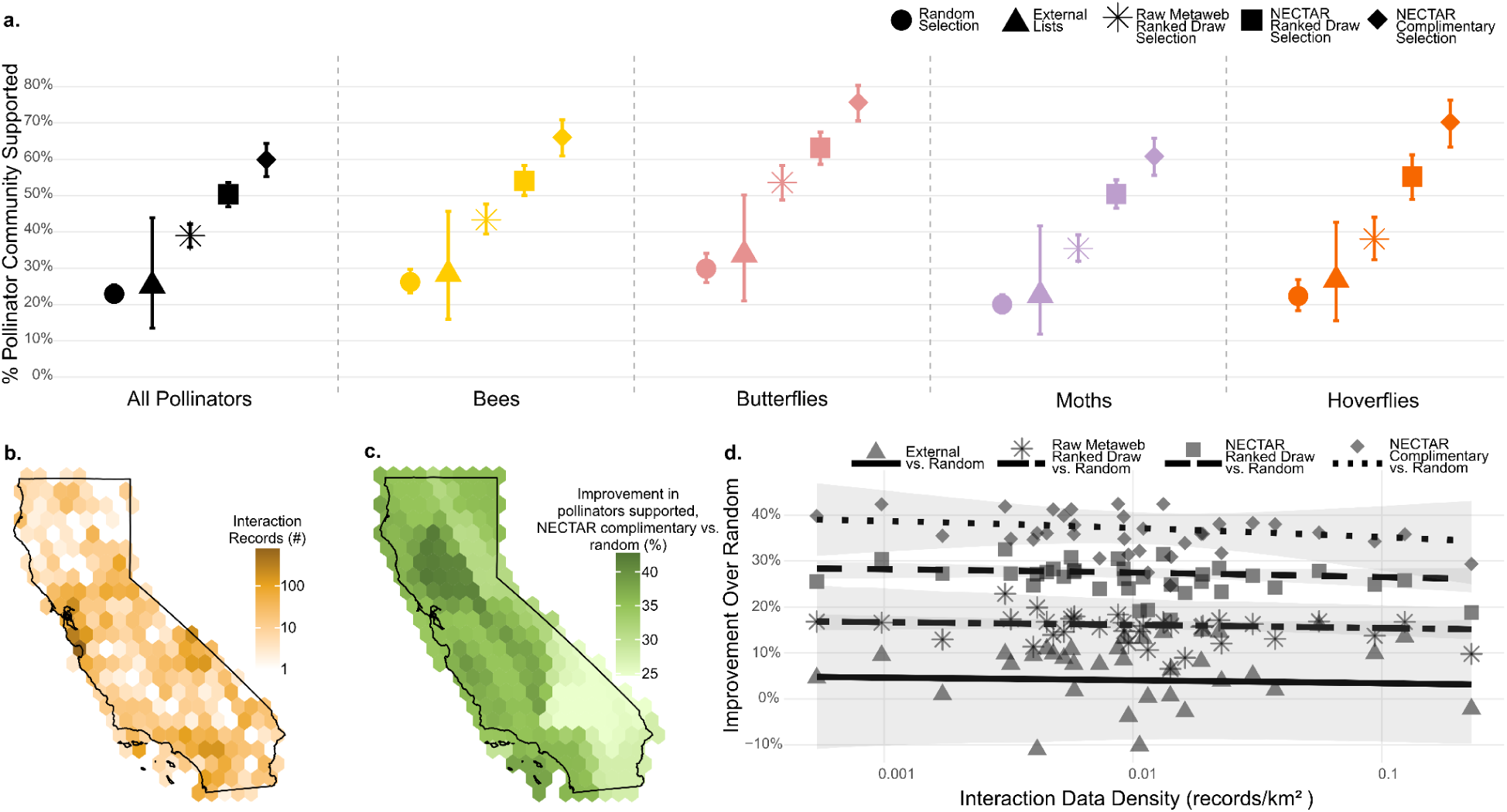
**a.** The proportion of the pollinator community expected to be supported by 10 plants selected increased as we shifted from: (i) random selection from the pool of regional native plants (circle), (ii) random draws from external expert-opinion based plant lists (triangles), (iii) random draws from the top ranked pollinator support plants we identify using a metaweb of the collated raw interaction data (stars), random draws from the top ranked pollinator support plants we identify with NECTAR (squares), and the optimal set of complimentary pollinator support plants identified with NECTAR (diamonds). When prioritizing plants for different taxonomic groups we found the same pattern, but with differing levels of support by random plants, where moths are generally the least supported while hoverflies and butterflies are the most supported. Error bars show 95% credible intervals of the mean expected pollinator support across ecoregions. **b.** The density map of spatially-explicit plant-pollinator interaction data across California was higher around urban centers such as San Francisco and Los Angeles (also see Supplementary Material 2 Fig. S13). **c.** The expected absolute increase in total pollinator richness supported by an optimized set of 10 plants compared to a random selection of plants across California was greater around areas with less interaction data. **d.** Trends of proportion of pollinator community supported across the gradient of interaction data density, showing that NECTAR (squares from the random draws of top degree plants and diamonds from the optimized selection) consistently supports a higher proportion of the pollinator community compared to random across the range of initial interaction data densities, while the improvement of the proportion of the pollinator community supported by external expert-opinion compared to random hovers around 4%. Points correspond to a comparison within one of the 35 ecoregions in our analysis, and lines show the marginal mean relationship between interaction density and increase in pollinator community support. Colors in this figure mirror color for taxonomic groups in all figures below. Ribbons represent 95% credible intervals.

### Pollinator restoration under planting constraints

We defined three hypothetical restoration scenarios with varying constraints and objectives, and used a genetic algorithm to identify the optimal set of six complementary plants (those that together maximize support for the local pollinator community) and compared expected pollinator support against random plant selection across California. However, for demonstration purposes, we visualize here for two locations: Redding (Cascade Range Foothills, top) versus Los Angeles (South Coast, bottom). For the rest of California, see Supplementary Material 3 Tables S1-S2 for the optimized plant lists and potential pollinator support. Across all scenarios, data-driven plant mixtures substantially outperformed randomly assembled mixes in terms of the diversity of pollinators supported, while still satisfying the other criteria, when compared using the predicted interaction networks (Fig. 3). In Scenario 1, our objective was to identify common plants (by removing rare plants, as defined by the California Natural Diversity Database (CNDDB)^78^) that naturally occur in Chaparral habitats which support high richness of all pollinator groups. In general, a greater proportion of bees, butterflies, and hoverflies were supported by optimized plant mixes compared to random selection of plants, with all taxonomic groups seeing significant relative improvement in support for the optimized mix (109% for bees, 130% for butterflies, 162% for moths, and 135% for hoverflies). In Scenario 2, our objective was to identify a garden plant mix composed of commercially available native plants (as defined by the California Native Plant Society) that require little water that best support pollen-specialist bees. We found that a random plant mix generally provided poor support for oligolectic bees, whereas an optimized list identified the regionally important plants for these species, on average supporting more than double the number of local oligoleges (41% of local oligolege community) even compared to the plant list optimized for all pollinators (20% of local oligolege community). For Scenario 3, our objective was to identify plants that offer both host and nectar support for declining Southwestern butterfly species^79^. On average across the state, we found that an optimized mix offers an eleven-fold increase in the number of species supported with both a host plant and a separate nectar plant compared to random (Fig. 3, Supplementary Material 3 Table S2), demonstrating that a data-driven approach can offer substantial benefit to targeted restoration efforts of imperiled species.

**Figure 3.**
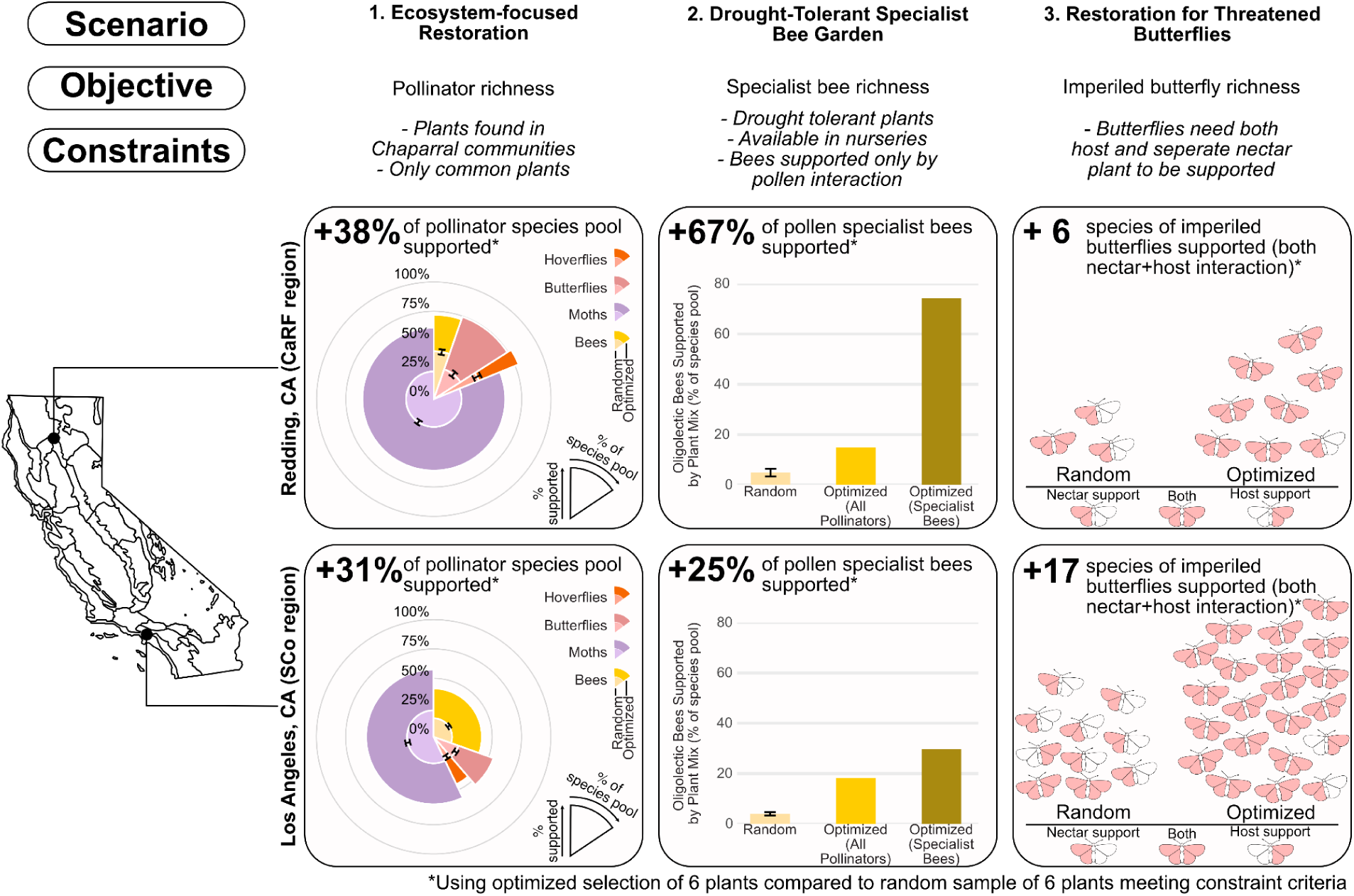
Across three restoration scenarios, we demonstrated that optimization based on pollinator support data can outperform random plant selection by 2-11X, even when both approaches use the same planting constraints (e.g., drought-tolerant, nursery-available plants). For each scenario, we quantified the improvement of data-driven optimization over random selection from the regional constrained plant pool. In (1) *Chaparral restoration scenarios* for two regions (Los Angeles/Redding), random selection left gaps in pollinator support, while optimized mixes supported a higher proportion of all taxa. Insets show grouped radial bar charts where the angular width of each slice (i.e., the width of the pie slice) represents the proportion of that pollinator group in the regional pollinator pool, and the radial extent (i.e., how far the pie slice extends outward from the origin) reflects the proportion of that taxa supported. The lighter inner slice represents pollinator support by random plant selection and the darker outer slice represents optimized selection, with slices independent (not summed) in radial extent. (2) *Specialist bee conservation*: General pollinator optimization misses key specialist supporting plants; targeted optimization for oligolectic bees nearly doubles specialist support on average. The percentage of known specialist bees supported through presumed pollen interactions by random selection, optimization for all pollinators, and optimization to support pollen interactions of oligolectic bees is shown in the inset. (3) *Declining butterfly recovery*: Random mixes can provide nectar support but miss host plants; optimization delivers 11X more host plant coverage while also maintaining or increasing nectar availability. The list of identified complementary plants for each scenario above and an unconstrained scenario can be found in Supplementary Material 3 Table S1, and the corresponding expected pollinator support for these plant mixes can be found in Supplementary Material 3 Table S2.

### How leveraging interactions enables effective restoration decision making

Our interaction prediction framework performed well under cross-validation, recovering 53.5% of withheld local interactions on average; significantly exceeding both a fully random null model (0.1%) and a phylogenetically constrained null model that assigns interactions at random among spatially co-occurring species within known host genera (25.1%; Supplementary Material 2 Figs. S14-15). Recall increased with the completeness of prior interaction data, especially for pollinators (Supplementary Material 2 Figs. S1, S16). We further validated NECTAR’s interaction predictions against four independent datasets not used by the model – coastal sage scrub surveys from San Diego (South Coast region)^80^, rare-plant pollinator surveys for plants from San Clemente Island (South Channel Islands region)^81^, opportunistic bumble bee interaction records (throughout the state from the bumble bees of North America dataset)^82^, and wildlife interactions in native plant gardens (throughout the state, from the newly created BUG project by CNPS)^83^ and found that NECTAR recovered significantly more of both regionally novel and entirely novel plant-pollinator pairs than either null model (Supplementary Material 2 Figs. S17-18). Specifically, recall of known interactions in new regions was over 90% on three datasets, and recall of novel interaction pairs was up to 60% in one dataset. Mean pseudo-precision was 2.33 (range across validation iterations: [2.23–2.43]) for withheld interactions, 4.18 for completely novel interactions from external datasets, and 4.68 for interactions novel to the focal ecoregion from external datasets. This indicates that validated interactions received 2.33–4.68-fold greater predicted support than the broader candidate interaction space. The higher recovery rate by incorporating prior information from observed interactions supports NECTAR’s ability to substantially improve inferred network structure.

Ecoregional plant rankings were generally robust to the interaction-plausibility threshold: shifting the classification threshold by ±5 percentile points changed the top 25–50 recommended plants per region by less than 10% on average (Spearman’s ρ > 0.88 in all cases), with plant ranks becoming increasingly unstable primarily when the threshold was lowered substantially below its default value of 25% and in the most remote ecoregions (Desert Mountains and North Channel Islands; Supplementary Material 2 Figs. S19-22, Table S4). Plant rankings were generally robust to alternative methods for identifying potential interactions, although sensitivity was greatest for moths, the pollinator group with the least interaction data (Supplementary Material 2 Figs. S21–22). Plant rankings were similarly robust to excluding rare species or those with poorly performing SDMs (median Spearman’s ρ = 1.0; 97.9% overlap among both the top 25 and 50 plants, respectively; Supplementary Material 2 Fig. S23).

Predicted degree of plants and pollinators in both flower visitation and Lepidopteran host use networks were positively correlated with range size and initial data availability (Supplementary Material 2 Figs. S26-29). Notably, whereas observed interaction datasets were heavily biased toward a small number of well-sampled bee genera, predicted flower visitation networks substantially expanded interaction representation for moths (i.e., the most species-rich pollinator group in our analysis); thereby capturing a broader spectrum of pollinator life histories and resource dependencies (Fig. 4a). Predicted regional interaction networks also exhibited pronounced spatial heterogeneity (Fig. 4b), driven by high turnover in plant and pollinator species composition, interaction identities, and overall network structure across ecoregions (Supplementary Material 2 Figs. S30-S41). This spatial variation fundamentally altered which plant species contributed most to supporting pollinator communities and how many pollinator species each plant supports. As a result, plant species that are highly valuable for restoration in one region can provide limited benefits in others. This provides evidence that network structure may further mediate restoration efficiency: in more nested networks, a smaller subset of plant species supports a larger proportion of the pollinator community. Together, these results show that explicitly modeling plant–pollinator interactions across space reveals spatially contingent restoration priorities that cannot be captured by plant occurrence data alone.

**Figure 4.**
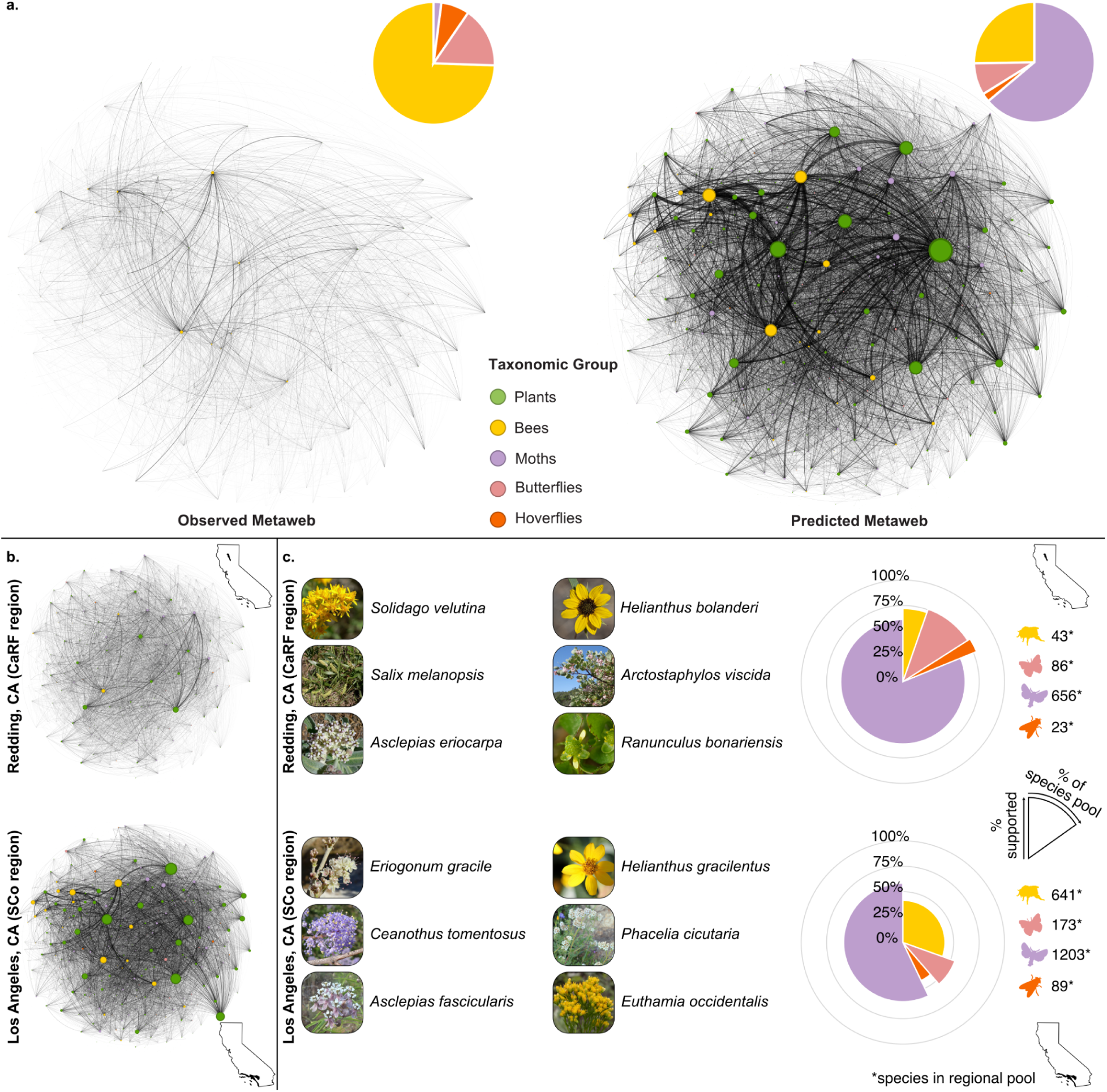
Predicted interactions expand taxonomic coverage and reveal spatial variation in network structure that helps inform location-specific plant recommendations. **a.** Observed versus predicted flower visitation metawebs demonstrate improved representation across pollinator taxa. Our predictions (right) greatly increase representation of moths (purple), the most diverse but under-sampled pollinator group, while maintaining the core interaction structure present in raw observations (left). Node size represents genus-level degree (number of species interaction partners within that genus); link width represents the number of unique species-level interactions between genera. Both networks are scaled identically for comparison with only plant-pollinator genera pairs with at least 10 shared interactions shown for visual clarity. **b.** Inferred regional networks differ dramatically in structure across California. Predicted genus-level flower visitation networks for Redding (Cascade Range Foothills, top) versus Los Angeles (South Coast, bottom) show substantial differences in network size, composition, and connectivity patterns. Redding’s network has 26% the number of interactions as the network around Los Angeles, ∼½ the number of plant species and ∼⅓ the number of pollinator species, and is less nested with a lower average degree (Supplementary Material 2 Figs. S30-32). Furthermore, moths make up a much larger percentage of the pollinator community in Redding (58% versus 39% in Los Angeles). These structural differences drive spatial variation in the identity of plants that provide the most conservation value to pollinator communities. Both networks are scaled identically for comparison with only plant-pollinator genera pairs with at least 10 shared interactions shown for visual clarity **c.** Optimal plant mixes differ by location due to network variation. The six most complementary plants for supporting local pollinator flower visitation interactions in Redding versus Los Angeles do not share any species in common. Radial pie charts show pollinator community composition (angular; i.e., how wide the pie slices are) and the proportion of each group supported by the optimized plant mix (radial; i.e.how far the pie slice extends outward from the origin). These six plants support just over ¼ of the pollinator community in both Redding and Los Angeles, but the relative makeup of who is supported differs substantially, reflecting species pools and network structure differences shown in panel b. Genus level networks visualized with Gephi^84^. Photographs in panel c from Calscape.org; *Asclepias fascicularis* by John Doyen, *Asclepias eriocarpa and Eriogonum gracile* by Neal Kramer, *Solidago velutina* by Stan Shebs, *Arctostaphylos viscida* by Scot Loring, *Salix melanopsis*, *Ceanothus tomentosus* and *Euthamia occidentalis* by Keir Morse, *Helianthus gracilentus* by Calscape, *Ranunculus bonariensis* and *Phacelia cicutaria* by Steve Matson, and *Helianthus bolanderi* by Richard Spellenberg.

## Discussion

Building on recent conceptual roadmaps^50,51^ and prior applications of plant–pollinator networks at local scales^40,85^, this study provides the first working example of how to scale a data-driven, network-based approach to pollinator habitat restoration across a large geographic region. Our novel framework, NECTAR, integrates multiple data modalities to generate spatially explicit plant–insect interaction networks that quantify plants’ ecological value to pollinator communities, which can be used to optimize plant mixes for pollinator conservation alongside other constraints. Implementing this framework using statewide data from California enabled us to generate one of the most comprehensive plant–pollinator metawebs to date^86^. NECTAR’s data-driven plant mix recommendations identified regionally important, high-value plant species overlooked by existing pollinator plant lists, increasing the richness of pollinators supported while enabling simultaneous optimization of multiple, real-world restoration objectives, including drought tolerant habitat design and targeted species conservation. We show that NECTAR’s interaction predictions are robust to spatial sampling biases and uneven sampling effort in plants by leveraging interaction information from other regions and from related plants (Supplementary Material 2 Fig. S16-18). Together, these results demonstrate that by explicitly addressing known data gaps and biases and providing quantitative criteria for decision making, NECTAR offers a highly robust and transferable framework for pollinator plant selection across regions and geographic scales.

NECTAR predicts interactions under ecological constraints designed to reduce overprediction and generate biologically plausible interactions. For example, interactions can only occur if the species overlap in space and time and there is evidence of interactions among related taxa. Our results show that incorporating these constraints improves interaction predictions, supporting the use of our approach for restoration and conservation. Mean recall of withheld interactions (0.54) was slightly higher than the median recall of 0.50 reported in a recent analysis of 217 empirical plant-pollinator networks^87^ and substantially higher than recall reported for a bipartite ant–plant metaweb (0.000–0.029 from ref.^88^). Further, although overprediction cannot be quantified directly from presence-only interaction data because true non-interactions are unknown, pseudo-precision calculations show that predicted interactions received 2.33-4.68-fold higher support relative to the background interaction space, providing evidence against indiscriminate overprediction. Although lower than the 6.55–7.17 reported by Kampe et al. (2026)^89^, their metric was calculated across all possible species pairs in a metaweb, whereas we evaluated a narrower set of plausible pairs identified from prior interaction data and further restricted to regions where both species co-occur. We also show that the threshold used for classifying spatially explicit interactions has only moderate influence on the relative rankings of pollinator support for plants (Supplementary Material 2 Fig. S19, Table S4). Finally, validating our interactions with new data showed that NECTAR performs exceptionally at identifying new locations where interactions are likely to occur, with top plant genera aligning with those previously identified from other studies (Supplementary Material 2. Table S5-6). Independent field surveys of residential yards in Los Angeles similarly found that floral visitors disproportionately used several of the top genera identified by NECTAR (e.g., *Eriogonum*)^90^. Nevertheless, predicted interactions represent informed estimates rather than direct observations, and we acknowledge that NECTAR can potentially overpredict regional interaction networks. Additional information like species traits and diel activity patterns^91,92^ could be incorporated into NECTAR’s interaction prediction steps to further reduce the chances of overprediction. As a follow up, phenological and distributional uncertainty can be incorporated explicitly to inform uncertainty of individual spatially-explicit interaction predictions.

Predicted interactions may also be skewed by sampling biases in the underlying data. Community science data is heavily biased towards easily-identifiable taxa like bumble bees or butterflies with striking patterns^64^, and while our results show that data integration and prediction helped address these taxonomic biases, there is still a need for data collection efforts focused on underrepresented pollinator taxa such as moths. NECTAR’s ability to predict interactions involving data-poor plants was fairly robust, but data-poor pollinators had both lower recall rates (Supplementary Material 2 Fig. S16) and fewer predicted interactions than those with more prior data (Supplementary Material 2 Fig. S26). Graph embedding can also be biased by uneven sampling in the source network. We reduced this risk by using each documented interaction only once, embedding plants at the genus level, and using graph embedding predictions only for pollinators lacking prior interaction data. Nevertheless, this uncertainty cannot be fully eliminated and may be particularly important for moths, the group with the fewest flower visitation interactions, and for which plant rankings were most sensitive to alternative methods for identifying potential interactions (Supplementary Material 2 Figs. S21–22). At the same time, the large number of moth interactions inferred through graph embedding is plausible since they represent the most species-rich pollinator group, with ample evidence that they nectar on a wide diversity of plant species despite being underrepresented in empirical datasets relative to diurnal pollinators^93–95^. Overall, sampling bias therefore remains an important source of uncertainty that NECTAR is designed to reduce, but cannot fully eliminate.

NECTAR reconstructs potential regional networks from a metaweb^96^, but does not attempt to downscale to realized local-scale networks. This approach assumes that species capable of interacting in one context may also interact in other regions where they co-occur spatially and temporally. However, realized networks in individual locations depend on both local-scale environmental filtering, such as filtering from urban environments^97^, as well as the biotic context, such as species relative abundances and relative interaction strengths. For example, insects may exhibit greater specialization at local scales despite being more generalized at broader scales because of relative abundances within plant communities^98,99^. Future modelling efforts at finer spatial resolution (e.g., by incorporating local abundance information and site-level biotic and abiotic conditions) could help capture these local-scale processes and bring predictions closer to true realized networks. Future work should also prioritize empirical validation of predicted interactions, particularly for those species with little interaction data, including via experimental field manipulations to assess the efficacy of our plant selection optimization criteria^100^.

The primary goal of this research is to enhance the value of habitat restoration to pollinator communities using data-driven insights. To this end, NECTAR is well-positioned to advance and complement the foundational resources provided by expert-curated pollinator plant lists. These resources^76,77^ are widely adopted by gardeners and restoration practitioners and are often the starting point for public participation in native plant gardening. In addition to plants’ pollinator value, they can reflect practical constraints including ease of establishment and local nursery availability, and are often curated via extensive trial and error and expert review. When evaluated against a metaweb of the raw interaction data, NECTAR’s optimized plant mixes were comparable to existing lists in improving pollinator support over random selection (Supplementary Material 2 Fig. S12), as neither approach is explicitly optimized to reproduce the raw observed interaction network. However, the reasons why neither perform as well as lists optimized using the raw data differ. NECTAR differs because it deliberately adds biologically plausible, validated interactions to address known data gaps, whereas expert lists’ likely differ not because of the quality of expert curation, but due to a combination of incomplete interaction information and the incorporation of additional practical constraints. When evaluated against NECTAR’s predicted networks, which address spatial and taxonomic biases in the underlying data, plant recommendations identified with the raw metaweb outperformed existing lists, and NECTAR’s recommendations substantially outperformed both, particularly in data-poor regions and for undersampled pollinator groups. Together, this indicates that, by leveraging observed interactions to fill spatial and taxonomic data gaps, NECTAR is capturing and extending ecological signal already present in empirical interaction data.

NECTAR also provides a quantitative measure of pollinator support, currently lacking in most plant selection resources. This facilitates balancing pollinator conservation with other objectives and constraints often considered in restoration and planting efforts^34,39,52,101,102^, including biodiversity goals (e.g., conserving at risk or endangered pollinators, supporting specialists), other environmental and sustainability goals (e.g., reducing stormwater runoff or improving climate resilience), social-cultural preferences (e.g., aesthetics), and practical constraints (e.g., cost, ease of establishment, space, and nursery availability). The ability to weigh pollinator support alongside these other constraints may be particularly useful for maximizing the conservation impact of wildlife gardening and habitat restoration in small spaces, where usually only a subset of the total plant species in a given ecosystem are used^103^. To illustrate this, our restoration scenarios explored the optimization of a set of six plants for projects facing real-world constraints and objectives. For example, recent reports of widespread butterfly declines in the United States^79^ underscore the urgency of butterfly conservation, which requires the coordinated provision of both larval host plants and adult nectar resources to support complete life cycles^104–106^. By optimizing plant lists using our predicted interaction networks to balance the multiple resource requirements of regionally threatened butterfly species, we increased the average number of threatened species supported with just six plants by an order of magnitude compared to random plant selection. These scenarios also illustrate that NECTAR’s optimization improves expected pollinator support even when the candidate plant pool is restricted by common practical constraints such as nursery availability and drought tolerance (Fig. 3).

### Applications and Implementation

NECTAR provides a scalable methodological approach with broad potential for global conservation applications. The R and Julia pipeline is intended for researchers and developers adapting NECTAR, and we provide complete code at github.com/EntoDataSciCornell/NECTAR. The workflow is modular: researchers can substitute regional occurrence databases, local interaction records, and region-specific spatial units while maintaining the core analytical pipeline, and can adjust settings based on data availability (e.g., stricter interaction probability thresholds for data-rich regions, coarser spatial resolution for data-poor areas). Based on our implementation in California, we found that many species had very few occurrence or interaction records. Generally, more data per species will lead to better spatial and temporal predictions; however, our use of a tiered approach enables the integration of data sparse species that may be conservation targets, which we show only slightly influenced overall planting recommendations (Supplementary Material 2 Fig. S23). NECTAR can incorporate regions with sparser data since it leverages information from closely related species and from other regions. Moreover, recent compilations of global plant-insect interactions^86,107^, occurrence datasets^108^, and advances in machine learning approaches to compile and predict interactions^109,110^ facilitates scaling this framework across regions, offering a timely approach to advance pollinator conservation through revegetation. In turn, the development of user-facing tools can enable practitioners and the public to access NECTAR’s outputs. We present the Pollinator Companion as one example of a data-driven tool for gardeners, farmers, and conservation practitioners that leverages NECTAR to improve the ecological relevance of plant selections (Box 1), and bulk data downloads are also available for researchers (doi.org/10.5281/zenodo.22214329). Because NECTAR includes biologically plausible but unconfirmed interactions, responsible deployment of such tools will require continued validation, practitioner and user feedback, and adaptation to local ecological and social contexts. To support transparency, Pollinator Companion distinguishes confirmed, previously observed interactions from those predicted by NECTAR, allowing users to interpret recommendations in light of the underlying evidence.

Beyond the examples explored here, our approach offers opportunities to advance ecological understanding and applied conservation, including: (i) advancing understanding of how ecological interaction networks and species roles vary across space and between different interaction types, which is essential for predicting and mitigating the effects of climate and land use change on biodiversity and ecosystem services^111–113^; (ii) advancing understanding of drivers of network assembly; (iii) identifying candidate plants with high pollinator value that are not currently available in nurseries, which can be included in propagation trials ^114,115^; (iv) revealing taxa, regions, and interaction types for which targeted data collection is still most needed, including identifying gaps where relatively small additions of interaction data could substantially improve network predictions; (v) modelling plant-pollinator networks to identify priority plants under future climate change scenarios, which can be used to advance climate-resilient restoration by considering both future abiotic and biotic conditions; (vi) evaluating pollinator support in existing communities, and modelling potential change under different vegetation-change scenarios; and (vii) supporting agricultural applications, including identifying companion plants for crop pollination or natural enemy support, especially given NECTAR’s modularity, which allows different interaction types and objectives to be incorporated.

#### Box 1. Improving plant selection tools with data-driven insights: Calscape’s Pollinator Companion

Translating complex ecological data into conservation action requires tools that can be adopted by diverse end-users with varying priorities, constraints, and levels of experience^116, 117^. One impactful group of stakeholders are land stewards in urban and agricultural areas, who help support pollinators through gardening and landscaping with native plants. To provide data-driven guidance to these audiences, we partnered with the California Native Plant Society to integrate our framework into Calscape’s Pollinator Companion, a pollinator gardening tool: pollinators.calscape.org/pollinator-companion/welcome. Developed through iterative user testing for accessibility to non-specialist audiences, this interactive tool brings users through a series of steps enabling them to optimize pollinator support with other planting needs or constraints (e.g., drought tolerant plants).

**Figure 5.**
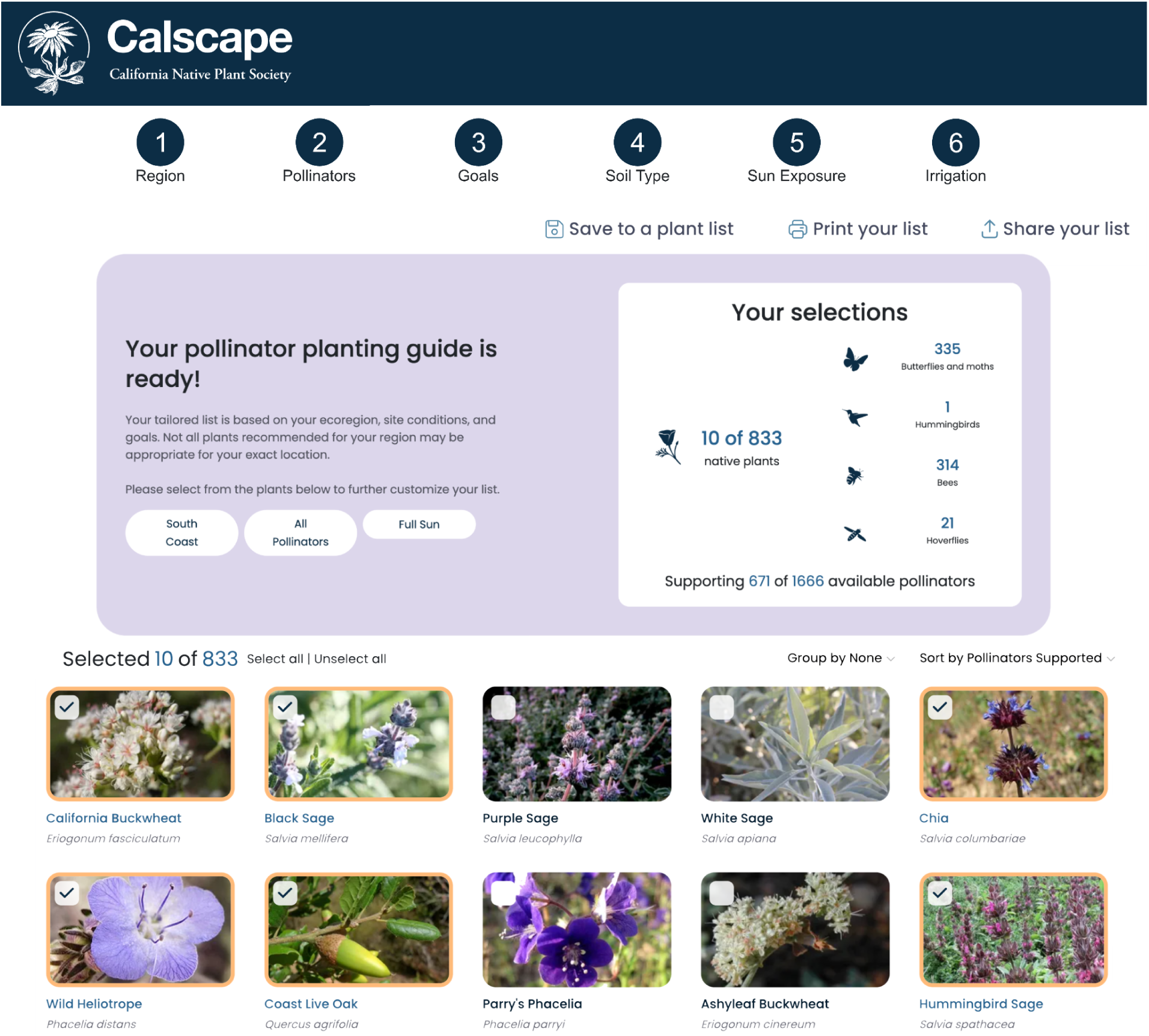
The California Native Plant Society Calscape Pollinator Companion, available through Calscape.org, presents multiple goals that users can select, including those related to pollinators and other planting priorities (e.g. improving soil health or finding crop pairings), as well as planting constraints due to soil type, irrigation amount, and sun exposure. Ultimately, the Pollinator Companion provides a list of plants that meet the constraints and maximize the user-specified goals. In the display, plants are ordered based on their value to the pollinator community, users can select plants to view lists of supported pollinators, and adding plants interactively changes the pollinator community potentially supported.

## Methods

Here, we describe a reproducible and scalable data-driven framework with R 4.3.3^118^and Julia^119^ that leverages large-scale biodiversity data to generate spatially explicit, data-driven native plant restoration recommendations for supporting pollinators (NECTAR), and demonstrate the added value of this workflow in California. We did this by first generating a metaweb of all potential plant-pollinator interactions among native plants, bees, moths, butterflies, and hoverflies using observed interaction data for both flower visitation and Lepidopteran larval host plant use. We then downscale to regional interaction networks for California’s 35 Jepson eFlora districts using a combination of spatial and phenological overlap between all potentially interacting plant-pollinator pairs (for flower visitation) and spatial co-occurrence alone (for Lepidopteran host plants) (Fig. 1). While we used the Jepson eFlora districts for our application in California, NECTAR can be used for any other spatial criteria (e.g. EPA ecoregions).

We used a multi-tiered approach for estimating both species distributions and phenological distributions of key life stage events (described below) because many species were rare and/or geographically restricted, and thus not able to be modelled with more data-intensive methods. This allowed us to include even species with a single record. While intensive modeling is not appropriate for these species, simpler approaches have been previously used in the literature for insects^120^. These rare and geographically restricted species are important to include because they may be targets of conservation, and have important ecological interactions due to the specialized niches they inhabit. Had we not included local convex-hull and buffered range approaches for estimating the distributions of rare species, or the interval/buffer approaches for estimating phenological distributions, 1,997 pollinators (572 bees, 29 butterflies, 113 hoverflies, and 1,283 moths) and 929 native plants would have been omitted from our analysis, thus removing 72,805 flower visitation interactions between 35,228 interaction pairs and 19,645 larval Lepidopteran host use interactions between 7,920 interaction pairs.

We then used these predicted regional interaction networks to identify the top pollinator-supporting plants and compared them to existing resources to demonstrate the potential improvement for pollinator restoration. Further, we designed several restoration scenarios with various goals and realistic constraints to highlight how a data-driven approach enables rapid integration of constraints and competing restoration objectives. Below is a concise description of our interaction prediction workflow, followed by our methodology for comparing the top plants identified using our predictive workflow to other restoration resources, designing the restoration scenarios, and estimating added value in each case.

### Occurrence data acquisition and cleaning

We used plant and pollinator occurrence records for all native plants, bees, moths, butterflies, and hoverflies in and within a 100 km buffer of California. Occurrence records for plants were obtained from Calflora^121^, a curated database of California wild plants, and the Consortium of California Herbaria^122^, an online repository hosting many digitized herbarium specimens across California. Occurrence records for pollinators were obtained from the Global Biodiversity Information Facility^123^, Ecdysis^124^, a curated dataset of bee occurrences for the United States^108^, and Moth Photographers Group^125^. We harmonized and cleaned all occurrence records using established cleaning workflows —namely Biodiversity Data Cleaning (bdc)^126^ and Bee Biodiversity Data Cleaning (BeeBDC)^108^. Our occurrence data cleaning consisted of removing records with missing scientific names, missing coordinates, or non-standard basis of record (we retained only human/machine observations and living, preserved, or pinned specimens); harmonizing variable names across datasets; spatial cleaning and filtering for geolocated occurrences within 100 km of California using the tigris^127^ TIGER/line delineation of the state, records with very imprecise coordinates (coarser than 1 decimal degree), transposed coordinates, or coordinates matching county centroids, capitals, and biodiversity institutions; temporal cleaning to extract collection date information from multiple columns and removal of records prior to 1900 (except for Moth Photographers Group data, since date information is not readily available for most records); taxonomic name cleaning with Global Names Parser (GNparser)^128^ taxonomic harmonization using Global Names Verifier (GNverifier)^129^ with manual corrections; removal of records not resolved to species level or outside our target taxonomic groups; removal of absence records; and record de-duplication.

### Taxonomic name harmonization

Any time that we utilized a new dataset in our workflow, we harmonized all taxonomic names. We did this by separating the list of names for plants and animals, since binomial names can be duplicated across Kingdoms, and for each, parsed the list of names with Global Names Parser (GNparser)^128^. We passed all unique parsed names to Global Names Verifier (GNverifier)^129^, returning all matches across taxonomic authorities, filtered to only matches that provide higher taxonomic information and then to those that place the species into the correct Kingdom, and from the remaining matches chose the one with the best sort score from GNverifier, which is based on how closely the name matches and the quality of the taxonomic source (manually curated authorities are valued higher than automatically curated and uncurated sources). We then cleaned the matched names with GNparser, and applied manual name corrections (e.g., removed matched species that are known non-natives or merging synonymous species not correctly identified by the matching names produced by GNverifier). Lastly, we harmonized all plant names to currently accepted names according to the Jepson eFlora^71^ using their list of accepted and synonymous plant names, first by trying to match with the original parsed names, and then by the selected harmonized name from Global Names Verifier. We used Jepson eFlora since it is widely considered the most curated and accurate authority for plants found in California.

### Species checklists

We generated species-level checklists of native plants and pollinators in each region they occur (Supplementary Material 1 Table S1) to be included in downstream modelling applications by cross-referencing our cleaned occurrence records with external lists of known non-native taxa^130–132^ and appended additional information on the rarity and endangered status of each species^78^. In all instances of merging information from external lists with our species checklists, we first harmonized all taxonomic names as described above. We filtered out known non-native species for both plants and pollinators from our checklists and occurrence dataset using ref.^130–132^. For example, we omitted honeybees (*Apis mellifera*) from the checklist of pollinators.

### Species distribution modelling

We generated species distribution models for all native plant and insect species using an ensemble modeling workflow that combined cleaned occurrence records with a set of 50 environmental predictors spanning climate, topography, soils, vegetation, hydrology, and human disturbance, at 5 arcminute resolution. We chose this resolution as a compromise between sufficient ecological resolution for estimating spatial overlap at the ecoregion scale and computational feasibility across >10,000 species; co-occurrence within grid cells was therefore treated as evidence of potential regional overlap, not a realized local interaction. We assembled predictors from WorldClim v2.1 bioclimatic variables, SRTM-derived terrain metrics, SoilGrids soil properties, ESA WorldCover land cover fractions, TerraClimate water balance variables (climatic water deficit, actual evapotranspiration, vapor pressure deficit, solar radiation), MODIS-derived NDVI and leaf area index, GPW population density, the 2018 Human Footprint dataset^133^, and distance-to-coast and distance-to-stream metrics. All layers were downloaded, processed, and aligned to a common 5 arcminute EPSG:4326 grid, with remote sensing products accessed via Google Earth Engine.

For each species, we first spatially thinned occurrences to one record per predictor raster cell. For species with >24 records, we applied environmental filtering using occfilt_env in the flexsdm package^134^, following the approach of Varela et al. ^135^ to reduce environmental sampling redundancy and associated sampling bias. For model calibration, we defined the accessible area as the ecoregions containing the retained occurrences plus a buffer scaled to data availability (25 km for <7 records, 50 km for 7–29 records, and 100 km for ≥30 records). This represents the geographic domain used to define available environmental conditions and background points against which the species’ occurrences are modeled. Within this accessible area, we extracted environmental conditions and reduced the predictor set using a two-step procedure implemented via the covsel^136^ package: (1) we removed highly correlated variables by iteratively retaining the most informative member of each correlated pair, and (2) we ranked the remaining variables using embedded selection methods within generalized linear models, generalized additive models, and regularized random forests to identify the top predictors for final modeling. We generated background points using a target-group sampling strategy, drawing from occurrence records of other species within the same taxon that fell within the species’ accessible area, thinned to one point per raster cell. A total of 5,000 background points were sampled for rare species (ESM models) and 10,000 for common species (ensemble models). We then fit models appropriate to each sample-size class: local convex-hull or buffered range models for ultra-rare taxa (1 ≤ number of occupied cells, n < 7), ensemble of small models (ESM) for rare taxa (7 ≤ n < 25), and full ensemble models (GLM, GAM, Random Forest, and MAXNET). For taxa with ≥7 occurrences, we constrained extrapolation using multivariate environmental similarity (MESS) analyses. For species with an intermediate number of records (7–100) showing strong spatial clustering (maximum/median nearest-neighbor distance >5), we applied the occurrence-based restriction method of Mendes et al.^137^, implemented in flexsdm. This post-filter removes isolated patches of predicted suitability that lack occurrences and lie beyond a species-specific distance threshold derived from the spacing among observed occurrences, thereby reducing unrealistic overprediction into geographically disconnected areas. Finally, to reduce overprediction, we masked all continuous predictions to the Jepson eFlora districts represented by each species’ occurrence records, and converted them to binary range maps using the 10% training omission threshold (OR10).

We evaluated the predictive performance of the full ensemble models and ESM models using spatial block cross-validation with 3-5 spatially separated folds. We used the continuous Boyce index as the primary performance metric, with area under the receiver operating characteristic curve (AUC), and maximum true skill statistic (TSS) as complementary measures (Supplementary Material 2 Fig. S9, Table S1). We did not evaluate the local convex-hulls/buffered range estimates for ultra-rare taxa (<7 occupied cells) because they are simple occurrence-based range estimates rather than fitted habitat suitability models. While we included all occurrence records within 100 km of California to reduce bias from truncating the niches of wide-ranging species while maintaining computational tractability across thousands of species, future efforts could improve on this by fitting fully range-wide or spatially nested SDMs ^138,139^. Outputs from our species distribution models are available at doi.org/10.5281/zenodo.22214329.

### Phenological metrics

Because many California species are active during winter when rainfall peaks, we treated the year as cyclical to avoid inflated estimates when active periods overlap the turn of the year, a common limitation of linear day-of-year methods (e.g., phenesse^140^). To estimate flowering periods for angiosperms and flight periods for adult insect pollinators, we used a three-tiered approach based on data availability at the relevant life stage. For species with >10 unique records on ≥3 unique days of the year (tier 1), we fit sine-skewed von Mises distributions^141^ to circular-transformed observation dates using maximum likelihood estimation and defined onset and offset as the 5th and 95th CDF percentiles (∼90% of observations). For species with <10 records on ≥2 unique days (tier 2), we used the minimum circular interval that encompasses all observations. For species with records on only one unique day (tier 3), we applied a ±7-day buffer. All species had phenological metrics of onset and offset estimated with only one of the tiered methods. This buffer was chosen to retain very data-limited species in the analysis while remaining conservative. Since narrower phenological overlap reduces predicted interactions in NECTAR, underestimates of phenological breadth for a given species will tend to underestimate predicted interactions. Because species could either be bivoltine or flower multiple times, we ignored the peak of the distribution and only estimated onset and offset; for those species, we are potentially overestimating duration and thus interactions.

Tier 1 estimates were calculated at four nested spatial scales (Jepson districts, regions, provinces, and statewide; requiring ≥10 observations on ≥3 unique days), with each district assigned the finest-resolution estimate available using a hierarchical approach, defaulting to successively coarser scales when district-level data were insufficient. However, for regions with sparse data, estimates were derived from aggregated data spanning multiple Jepson eFlora districts, which likely inflated estimated phenological breadth. Tiers 2 and 3 estimates were calculated statewide and applied uniformly across a species’ known range; note that tier 1 estimates for data-poor regions may also overestimate phenological breadth due to aggregation across regions. For insects, only adult records were retained; for plants, we used annotated flowering records from the Consortium of California Herbaria and Phenovision^142^, a machine learning method for identifying flowering records from iNaturalist images. Phenological estimates of onset and offset for all species-regions included in our analysis are available at doi.org/10.5281/zenodo.22214329.

### Interaction data acquisition and cleaning

We compiled flower visitation records for all pollinator taxa and host plant records for Lepidoptera from multiple sources, including the open-access databases Global Biotic Interactions (GloBI)^60^, the Natural History Museum’s HOSTS Lepidoptera hostplant database^143^, North America Lepidopteran host plant information from Shropshire and Tallamy^144^, and CropPol^145^. Using custom R scripts, we also scraped interaction records from the Discover Life website^146^, Catalog of Hymenoptera in America North of Mexico, Volume 2^147^, California Plants as Resources for Lepidoptera: a guide for gardeners, restorationists and naturalists^148^, and Pollen Specialist Bees of the Western United States^149^. Finally, we integrated butterfly host plant records supplied by Doneski et al.^150^, a dataset compiled by integrating dozens of review papers, more than 100 field guides, online databases such as HOSTS, and through new methods including unpublished lists from dozens of experts and through a collaborative effort between botanists and lepidopterists to extract host information from photos of larvae on iNaturalist. When available, we retained all location information.

With the raw interaction data, we perform a variety of cleaning and filtering steps to address variation in taxonomic names, resolution of interaction information reported, and interaction types. We first performed the same taxonomic name cleaning and harmonization steps described above for occurrence records for all plant and pollinator names, and classified interaction types into two categories (“visitsFlowersOf” for pollinator flower visitation records, and “hasHost” for Lepidopteran host plant associations) using reported life stage (i.e., larval or adult) to reclassify ambiguous Lepidopteran interaction records into flower visitation or host plant. Using the harmonized plant and pollinator identities, we removed duplicate interaction records and retained only records where both the pollinator and plant genera match with genera found in California (i.e., we retained only records where the plant and pollinator were both identified to species or genus and both the plant and pollinator belonged to genera known to occur in California). For taxon identities at a higher resolution than species, we reverted to species by removing infraspecific epithets. Thus, our final interaction dataset is a binary network of all observed interactions for genera native to California regardless of location. These cleaned interactions were then used to generate candidate predicted interactions at the taxonomic resolution of pollinator species and plant genus using phylogenetic constraints and graph embedding (described below). Interactions where both the plant and pollinator were identified to species and a locality in California was present were used to select the interaction plausibility threshold and to validate predicted regional interactions (described below).

### Predicting interaction networks

We predicted unobserved plant-pollinator interaction networks for each of California’s 35 Jepson districts by integrating phylogenetic information from known interactions, spatial co-occurrence, and phenological overlap of relevant life stages. We separated this workflow into two interaction types – flower visitation and Lepidopteran larval host plant use – because host plant use is independent of flowering phenology. For flower visitation interaction networks, we used multiple means to constrain the matrix of potential interactions to remove highly implausible pairings (forbidden interactions), each successively adding more interactions but with less certainty around the plausibility of those relationships. In the simplest sense, for pollinator species with at least one interaction record, we filtered potential interactions to all species within a plant genus with which that pollinator had previously been observed to interact with (phylogenetic constraint), a biologically informed assumption supported by evidence that closely related plants are more likely to share pollinators ^151,152^. Separately, we inferred potential interactions using graph embedding and transfer learning (described below). Unless otherwise noted, all analyses presented in this paper relied on predicted interactions for potential plant partners identified using the simple phylogenetic constraint for pollinators with some existing interaction data and from graph embedding and transfer learning for pollinators with no previous interaction data. Our predicted interaction networks are available at doi.org/10.5281/zenodo.22214329.

#### Graph embedding and transfer learning

For a given taxon group (e.g., bees), we inferred potential interactions between all native pollinator species that have phylogenetic information and plant genera that have at least one previous interaction record with this taxon group. To do so, we designed a workflow which is adapted from Strydom et al.^153^. Our workflow uses the machine learning technique graph embedding to convert an interaction network (i.e., a network of nodes and edges) into a continuous multi-dimensional space, where we can simplify to a lower dimensional representation of the network^72^. Every node (every pollinator and plant genus from the interaction data) thus has a vector of numerical values in this reduced space (via t-SVD^154^). Using their vectors, we can calculate the dot product between every pair of pollinator species and plant genus. A dot product measures how similar two vectors are; species with similar interaction patterns in the network will have a high value. We refer to this value as the interaction dot product.

Using the quantitative index, we classified each value into an interaction or non-interaction based on a threshold. A quantitative index above the threshold indicates a potential interaction, and one below indicates a potential non-interaction. To obtain this threshold, we compared observed interactions (those used to construct the network) to the predicted quantitative index, and the threshold is the value of the quantitative index that maximizes the number of observed interactions predicted to occur while also minimizing the unobserved interactions (i.e. maximizing the sum of true positive and true negative rates based on the classification result)^155^. Note that unobserved interactions could include both true non-interactions and interactions simply not detected during sampling, as there was no way to distinguish between the two.

Using this threshold, we can predict new interactions for pollinator species through two approaches depending on whether the pollinators had any observed interactions. First, for pollinators with no observed interaction data that are known to exist in the region, we assumed that more closely related pollinator species share more similar plant visitations and predicted their interactions via transfer learning (out-of-sample learning). In this transfer learning approach, we first generated a phylogeny for all species present in the checklist based on existing phylogenies. Then, we used ancestral character estimation to reconstruct the interaction dot product for the species without interaction data^153^. This resulted in a set of potential interactions between pollinator species and all species within a plant genus. Second, for pollinator species that already had interaction data, we can also readily calculate the interaction dot product for plant pairs in the interaction network that were previously unobserved (in-sample learning). While the results shown throughout only use the interactions predicted from the transfer learning, for completeness, we predicted an additional 406,892 interaction pairs for pollinators that had some previous interaction data available. We present the additional predicted interactions from in-sample learning in the Zenodo data repository.

We implemented this workflow in Julia ^119^ using SpeciesInteractionNetworks.jl^156^ to conduct the graph embedding, and PhyloNetworks.jl^157^ to conduct the ancestral character estimation. To provide the phylogenetic information required for the ancestral character estimation, we subsetted the Open Tree of Life^158^ according to the pollinator species in the checklist and interaction data. While some groups, like bees and butterflies, have better resolved phylogenies (e.g. Henríquez-Piskulich et al. (2024)^159^), we decided to use the Open Tree of Life because it allowed us to maintain a consistent workflow for all pollinator groups, and provided a standardized way to graft tips into the phylogeny. Branch lengths of all phylogenies were computed using Grafen’s method^160^ with a power transformation of 0.5. We omitted species that lack phylogenetic information from the Open Tree of Life in the transfer learning, which means species that lacked both observed interactions and phylogenetic information were excluded in transfer learning. To classify the interaction dot product into interaction and non-interaction from transfer learning, we identified an average threshold by performing a 10-fold cross-validation (90% of pollinator species in training, 10% in validation). This process was only done for species with observed interactions, and we used a rank that captures 100% variance in t-SVD from the observed interactions, meaning no observed interactions are lost in the training data. Within each fold, we identified a threshold that maximizes the number of observed interactions predicted to occur while also minimizing the unobserved interactions in the validation data. The final threshold was the average threshold value across the 10 cross-validations (Supplementary Material 2 Tables S7-10). To evaluate the performance of using this average threshold in transfer learning, we performed a 3-fold cross-validation (90% of pollinator species in training, 10% in validation) to calculate the recall rate (true positives / (true positives + false negatives)) in the validation data for each fold (Supplementary Material 2 Tables S11-14). A higher recall rate indicates that more true interactions are successfully recovered. Supplementary Material 2 Table S15 summarizes the coverage of species used and link predictions from graph embedding and transfer learning in this study.

#### Flower visitation interaction networks

For all potential flower visitation interactions–determined either by the observed interaction data, the simple taxonomic rule, or the graph embedding–we calculated the plausibility that pollinator *i* and plant *j* interact in ecoregion *k* as

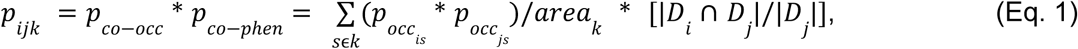

where *_pocc is_*and *_pocc js_* are the plant and pollinator occupancy probabilities for each grid cell site *s* (derived from the species distribution models produced in Step 4), *_areak_* is the total area the ecoregion *k*, and *_Di_* and *_Dj_* are the set of days that we estimate the pollinator *i* to be active and plant *j* to be flowering using the phenological metrics produced in Step 5. We calculated the 25^th^ percentile of all *_pijk_* values for previously confirmed interactions (where pollinator *i* was observed interacting with plant *j* in ecoregion *k*) and used this as a threshold to classify potential but previously unobserved interactions (*_pijk_* > threshold).

#### Lepidopteran host plant interaction networks

We used a separate and more conservative approach to predict Lepidopteran host interactions, due to the high degree of host specificity and the strong phylogenetic conservatism in these interactions. Here, we restricted predictions to spatially explicit interactions only for plants in genera already documented as hosts for a given Lepidopteran species because most Lepidoptera host plant information is available at the genus level. We calculated the plausibility of Lepidoptera species *i* using plant *j* as a host in ecoregion *k* as *_pco_*_−*occ*_ , the normalized spatial co-occurrence term alone, since host plant use can occur across the year and is independent of the flowering phenology term used for flower visitation. As with flower visitation interactions, we applied a threshold based on the 25th percentile of *_pijk_* values for previously confirmed host interactions to distinguish predicted interactions from those unlikely to occur (Supplementary Material 2 Fig. 1B).

### Validating predicted interactions and plant recommendations

We used three complementary approaches to assess the reliability of our predicted flower visitation interaction networks and the robustness of plant recommendations derived from them: cross-validation against withheld spatially explicit interactions, validation against fully independent external datasets not included in the analysis, and sensitivity analysis of plant rankings to the interaction-plausibility threshold. We did not validate Lepidopteran host networks using this approach due to the sparsity of spatially explicit host plant interaction data; however, we assume our host plant predictions are reasonable since we restrict them exclusively to known host genera.

First, for flower visitation networks, we iteratively withheld 20% of unique spatially explicit interaction pairs (from the 5,742 spatially explicit interactions that were recorded at the species level for both plant and pollinators, were for species native to California, and had a georeferenced locality in California), predicted spatial interaction networks using Eq. 1, classified interactions as probable or improbable using the thresholding approach described above, and calculated the recall rate (TP/(TP+FN)) for the withheld interaction pairs. We compared this recall rate to two null models: interactions drawn at random from both the full potential interaction pool (any combination of plant and pollinator species in that ecoregion), and interactions drawn at random from the filtered pool remaining after removing forbidden links based on previous pollinator species-plant genera interaction pairs. For each of 25 iterations of withholding data, we generated 100 draws from each null model and calculated the proportion of null draws that recovered as many or more of the withheld interactions as our predicted network. We used recall rate rather than accuracy because we cannot distinguish true negatives (interactions that genuinely do not occur) from interactions that simply have not yet been observed. We excluded the graph embedding step from this validation procedure because withholding a portion of spatially explicit interaction data would be unlikely to eliminate many unique pollinator species–plant genus interaction combinations, instead validating this step independently (see above). We additionally examined how recall varied with respect to the total number of interactions recorded in our collated interaction dataset by plant and pollinator taxa.

Second, we compared our predicted regional interaction networks to four independent datasets with spatially-explicit plant-pollinator interactions in California, covering coastal sage scrub habitat in San Diego^80^, rare-plant pollinator visitation interactions on San Clemente Island^81^, bumble bee interaction records through the state of California^82^, and biodiversity interactions in California urban gardens^83^. We filtered these datasets to interactions involving only plant and pollinator species included in our checklist, and further subsetted them to (i) interaction pairs that were not recorded in the focal ecoregion and (ii) interaction pairs that were not recorded at all in our collated interaction dataset (interactions already observed in the focal region would have a recall rate of 100% as per the design of our reconstructed interaction networks). Using the classification threshold from the full predicted network (since no data were withheld for this analysis), we calculated recall rates for each novel interaction set and compared them to the same two null models described above.

Third, to quantify the robustness of plant recommendations to the interaction-plausibility threshold, we recalculated each plant’s predicted degree at six alternate thresholds (10th, 15th, 20th, 30th, 35th, and 40th percentiles of confirmed interaction scores), in addition to our default 25th-percentile threshold. For each ecoregion, we compared the percentage of the top 25 and top 50 plants (ranked by degree) that were shared between the default and each alternate threshold, and calculated Spearman’s rank correlation between plant rankings at the default and alternate thresholds, both restricted to the top 25/50 plants and across all ranked plants. We additionally tested sensitivity to (i) ultra-rare taxa (<7 occupied cells, which were represented by simple range buffers rather than SDMs), by excluding these taxa and their interactions; (ii) poorly performing species distribution models, by removing all species with CBI <= 0.25 and their associated interactions from each regional network and; (iii) the method(s) used to identify potential interaction links, by recalculating plant degree and comparing plant rankings with the full networks using the same rank-correlation and top 25/top 50 overlap metrics.

To assess potential overprediction, we calculated pseudo-precision following Kampe et al. (2026)^89^ for (i) the leave out validation and (ii) novel interactions identified with the four independent datasets as the mean interaction plausibility score of withheld or novel interactions divided by the mean interaction plausibility across all candidate pairs (for the leave out validation, this is constrained to phylogenetically plausible candidate pairs in each ecoregion; and for the external data validation, this also includes candidate pairs identified with graph embedding for pollinators with no prior interaction data). This metric evaluates whether validated interactions are preferentially assigned higher probabilities than the broader candidate interaction space, providing a relative measure of discrimination when true non-interactions are unknown^89^.

### Comparing to existing planting resources

We compared potential pollinator diversity supported by our data-driven approach against two existing spatially explicit plant recommendation resources–Pollinator Partnership^77^ (which uses Bailey Ecoregions^161^) and the Xerces Society^76^ (which uses custom boundaries that we digitized in QGIS^162^; see Supplementary Material 2 Fig. S42)--and against random draws from plants native to each region. All comparisons used mixes of 10 plants selected by one of five methods:

(1) random native plants, (2) random draws from Pollinator Partnership or Xerces Society regional lists, (3) random draws from top-ranked plants derived from a metaweb of the raw interaction data, (4) random draws from NECTAR’s top-ranked plants, and (5) a complementary set optimized by NECTAR’s genetic algorithm (described below). Because Jepson, Bailey, and Xerces boundaries are not perfectly congruent, we subdivided all regional boundaries into unique combinations of the three classification systems (Jepson eFlora districts, Bailey Ecoregions, and Xerces regions).

For methods 3, 4, and 5, we identified top-ranked plants as the N highest-degree species (those supporting the greatest pollinator richness) from either a metaweb of previously observed interactions (3) or predicted flower visitation networks for each Jepson district (4 and 5), where N matched the length of the external plant lists in that region. We then either randomly sampled 10 plants from this list or selected the complementary optimized set using a genetic algorithm.

Pollinator support was defined as the number of pollinator species with at least one interaction among the sampled plants.

To quantify variance in the estimates of pollinator support ^163^across plant selection methods, we fit Bayesian mixed-effects models with brms^163^ for each of five pollinator groups (all pollinators, bees, hoverflies, butterflies, and moths) as

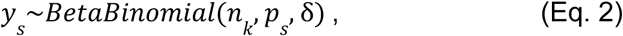

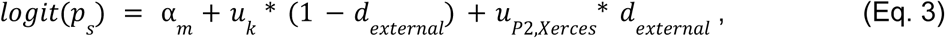

where *_ys_* is the unique number of pollinators in the focal taxonomic group supported by the *s*th plant sample, *_nk_* is the total number of pollinators for that group in ecoregion k in which sample *s* was drawn from, p_s_ is the expected proportion of the pollinator community supported by sample s, and δ is the beta-binomial overdispersion parameter. α*_m_* is the fixed intercept for the method *m* used for selecting that plant mix. Lastly, to control for spatial non-independence of samples drawn, we include *_u_*_k_ , which is a random effect of Jepson eFlora district *k*; *_uk_* _*P*2,*Xerces*_ , a multi-membership random effect of the Pollinator Partnership and Xerces Society regions; and *_dexternal_* , a dummy variable indicating which samples were drawn from the external plant lists so the random effects can be appropriately attributed.

We also ran a separate model where the linear equation was defined as

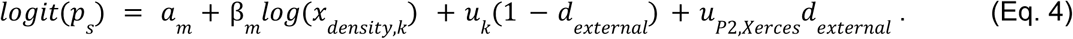

Eq. 4 includes β*_m_* , a fixed slope of interaction data density in ecoregion *k* (*_xdensity_*_,*k*_) with respect to the method used to select the plant mix in sample *s*. We included this model separately to assess whether the quality of plant choices varied in any sampling scheme by the initial density of interaction data. In all models, we used weakly informative priors. Specifically, we assumed an prior normal distribution centered at 0 with a standard deviation of 1 for the fixed effects of selection method, a normal distribution centered at 0 with a standard deviation of 0.5 for the method-by-interaction-density slopes, and a generalized Student’s t-distribution with 3 degrees of freedom centered at 0 with a scale of 2.5 of the ecoregion and external-list-region random effects. Because all priors on method effects were centered at zero, they did not assume any directionality in how the selection method would affect pollinator support a priori.

It is worth noting that using the predicted interaction networks from NECTAR as the basis for comparing plant mixes from different methods assumes plants only support pollinators in regions they are found natively. Thus, plants recommended by external lists outside of regions they are known to occur in the wild can result in no predicted pollinator support. We also repeated this comparison using a metaweb of observed plant-pollinator interactions (See Supplementary Material 2 Fig. S12), assuming any pollinator recorded in a Jepson district can interact with a sampled plant if that interaction is documented anywhere in collated interaction records. In both instances, we matched external plant lists to the interaction datasets at the species level, meaning we may have overemphasized the value of cultivars suggested by external lists to pollinators, since cultivars often support fewer pollinators than the species they are derived from^164^.

### Restoration scenarios

To test the utility of our approach under realistic conditions, we developed three restoration scenarios, each with distinct objectives and constraints, and compared the pollinator support achieved using our data-driven approach versus random plant selection without interaction data (as our null model) from the ecoregional species pool that otherwise meets the scenario-specific constraints. In each scenario, we identified the top 10 plants that together maximize the stated objective using a genetic algorithm. The genetic algorithm is similar to that proposed by M’Gonigle et al.^52^ and iteratively evolved 200 “populations” of 10 unique plants (among the constrained suitable plant pool) over 200 generations. In each generation, the algorithm created new plant combinations by selecting the high-performing combinations of parents (plant sets from the previous generation), combining the plants from these parent lists, and occasionally replacing plants with randomly selected alternatives (according to a mutation rate of 0.1). We compared these optimized plant lists to the average pollinator support of 100 draws of 10 randomly selected plants from the corresponding Jepson district’s species pool that met all planting constraints. For all scenarios, we visualized the proportion or number of species of the focal pollinator community supported by the optimized plant lists compared to random selection in two separate regions: Los Angeles (South Coast district) and Redding (Cascade Range Foothills district). However, we ran the analysis for all regions in California. The list of identified plants and corresponding pollinator support is available in Supplementary Material 3 Tables S1-2.

#### Scenario 1: Chaparral community restoration

This scenario aimed to maximize pollinator diversity using plants suitable for chaparral community restoration that are relatively common and feasible to collect for seed-based restoration. We constrained candidate plants to those associated with chaparral community (using Calflora’s plant community association data^121^) and excluded rare or endangered species by removing all plants with California Rare Plant Rankings of 1–3^78^. The genetic algorithm’s fitness function maximized the number of unique pollinator species (bees, butterflies, moths, and hoverflies) supported by flower visitation interactions across the selected plant combination.

#### Scenario 2: Drought-tolerant specialist bee garden

This scenario optimized support for oligolectic (pollen specialist) bee diversity in a drought-tolerant garden context. We constrained candidate plants to species requiring low water usage and available in at least one California nursery using Calscape plant data^165^. We ran two separate optimizations: one that maximized overall pollinator diversity (all flower-visiting species), and another that specifically maximized oligolectic bee diversity. For the oligolectic bee optimization, we used data from Fowler (2020)^149^ to identify which of our predicted interactions were between oligolectic bees and plants in their known host genera.

#### Scenario 3: Restoration to support declining butterflies

This scenario identified plants that best support declining southwestern butterfly species through both larval host plant and adult nectar support. We collated all butterflies classified as “Declining” in the Southwest region from ref.^79^ and used the genetic algorithm to identify plant sets maximizing support for this subset of butterfly species, with the constraint that host and nectar support must come from different plant species (i.e., a butterfly was only considered supported if the plant combination included at least one host plant and at least one distinct nectar plant for that species). Among plant combinations supporting equal numbers of declining butterflies, we selected the set that additionally supported the greatest overall pollinator diversity across all interaction types.

### Quantifying spatial biases and exploring network structure

We summarised the proportion of observed interactions present in the collated interaction data belonging to different genera of plants and pollinators and compared this to our predicted interaction networks. We visualized the degree of each genus in the raw and predicted interaction networks for California native species by aggregating all unique plant-pollinator pairs that occur anywhere (for the raw data) or in any region (for predicted interaction networks), summarising the number of unique pollinator species to plant species interactions between all combinations of plant and pollinator genera, and then plotting them on a graph with the graph visualization software Gephi^84^.

Further, we showed how species and interaction turnover contribute to differences in network structure using the predicted interaction networks for each Jepson district in California. We created maps of expected plant and pollinator diversity across space by summing the relative habitat suitabilities from the ensemble species distribution models (Supplementary Material 2 Figs. S24-25). We also correlated predicted degree of each species in flower visitation and Lepidopteran host user networks to their range size, number of occurrences, and number of raw interactions (Supplementary Material 2 Figs. S26-29). We calculated and plotted nestedness, connectance, and mean degree for pollinators for the predicted regional flower visitation and Lepidopteran host use networks for each Jepson district (Supplementary Material 2 Figs. S30-35). Lastly, we calculated mean interaction network dissimilarity for each pairwise combination of predicted regional interaction networks and partitioned the differences into species turnover and interaction rewiring components (Supplementary Material 2 Figs. S36-41)^166,167^.

*The code for NECTAR and all analyses presented in this manuscript can be found at* github.com/EntoDataSciCornell/NECTAR*, alongside the major data products produced at* doi.org/10.5281/zenodo.22214329*. We used R (v4.2.3)*^168^ *for most analyses. For data handling and manipulation we used tidyverse (v2.0.0)*^169^*, and data.table*^170^ *(v1.16.2). For spatial data handling we used sf (v1.0.21)*^171^*, terra (v1.8.70)*^172^*, raster (v3.6.32)*^173^*, tigris (v2.2.1)*^127^*, maps (v3.4.3)*^174^*, rmapshaper (v0.5.0)*^175^*, and nngeo (v0.4.8)*^176^*. For occurrence data download and access we used rgbif (v3.8.3)*^177^*, ridigbio (v0.4.1)*^178^*, rvest (v1.0.4)*^179^*, and httr (v1.4.7)*^180^*. For occurrence data cleaning we used CoordinateCleaner (v3.0.1)*^181^*, bdc (v1.1.5)*^126^*, and BeeBDC (v1.3.1)*^108^*. For species distribution modelling we used flexsdm (v1.3.9)*^134^*, biomod2 (v4.2.6.2)*^182^*, ecospat (v4.1.2)*^183^*, blockCV (v3.2.0)*^184^*, CovSel (v0.1.0)*^185^*, dismo (v1.3.16)*^186^*, adehabitatHR (v0.4.22)*^187^*, mda (v0.5.5)*^188^*, earth (v5.3.4)*^189^*, geodata (v0.6.6)*^190^*, rnaturalearth (v1.1.0)*^191^*, nhdplusTools (v1.3.2)*^192^*, and FedData (v4.3.0)*^193^*. For phenology modelling we used circular (v0.5.2)*^194^ *and boot (v1.3.28.1)*^195^*. For interaction data scraping and preparation we used rvest (v1.0.4)*^179^ *and rotl (v3.1.0)*^196^*. For phylogenetic data handling we used ape (v5.8.1)*^197^*. For interaction prediction and network analysis we used bipartite (v2.23.0)*^198^ *and nngeo (v0.4.8)*^176^*. For the genetic algorithm plant selection analyses we used futureverse (v1.34.0)*^199^*. For Bayesian statistical modelling and analysis we used brms (v2.23.0)*^163^*, cmdstanr (v0.9.0)*^200^*, tidybayes(v3.0.7)*^201^ *. For visualization we also used patchwork (v1.3.0)*^202^*, cowplot (v1.1.3)*^203^*, ggdist (v3.3.3)*^204^*, ggalluvial (v0.12.5)*^205^*, tidyterra (v0.7.2)*^206^*, and units (v0.8.5)*^207^*. For reading external data files we used readxl (v1.4.3)*^208^*. For the graph embedding and transfer learning, we implemented them in Julia (v1.10.10)*^119^ *using packages CSV.jl (v0.10.15)*^209^*, DataFrames.jl (v1.7.0)*^210^*, Distributions.jl (v0.25.120)*^211^*, JLD2.jl (v0.6.2)*^212^*, PhyloNetworks.jl (v0.16.4)*^157^*, Pipe.jl (v1.3.0)*^213^*, Random.jl (a core standard library in Julia)*^119^*, SpeciesInteractionNetworks.jl (v0.0.1)*^156^*, and StatsBase.jl (v0.34.6)*^214^.

## Supplementary

*Supplementary Material 1* - Checklist of native plants and pollinators that could potentially have been included in NECTAR, including the 5,130 plant and 5,131 pollinator species that we include in NECTAR’s interaction prediction (species which we were able to run SDMs and phenometric estimations for).

*Supplementary Material 2* - Additional figures and tables

*Supplementary Material 3* - Optimized plant lists for each restoration scenario presented and the corresponding pollinator support scores.

## Acknowledgements

We want to thank Matt Forister, Matt Pennell, Russel Dinnage, Lora Morandin (Pollinator Partnership), Jennifer Hopwood, Mace Vaughan (Xerces Society), and Jim Strittholt for thoughtful feedback throughout the project. We also thank Cynthia Powell and the rest of the CalFlora team for providing us with the context and input on native plant data. We also thank Steve Nanz from the Moth Photographers Group for providing key Lepidoptera occurrence data. This work resulted from the Morpho ‘Plant Prioritizer’ Working Group conducted at the National Center for Ecological Analysis and Synthesis at the University of California, Santa Barbara. We also acknowledge funding from the California Department of Food and Agriculture (Grant Number: 23-0743-000-SG) for supporting the development of the Calscape Pollinator Companion. LMG, TB, YYJC were supported by Cornell University, CTC was supported by Conservation Biology Institute.

