## SupplementaryMaterial2 for "Local interaction networks reconstructed from global biodiversity data improve pollinator restoration decision making"

### Supplementary

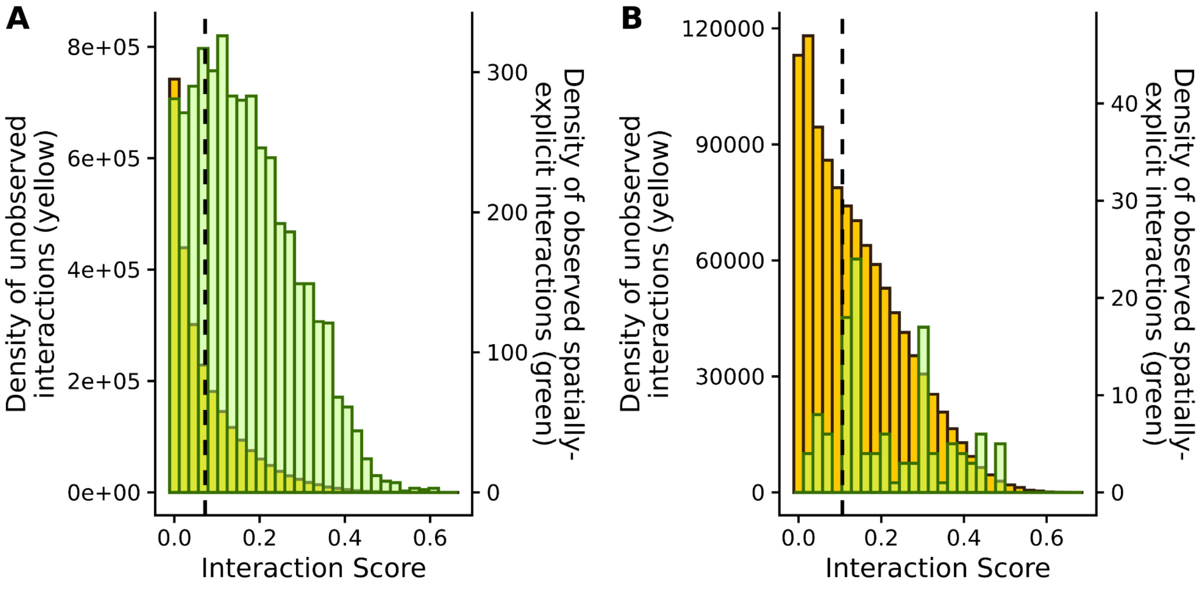


**Figure S1**. Distribution of calculated interaction plausibility scores for observed spatially-explicit plant-pollinator interaction pairs in California (green) and unobserved interaction pairs (yellow) among potential plant-pollinator pairs identified using the phylogenetic constraint (pollinator species may potentially interact with any plant in genera they have previously been observed to interact with), separately for flower visitation interactions (A) and Lepidopteran host plant interactions (B). Additionally, we visualize the 25th percentile values used to categorize unobserved interactions as plausible vs. implausible (dashed line).

**Data summary**

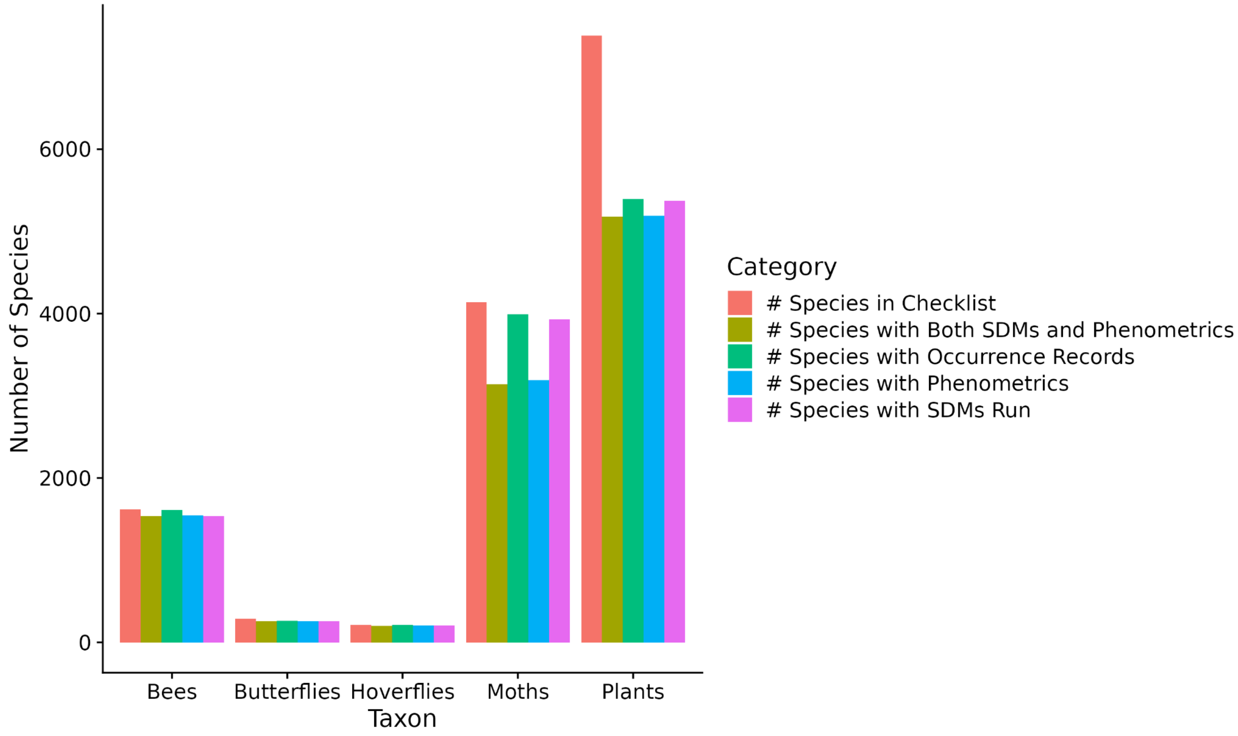


**Figure S2**. Number of species in checklist by taxon group (red), alongside the number of species that also had at least one occurrence records (green), that we estimated their spatial distribution for (purple), that we estimated phenometrics for (blue), and that had both SDMs and phenometrics estimated for (yellow).

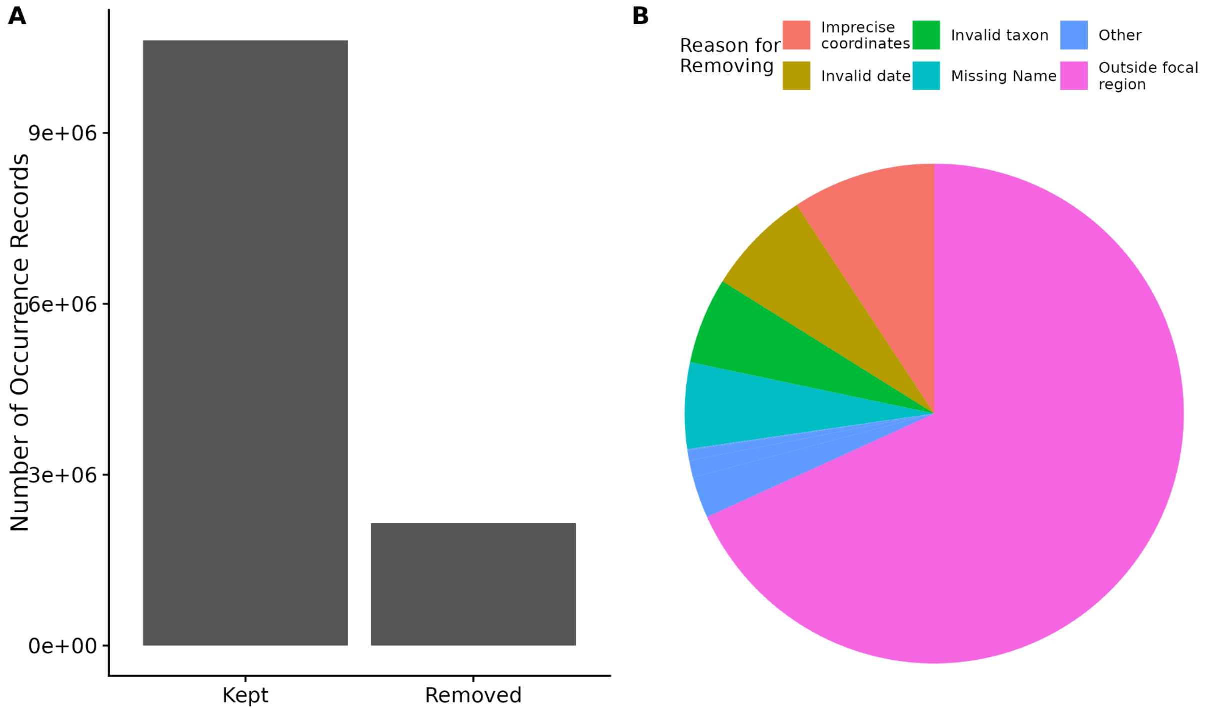


**Figure S3**. Summary of occurrence record cleaning across all taxon groups. **A.** The number of records removed, relative to the number remaining, following all occurrence cleaning steps. **B.** Pie chart showing the cause of removal for occurrence records. The majority of records removed were due to being outside the focal region (California plus a spatial buffer; pink) or other coordinate issues (red).

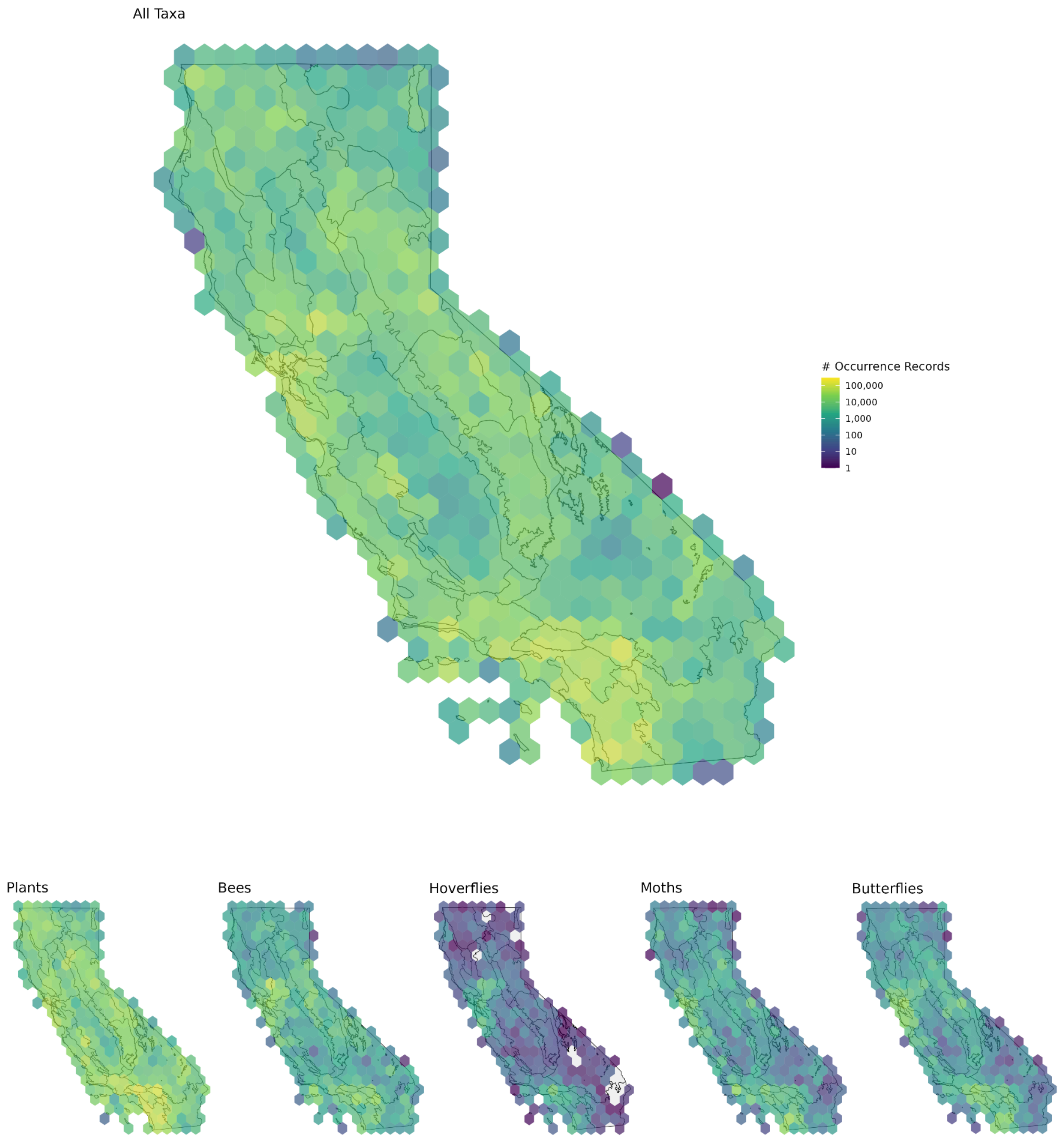


**Figure S4**. Maps of the spatial distribution of all occurrence records within the boundaries of California across all taxonomic groups (top), and broken up by taxon group (bottom). The legend in all cases has been normalized for comparability across taxonomic groups.

**Species Distribution Models**

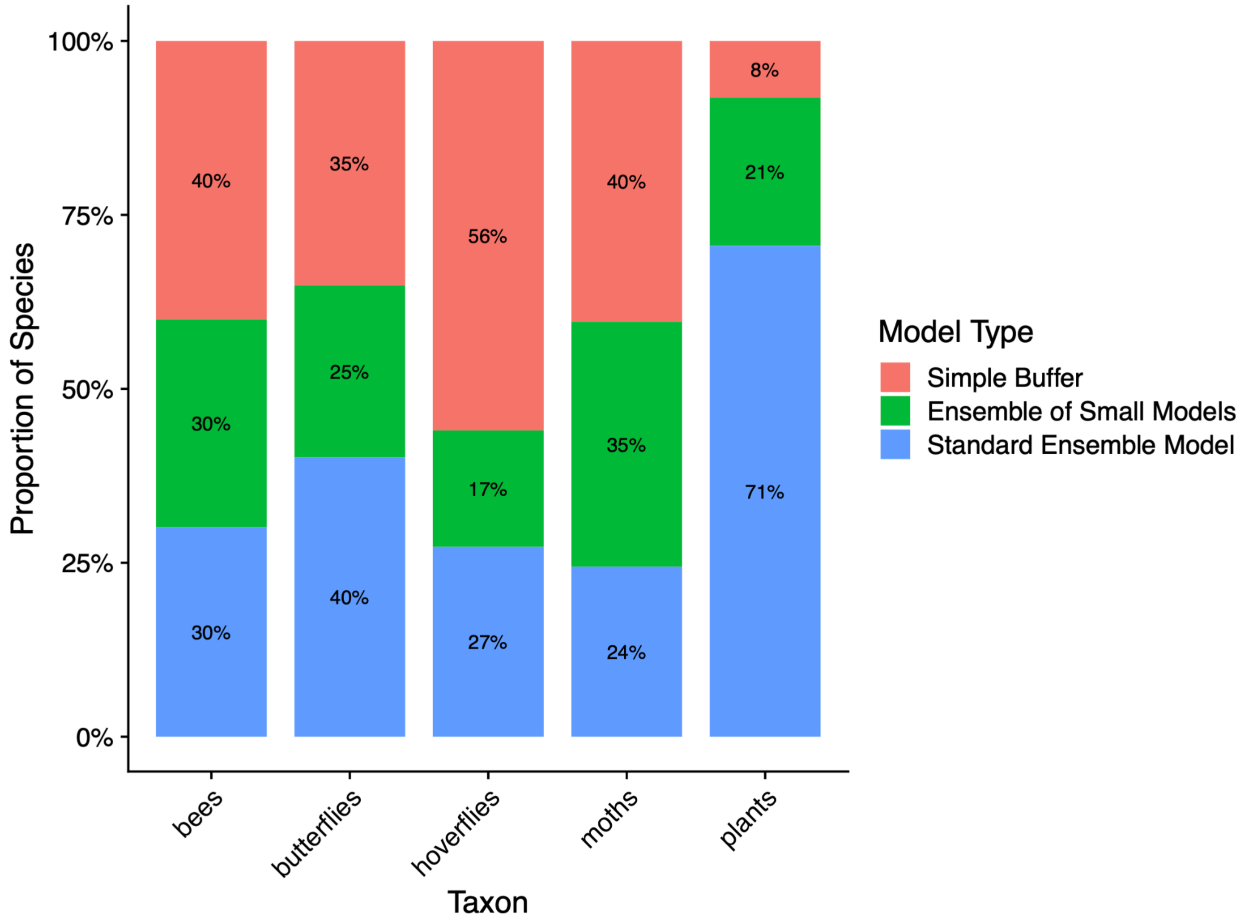


**Figure S5**. Breakdown of the proportion of species in each taxonomic group that were modelled with standard Ensemble Species Distribution Models (blue), Ensemble of Small Models (green), or had their spatial distribution estimated using a simple buffer approach (red).

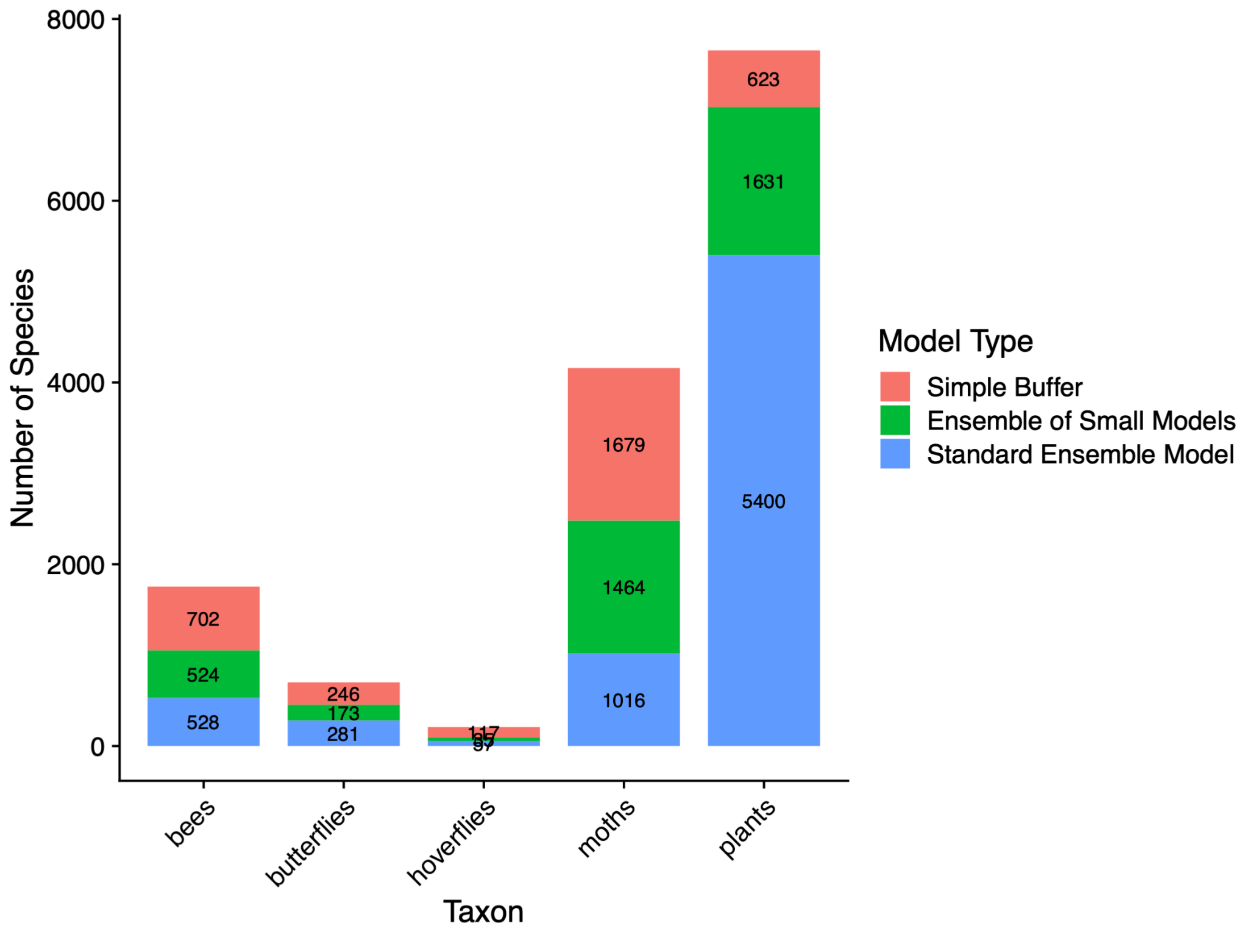


**Figure S6**. The total number of species in each taxonomic group that were modelled with standard Ensemble Species Distribution Models (blue), Ensemble of Small Models (green), or had their spatial distribution estimated using a simple buffer approach (red).

**Phenometrics**

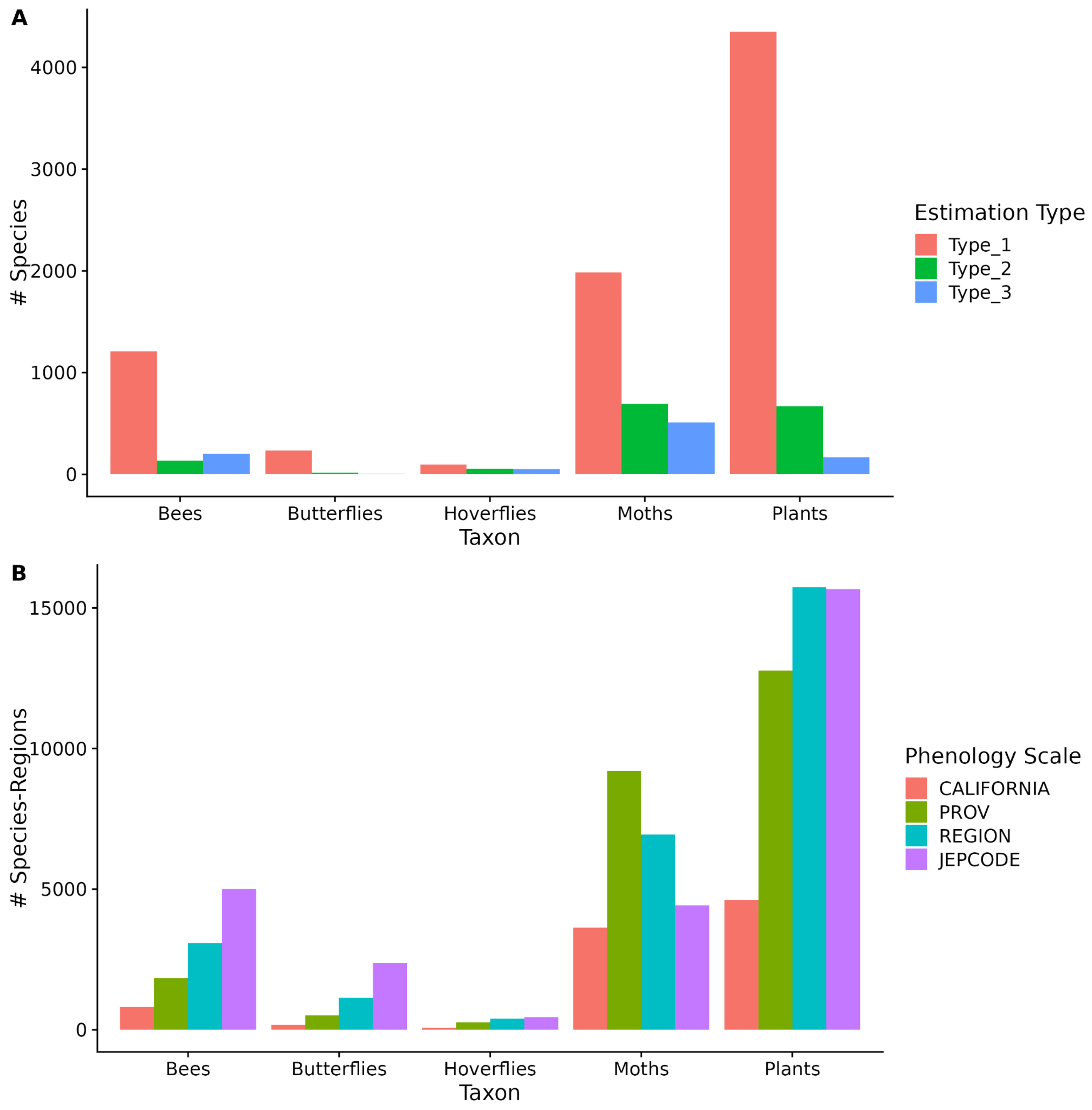


**Figure S7**. Summary of method and scale used to estimate species phenometrics. **A.** The relative number of species by taxon group estimated with the Type 1 method (estimating spatially-explicit onset and offset using maximum likelihood fit of skewed Von Mises distribution using dated occurrence records for the appropriate life stage; red), Type 2 method (species-level circular interval approach; green), and Type 3 method (species-level seven day buffer around dated observation; blue) . **B.** The breakdown of number of species-region combinations that were estimated at different spatial scales for species estimated using the Type 1 method. Bees, butterflies, and hoverflies had the most species-regions combinations estimated at the finest spatial resolution used in our analysis (Jepson district regions, i.e., JEPCODES), with Jepson regions without adequate data pooling from successively coarser scales.  Moths had the largest proportion of data-poor species, requiring them to pool more information from coarser spatial scales to estimate phenometrics compared to other taxonomic groups.

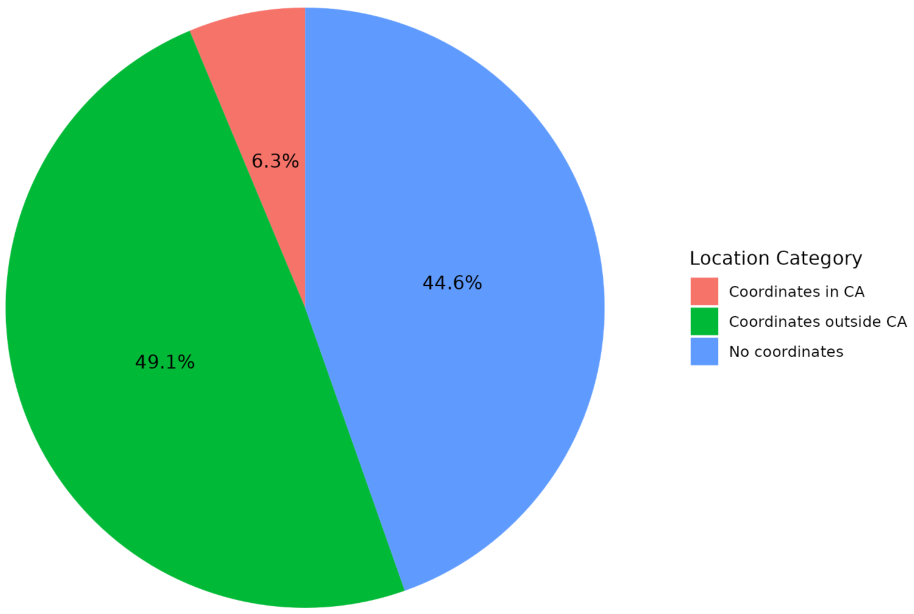


**Figure S8**. Pie chart showing % of interaction data used containing coordinates in California (red), outside of California (green), and without detailed spatial information (blue).

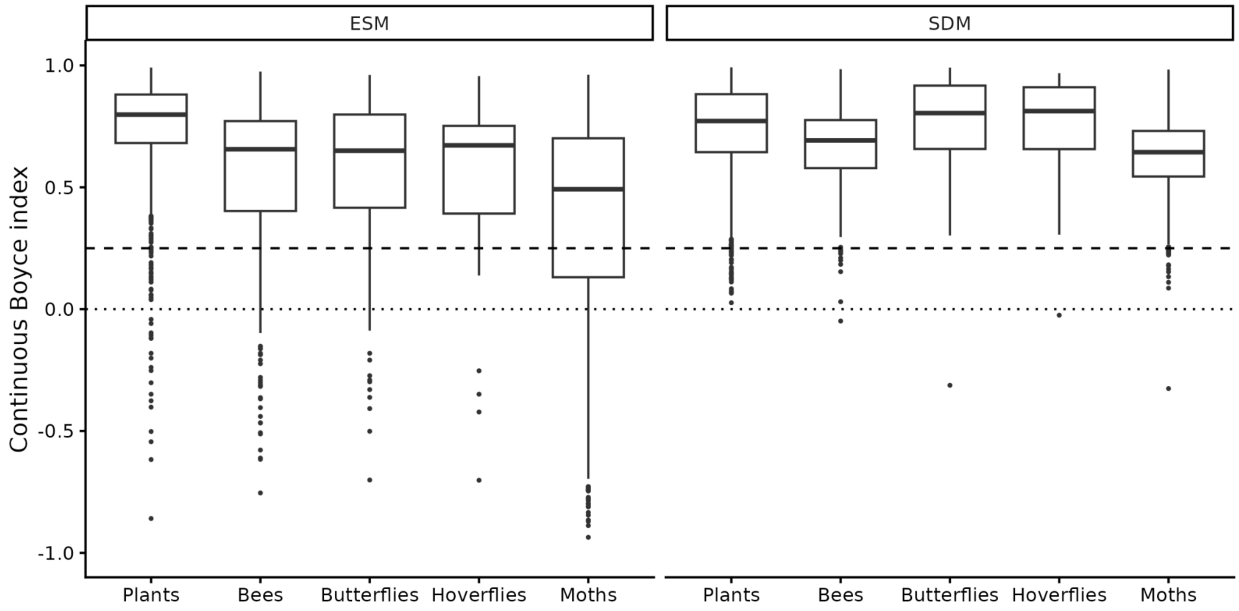


**Figure S9.** Distribution of spatial cross-validation performance across plant and insect species distribution models. Boxplots show species-level continuous Boyce index values for ensembles of small models (ESM) and full ensemble models (SDM) across taxonomic groups. Horizontal lines indicate random predictive performance (CBI = 0) and the conservative reference benchmark CBI = 0.25.

#### Interaction Data and Predictions

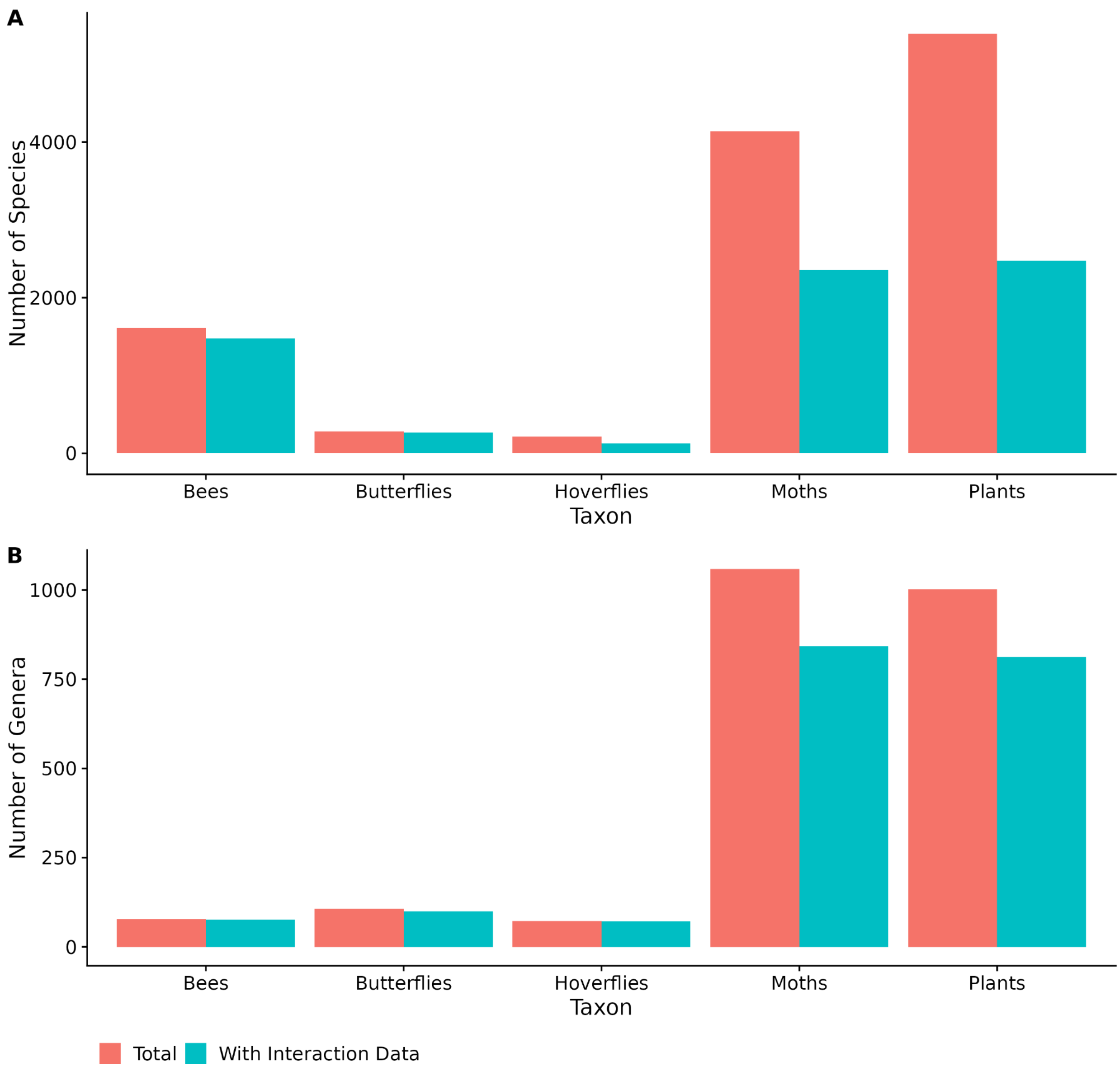


**Figure S10.** Number of species (A) and genera (B) in our analysis (red) and with at least one interaction record (blue), broken up by taxon group. Moths and plants have both the largest number of species and genera, and have the lowest representation of interaction data across both of these taxonomic scales.

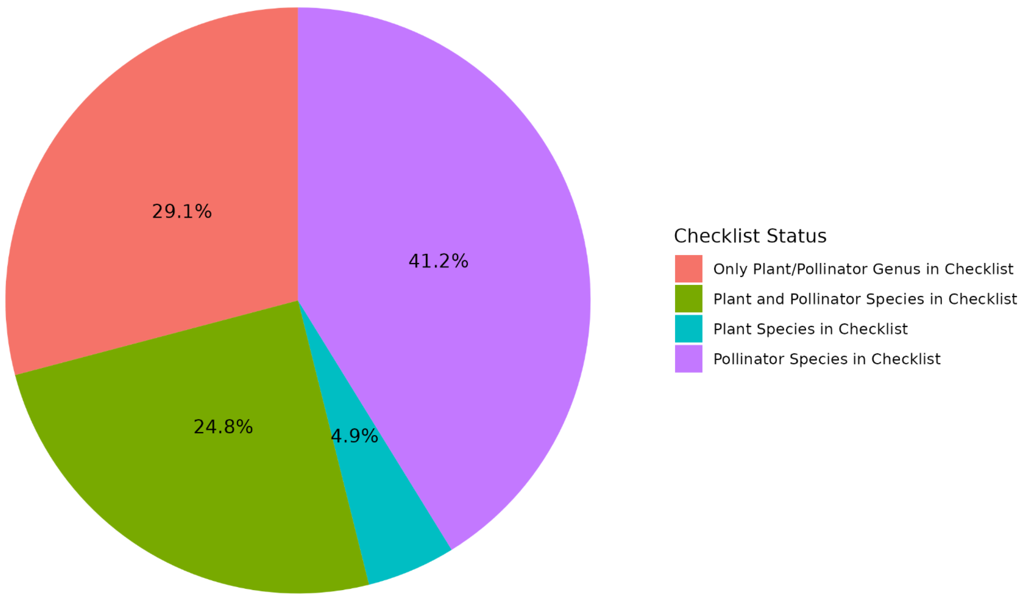


**Figure S11**. Pie chart showing the taxonomic scale of interaction data used in our analysis. This plot shows among all interaction records used, only ~¼ of records were specific to both plant and pollinator species occurring in California (green). We also leverage interaction data where only the pollinator species is native to California (purple), only the plant species is native to California (blue), or where only the nominal plant and pollinator genus is native to California (red). In all cases, the absence of species information could also cause that record to be classified only at the genus level.

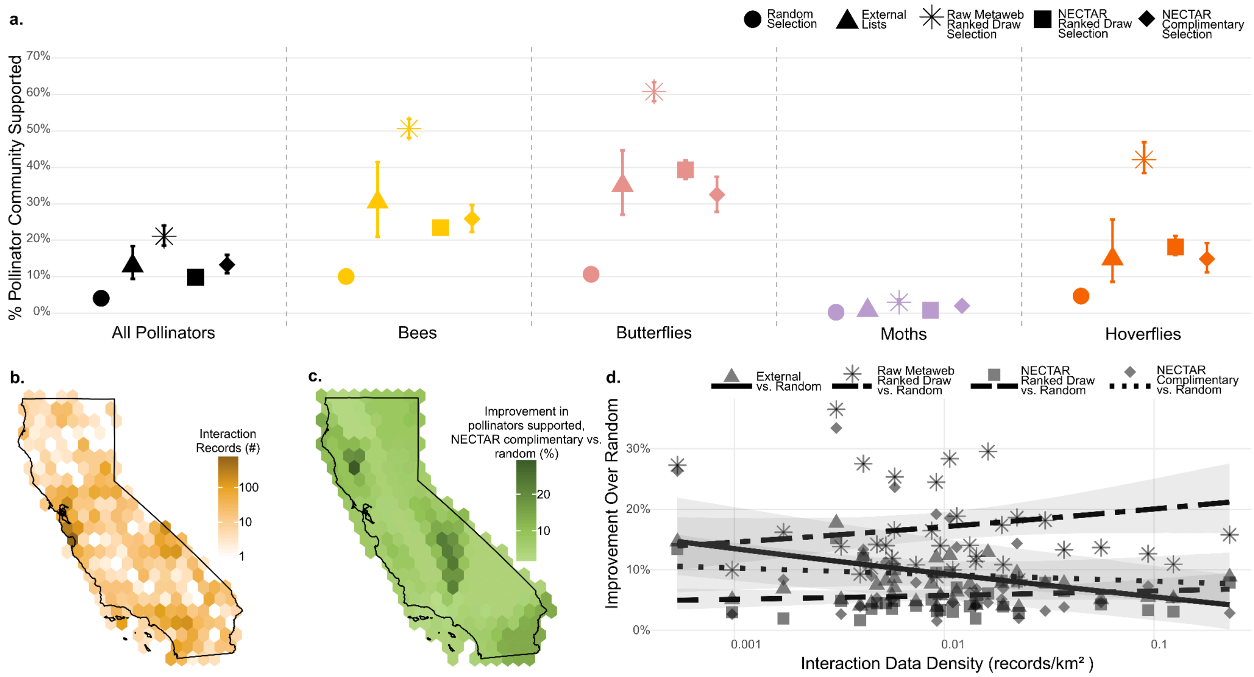


**Figure S12**. Identical to Figure 2, except that here the proportion of the pollinator community potentially supported by the plant mixes in each selection approach is based on a single metaweb using only previously observed species-level interactions without predicted interactions from NECTAR. **a.** The proportion of the pollinator community expected to be supported by 10 plants selected increased as we shifted from: (i) random selection from the pool of regional native plants (circle), but it stayed approximately consistent between (ii) random draws from external expert-opinion based plant lists (triangles), (iii) random draws from the top ranked pollinator plants we identify using a metaweb of the collated raw interaction data (stars), (iv) random draws from the top ranked pollinator support plants we identify with NECTAR (squares), and (v) the optimal set of complimentary pollinator support plants identified with NECTAR (diamonds). When prioritizing plants for different taxonomic groups we found the same pattern, but with differing levels of support by random plants, where moths are generally the least supported while butterflies are the most supported. Error bars show 95% credible intervals. **b.** The density map of spatially-explicit plant-pollinator interaction data across California was higher around urban centers such as San Francisco and Los Angeles (also see Supplementary

Material 2 Fig. S12). **c.** The expected absolute increase in total pollinator richness supported by an optimized set of 10 plants compared to a random selection of plants across California was greater around areas with less interaction data, especially the Southern Sierra Nevada and Mendocino National Forest. **d.** Trends of proportion of pollinator community supported across the gradient of interaction data density, showing that plant recommendations from NECTAR (squares from the random draws of top degree plants and diamonds from the optimized selection) and expert-opinion lists were comparable and consistent across data density, while plant recommendations derived from a metaweb of the raw interaction data (stars) improved in expected pollinator support with respect to interaction data density. Points correspond to a comparison within one of the 35 ecoregions in our analysis, and lines show the marginal mean relationship between interaction density and increase in pollinator community support. Colors in this figure mirror color for taxonomic groups in all figures below. Ribbons represent 95% credible intervals.

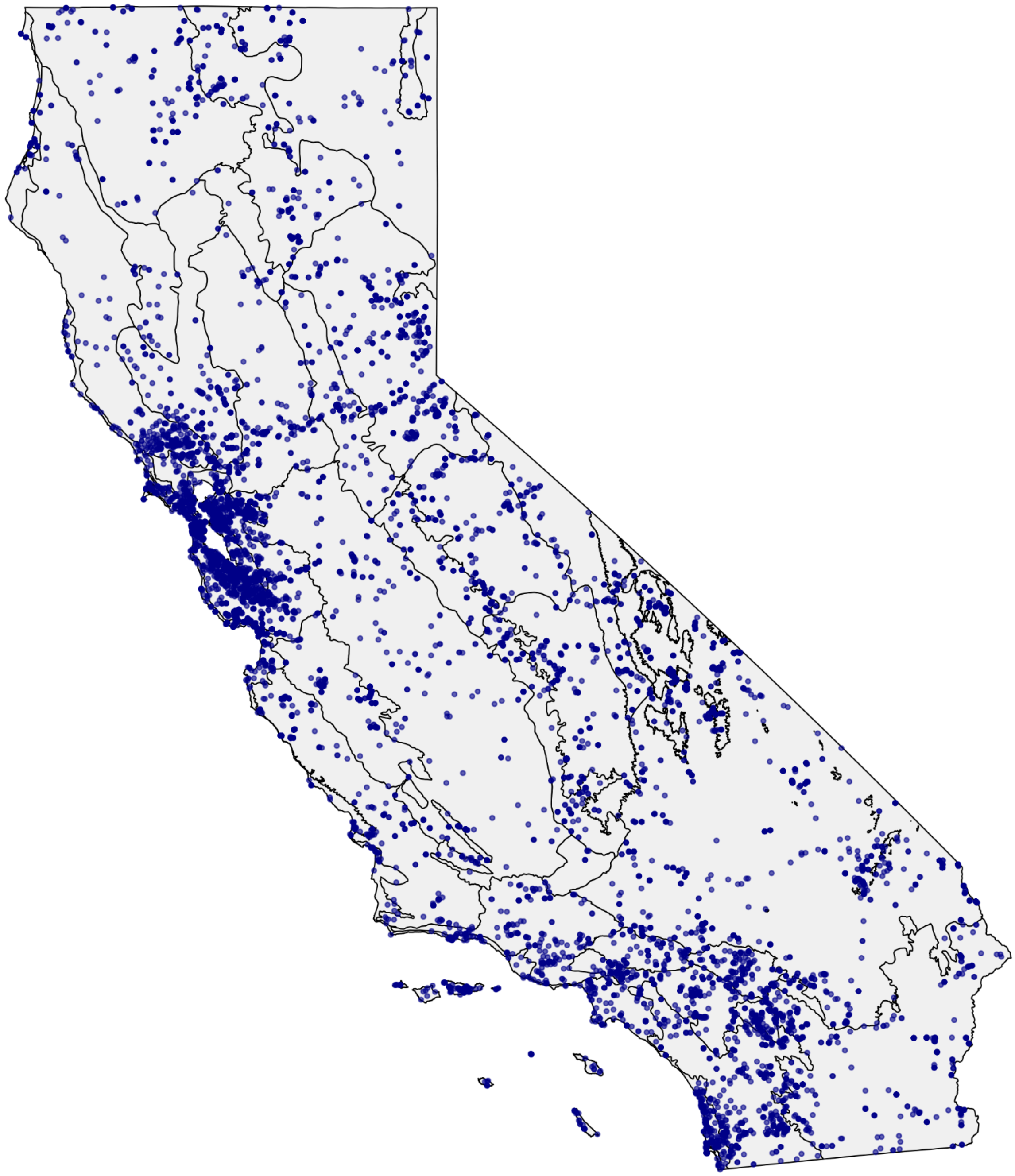

**Figure S13**. Map of locations of spatially-explicit interaction records within the boundaries of California, shown on top of the 35 Jepson region boundaries used in our analysis.

**Validation**

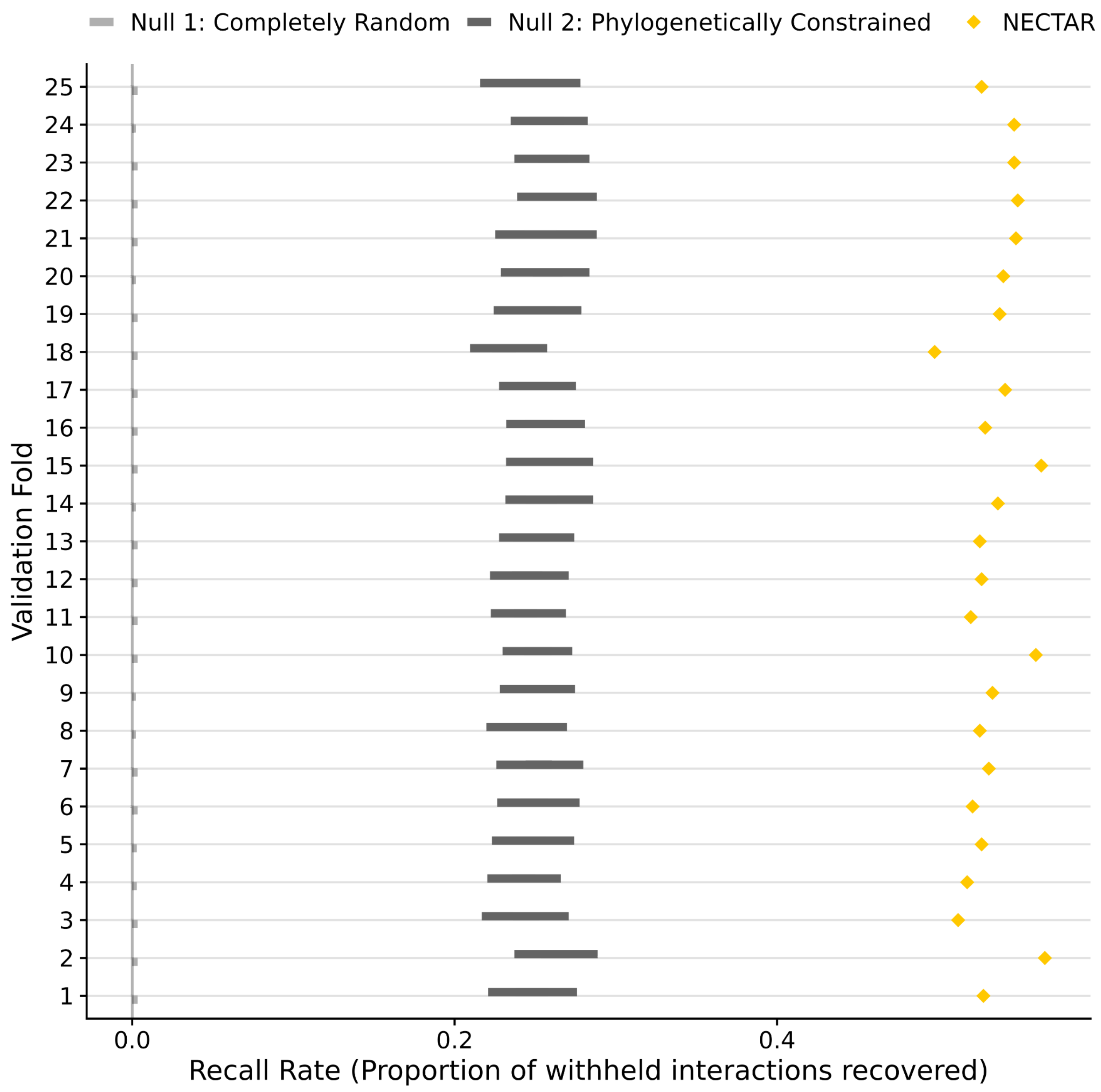


**Figure S14**. Distributions of recall rate under two distinct null models and NECTAR (yellow) for 25 validation iterations that omitted 20% of spatially-explicit interactions between native California plants and pollinators with locality within California. Null model 1 (grey) used an entirely random approach to select the same number of interactions predicted by NECTAR from the entire set of plant-pollinator combinations that have historically occurred in a given ecoregion. Null model 2 (black) randomly selects the same number of interactions predicted by NECTAR from the constrained set of interaction pairs that NECTAR calculates interaction plausibility scores for. Mean pseudo-precision score of spatially-explicit withheld interactions was 2.33 across validation iterations (range: [2.23, 2.43]), where the candidate species pair was constrained to plant-pollinator pairs identified using the phylogenetic constraint (pollinator species can potentially interact with congeners of plants they have previously been observed on) in ecoregions where both species in candidate pairs have historically been observed.

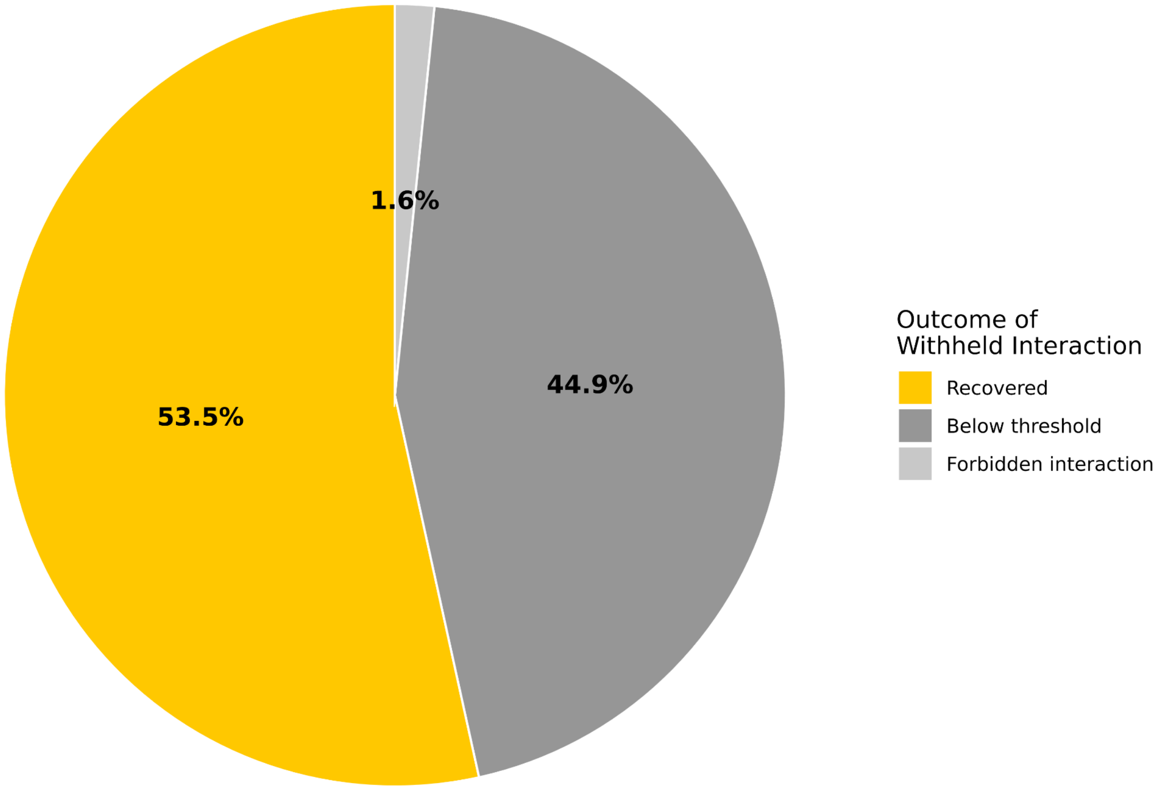


**Figure S15**. Breakdown of the outcome of withheld flower visitation interactions using NECTAR across all validation iterations. On average, NECTAR recovered 53.8% of interactions withheld during validation (yellow), and the vast majority of interactions not recovered were because they fell below the interaction plausibility score threshold (44.6%; dark grey), while only 1.6% of interactions were not recovered because those links became forbidden from the phylogenetic constraint (light grey).

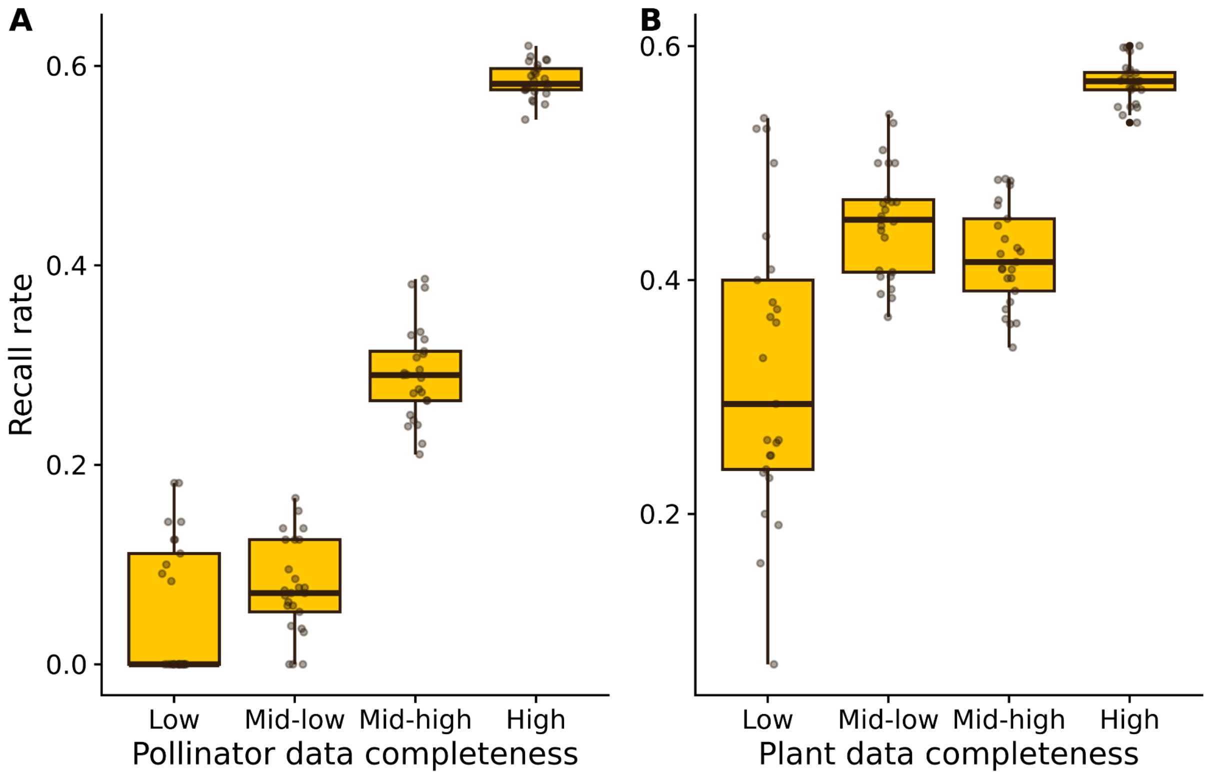


**Figure S16.** Recall rate distributions across data-completeness quartiles for pollinator (A) and plant (B) taxa from the leave-out interaction prediction validation. Completeness quartiles were defined from the total number of flower visitation records for each taxon in the full interactions dataset (not restricted to spatially-explicit California records). Each point represents the recall rate calculated within a single validation run for that completeness bin.

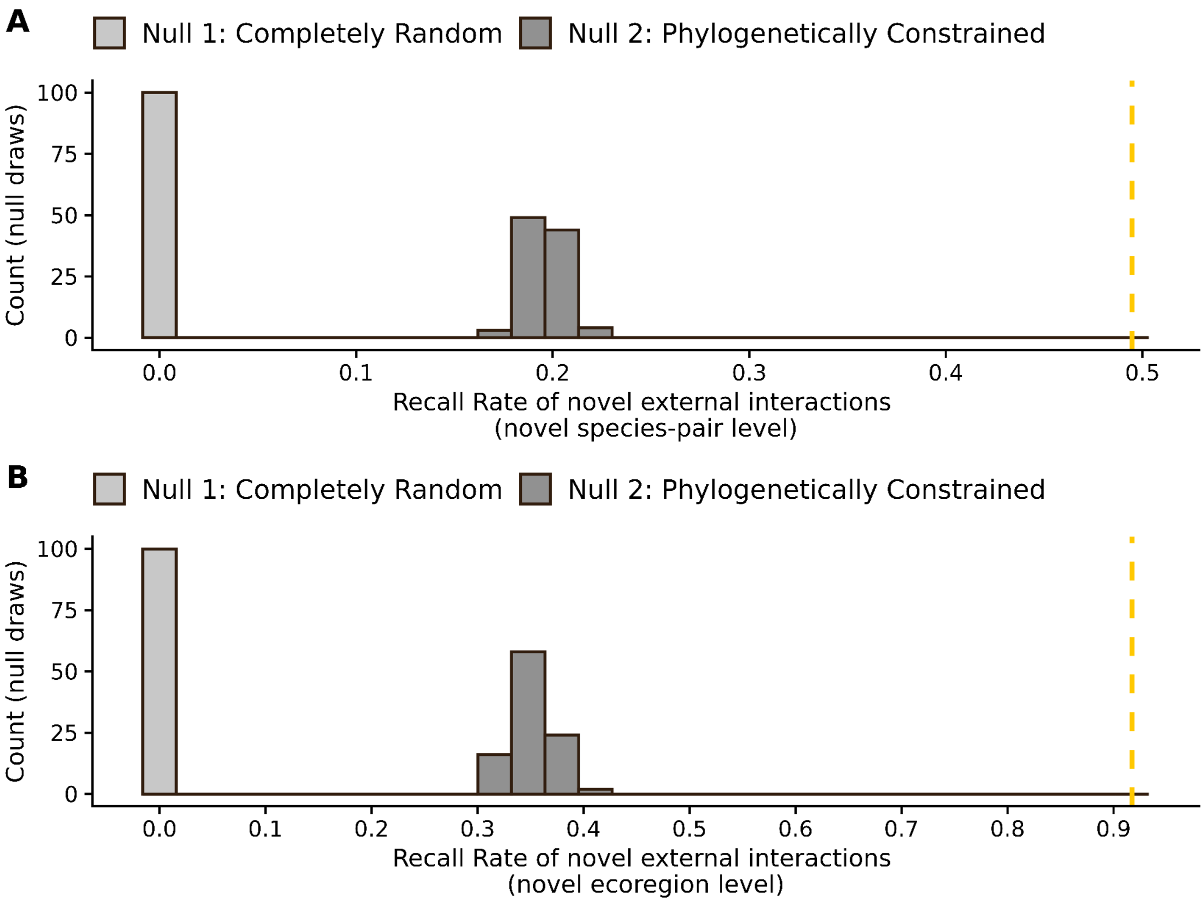


**Figure S17.** Recall rates (dashed yellow line) of novel spatially-explicit interactions from external datasets not used in our predictive pipeline whereby the plant-pollinator pair had not been recorded in the interaction data we collated at all (A) or for that specific ecoregion (B). The histograms show the distribution of recall rates under two null models. NECTAR recalled 49.5% of previously unobserved interaction pairs, which was greater than both null models (A; Null Model 1: mean = 0.1%, range = [0.0%, 0.4%]; Null Model 2: mean = 19.6%, range = [17.4%, 22.2%]); and 91.7% of interactions that were previously observed in other regions, which was greater than both null models (B; Null Model 1: mean = 0.1%, range = [0.0%, 0.6%]; Null Model 2: mean = 35.2%, range = [31.6%, 39.8%]). The computed pseudo-precision score was 4.18 for entirely novel interactions and 4.68 for previously observed interactions novel to a new ecoregion. Here, the candidate species pair was constrained to plant-pollinator pairs identified using the phylogenetic constraint (pollinator species can potentially interact with congeners of plants they have previously been observed on) and with graph embedding plus transfer learning (for pollinator species with no prior interaction data) in ecoregions where both species in candidate pairs have historically been observed.

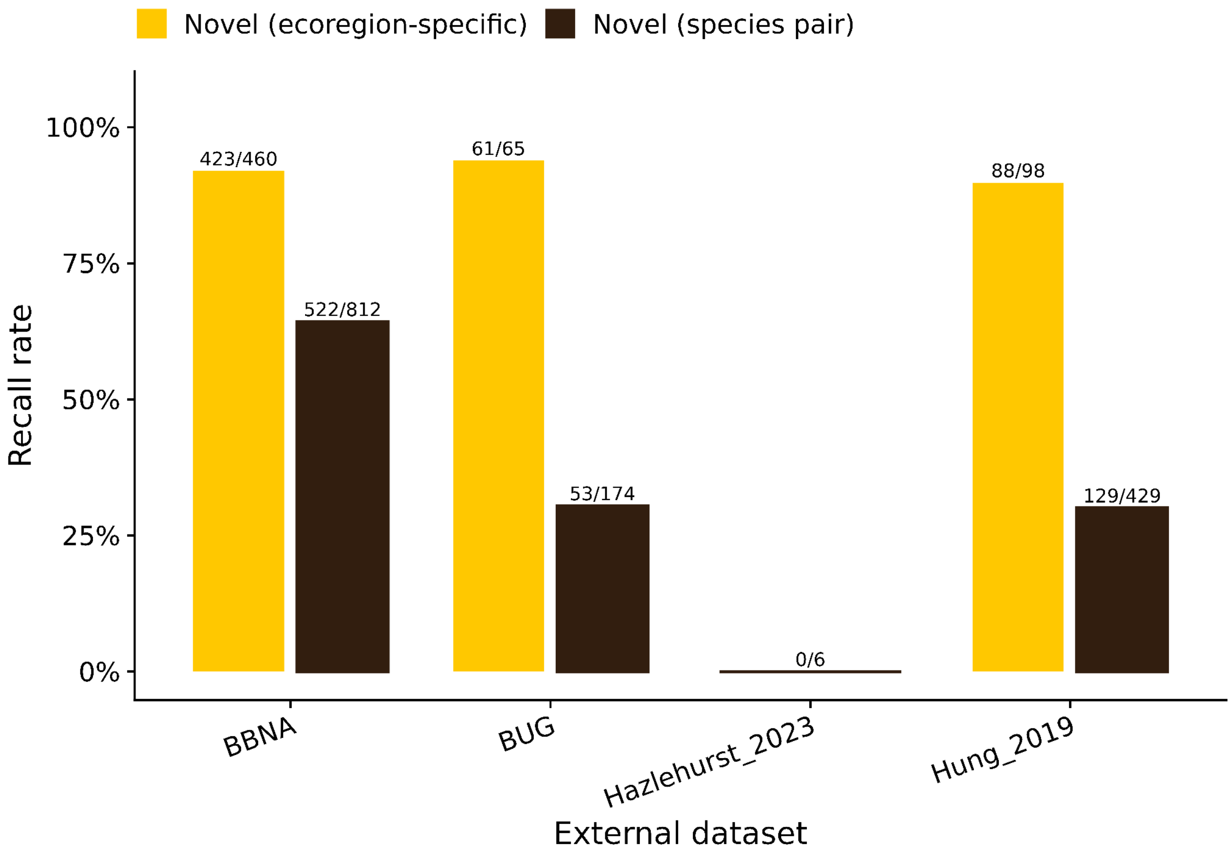


**Figure S18.** Recall rates of novel spatially-explicit interactions broken up by external dataset. For interaction pairs that have been previously observed previously in other regions, NECTAR exhibited the highest recall rates for the Biodiversity in Urban Gardens (BUG) community science project led by California Native Plant Society, followed closely by the Bumblebees of North America dataset (BBNA), which was the most geographically complete of the four assessed, Hung et al.'s (2019) plant-pollinator surveys of San Diego remnant coastal sage scrub habitats, and lastly by Hazlehurst et al.'s (2023) pollinator surveys of five rare plants of San Clemente Island, which included only six interactions among plants and pollinators included in our analysis. We observed the same recall pattern for interaction pairs which have not previously been observed anywhere, with the exception that novel interactions observed in BBNA had a higher recall rate than from those observed in BUG. Across datasets, most unique interactions (52.2% in BBNA, 69.6% in BUG, 100% in Hazlehurst et al., and 76.7% in Hung et al.) were not previously recorded in any of the primary datasets collated for this study. Further, across datasets, most observations which had been previously recorded were new to the ecoregion (43.9% in BBNA, 78.3% in BUG, and 75.4% in Hung et al.).

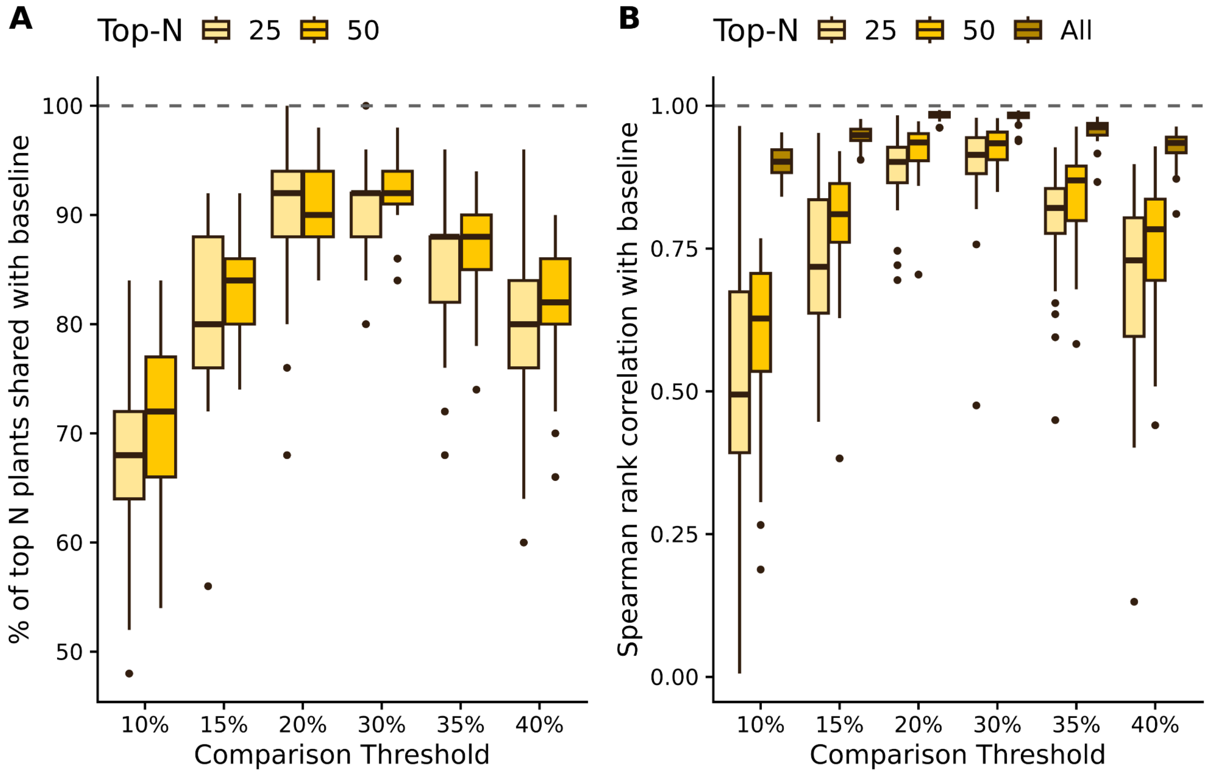


**Figure S19.** Sensitivity of ecoregional plant rankings to the interactionScore threshold used to define supported plant–pollinator interactions. In all cases, various thresholds were compared against the 25% baseline threshold derived from the interaction score distribution of observed spatially explicit interactions (Figure S1). **(A)** Percent of baseline top-25 (yellow) and top-50 (orange) plants per ecoregion that are also present in the corresponding list at each alternative threshold. **(B)** Spearman rank correlation between baseline and alternative-threshold plant rankings per ecoregion, calculated among the top 25 (yellow), top 50 (orange), and all ranked plants (dark gold).

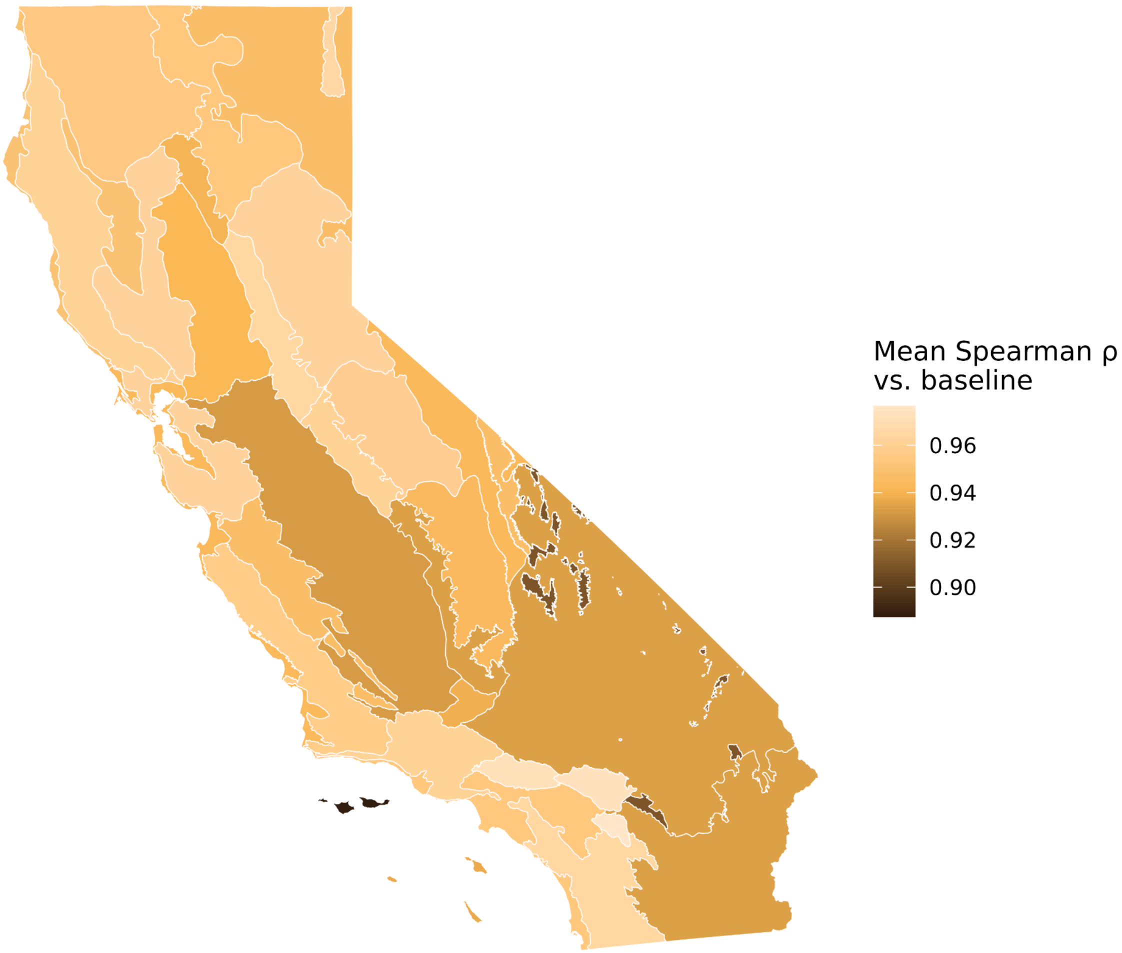


**Figure S20.** Average Spearman rank correlation of ecoregional plant pollinator-support rankings between the 25% baseline threshold (derived from the interaction score distribution of observed spatially-explicit interactions; Figure S1) and each of the other thresholds tested in the threshold sensitivity analysis (10%, 15%, 20%, 30%, 35%, and 40%). Plant rankings are most sensitive to threshold choice in the Desert Mountains and North Channel Islands ecoregions, and to a lesser extent in the Mojave and Sonoran Deserts and San Joaquin Valley regions.

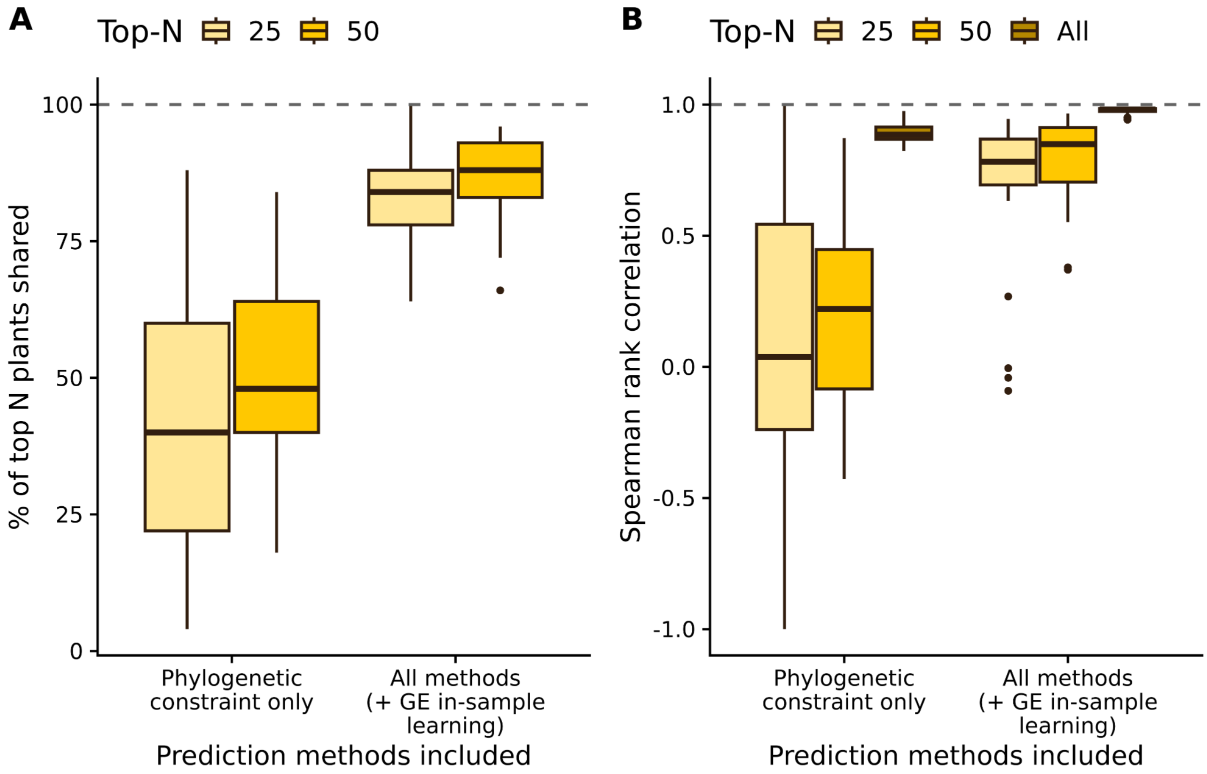


**Figure S21.** Sensitivity of ecoregional plant rankings to the method(s) used to define potential interaction links. Specifically, we compared the default (phylogenetic constraint – where we assume a pollinator could potentially interact with any plant in a genus it has previously been observed to visit – for pollinators with existing data and plant genera identified from graph embedding with transfer learning only for pollinators with no existing data) to only the phylogenetic constraint (thus omitting all pollinators with no pre-existing interaction data) or to the default plus additional potential links identified from graph embedding for species with existing interaction data. **(A)** Percent of baseline top-25 (yellow) and top-50 (orange) plants per ecoregion that are also present in the corresponding list in each scenario. **(B)** Spearman rank correlation between baseline and alternative-assumption scenarios' plant rankings per ecoregion, calculated among the top 25 (yellow), top 50 (orange), and all ranked plants (dark gold).

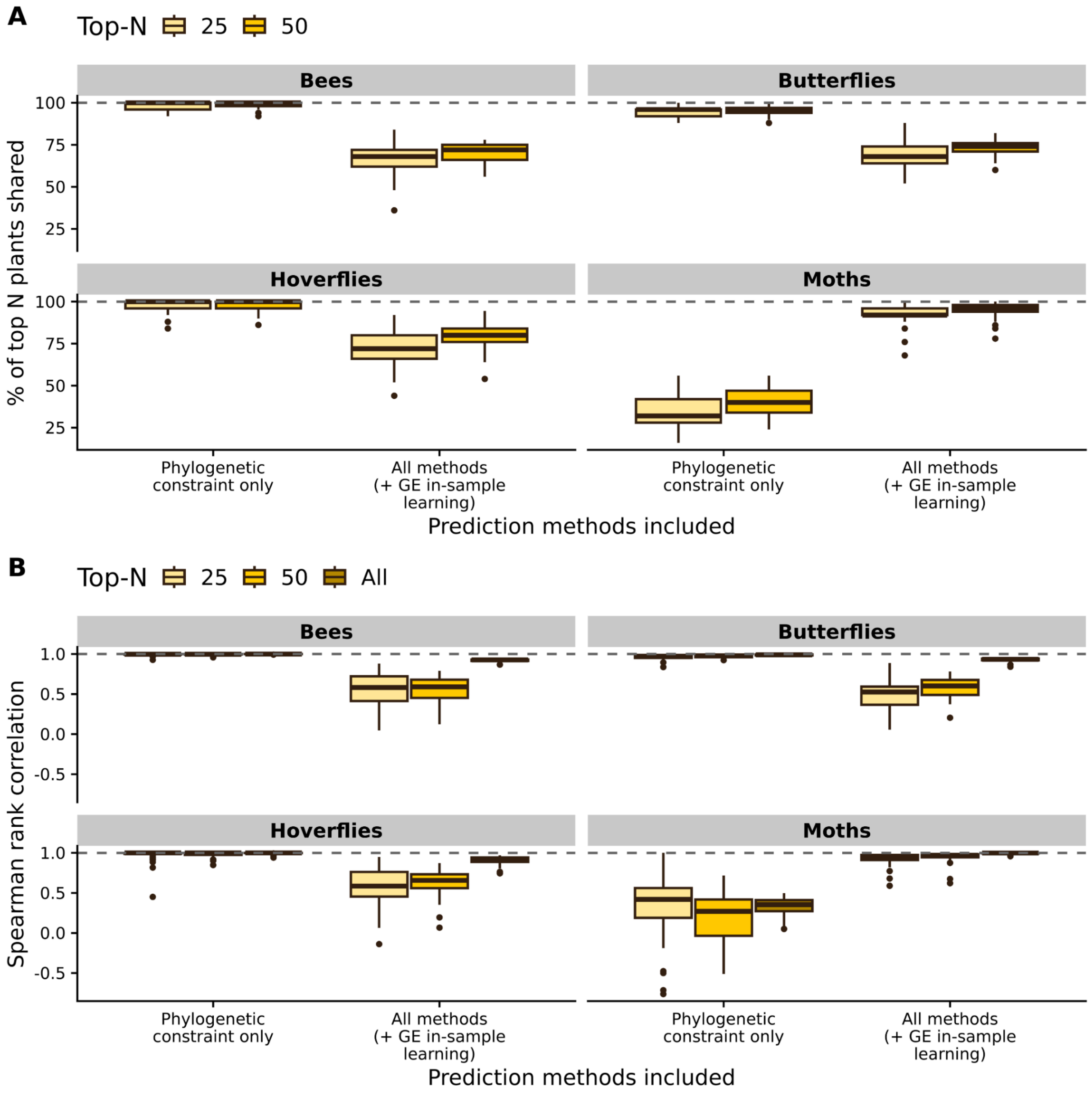


**Figure S22.** Sensitivity of ecoregional plant rankings, specific to each target pollinator taxon group, to the method(s) used to define potential interaction links. Specifically, we compared the default (phylogenetic constraint – where we assume a pollinator could potentially interact with any plant in a genus it has previously been observed to visit – for pollinators with existing data and plant genera identified from graph embedding with transfer learning only for pollinators with no existing data) to only the phylogenetic constraint (thus omitting all pollinators with no pre-existing interaction data) or to the default plus additional potential links identified from graph embedding for species with existing interaction data. Here, the top plant lists are identified based on predicted interactions for each focal taxonomic group separately (i.e., optimizing for bees separately from butterflies, etc.). **(A)** Percent of baseline top-25 (yellow) and top-50 (orange) plants per ecoregion that are also present in the corresponding list in each scenario. **(B)** Spearman rank correlation between baseline and alternative-assumption scenarios' plant rankings per ecoregion, calculated among the top 25 (yellow), top 50 (orange), and all ranked plants (dark gold).

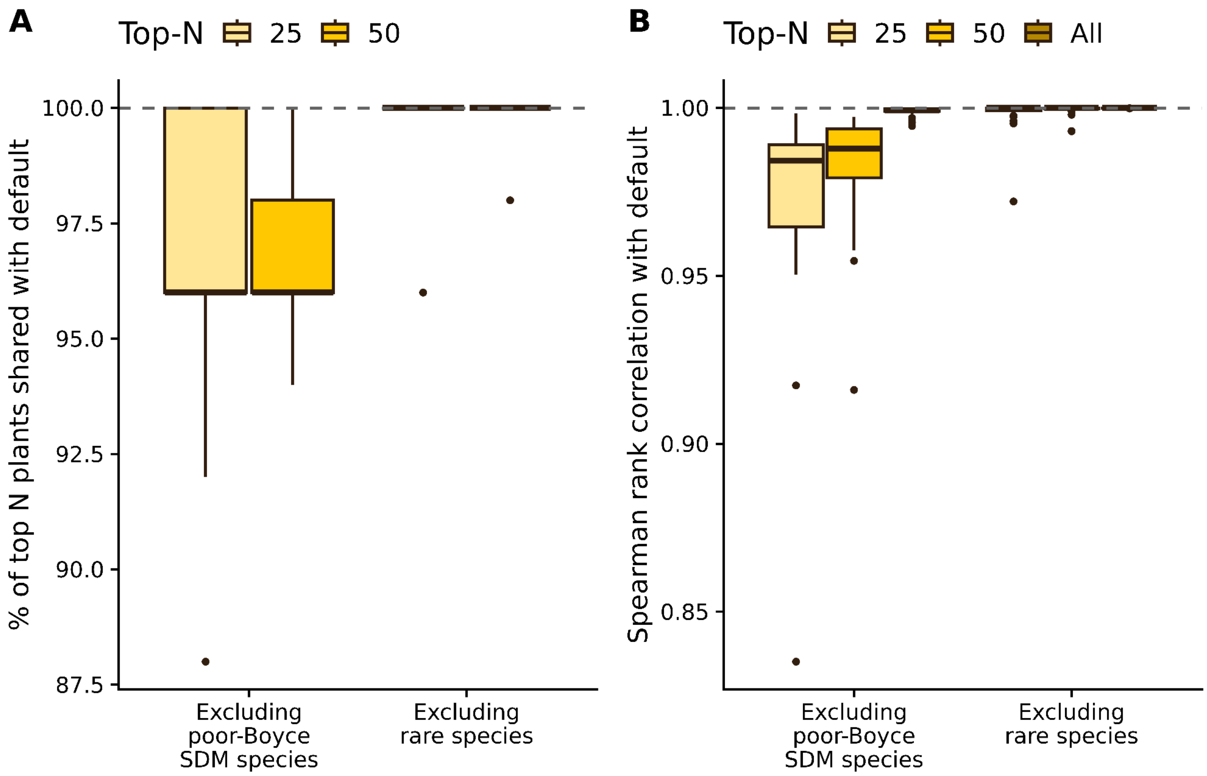


**Figure S23.** Sensitivity of the ecoregional plant rankings to the exclusion of species whose SDMs had poor fit (continuous Boyce index <= 0.25) or very rare taxa (less than 7 unique grid cells with an occurrence used for the SDMs). **(A)** Percent of baseline top-25 (yellow) and top-50 (orange) plants per ecoregion that are also present in the corresponding list in each scenario. Note that the y-axis starts at approximately 87.5%.**(B)** Spearman rank correlation between baseline and alternative-assumption scenarios' plant rankings per ecoregion, calculated among the top 25 (yellow), top 50 (orange), and all ranked plants (dark gold). Note that the y-axis starts at approximately 0.825.

**Diversity**

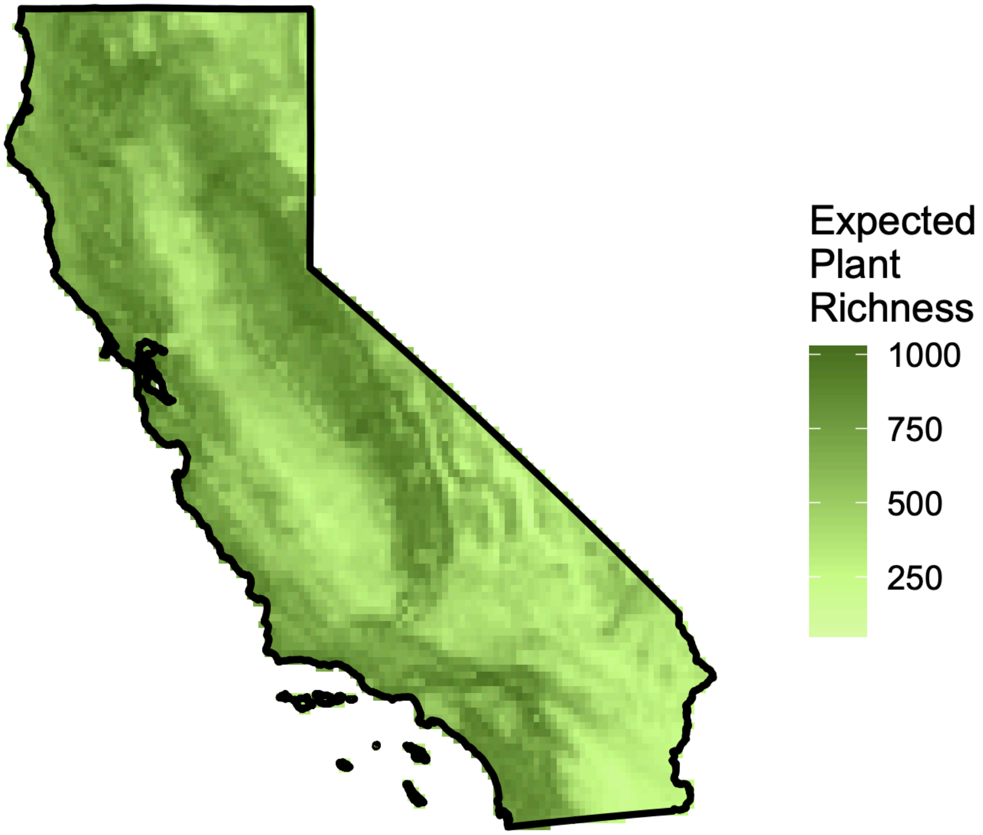


**Figure S24.** Expected richness of the 5,178 plants included in our analysis across space. Expected richness was calculated here as the sum of the relative habitat suitabilities from the Ensemble Species Distribution Models.

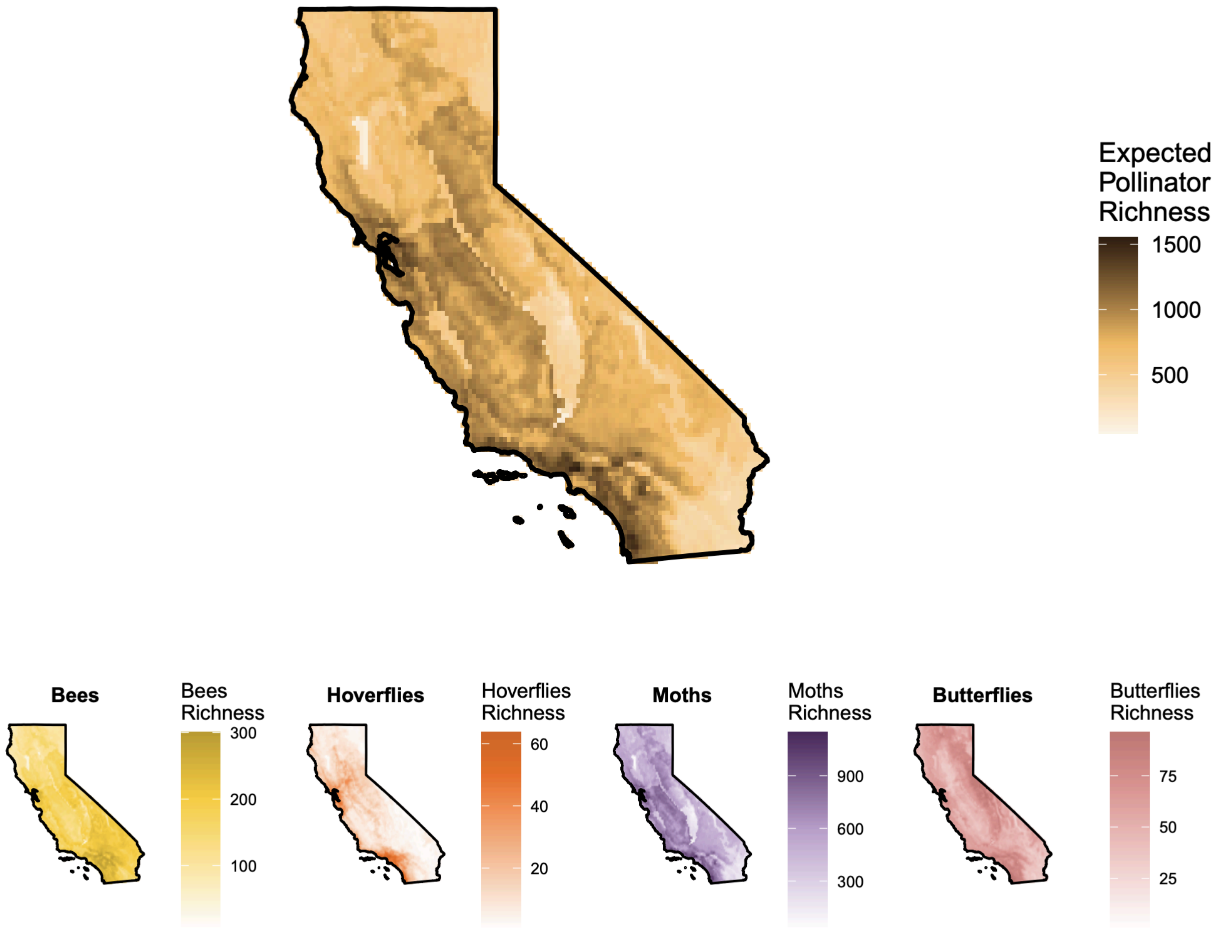


**Figure S25.** Expected richness of the 5,131 pollinators included in our analysis across space (top) and broken up by pollinator taxonomy group (bottom). Expected richness was calculated here as the sum of the relative habitat suitabilities from the Ensemble Species Distribution Models.

#### Network Structure

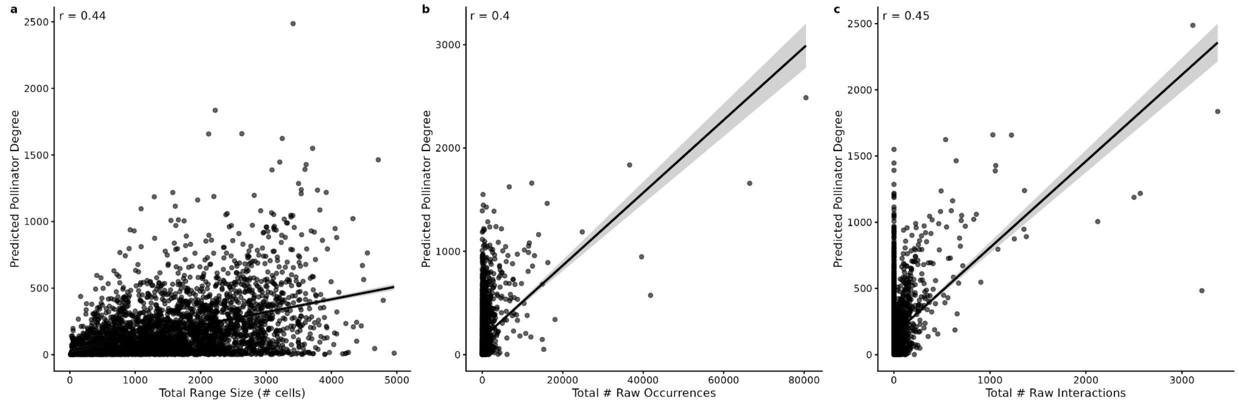


**Figure S26.** Scatterplot of overall pollinator degree across California from our predicted flower visitation interaction networks (y axis) versus range size of pollinator in California (a, determined using the sum of grid cells from the binary SDMs), total number of raw occurrences (b), and total number of raw flower visitation interaction records recorded for that species (c).

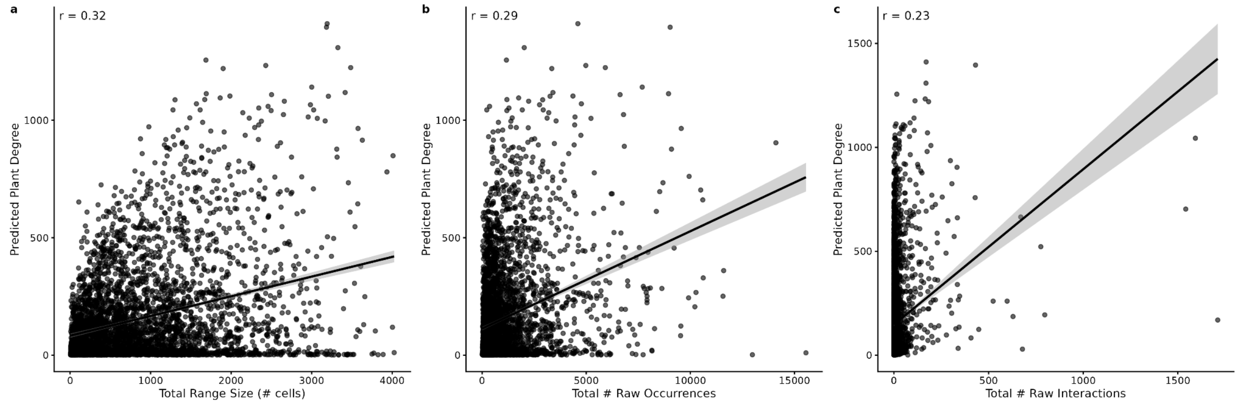


**Figure S27.** Scatterplot of overall plant degree across California from our predicted flower visitation interaction networks (y axis) versus range size of plant in California (a, determined using the sum of grid cells from the binary SDMs), total number of raw occurrences (b), and total number of raw flower visitation interaction records recorded for that species (c).

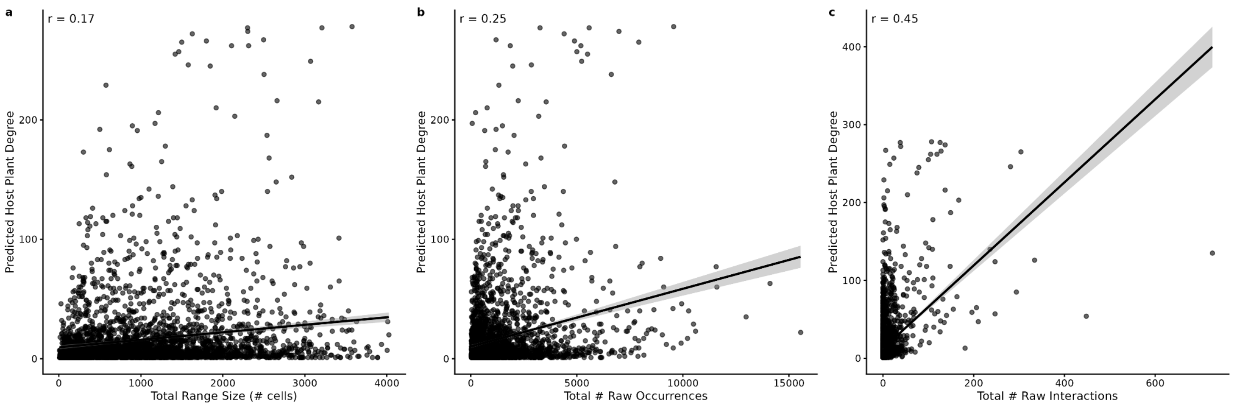


**Figure S28.** Scatterplot of overall pollinator degree across California from our predicted Lepidopteran host use networks (y axis) versus range size of pollinator in California (a, determined using the sum of grid cells from the binary SDMs), total number of raw occurrences (b), and total number of raw host use interaction records recorded for that species (c).

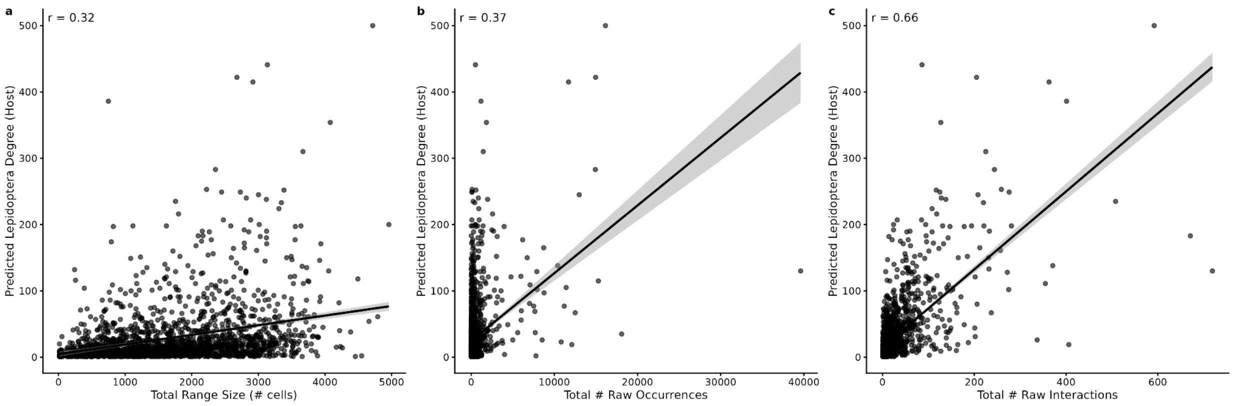


**Figure S29.** Scatterplot of overall plant degree across California from our predicted Lepidopteran host use networks (y axis) versus range size of plant in California (a, determined using the sum of grid cells from the binary SDMs), total number of raw occurrences (b), and total number of raw host use interaction records recorded for that species (c).

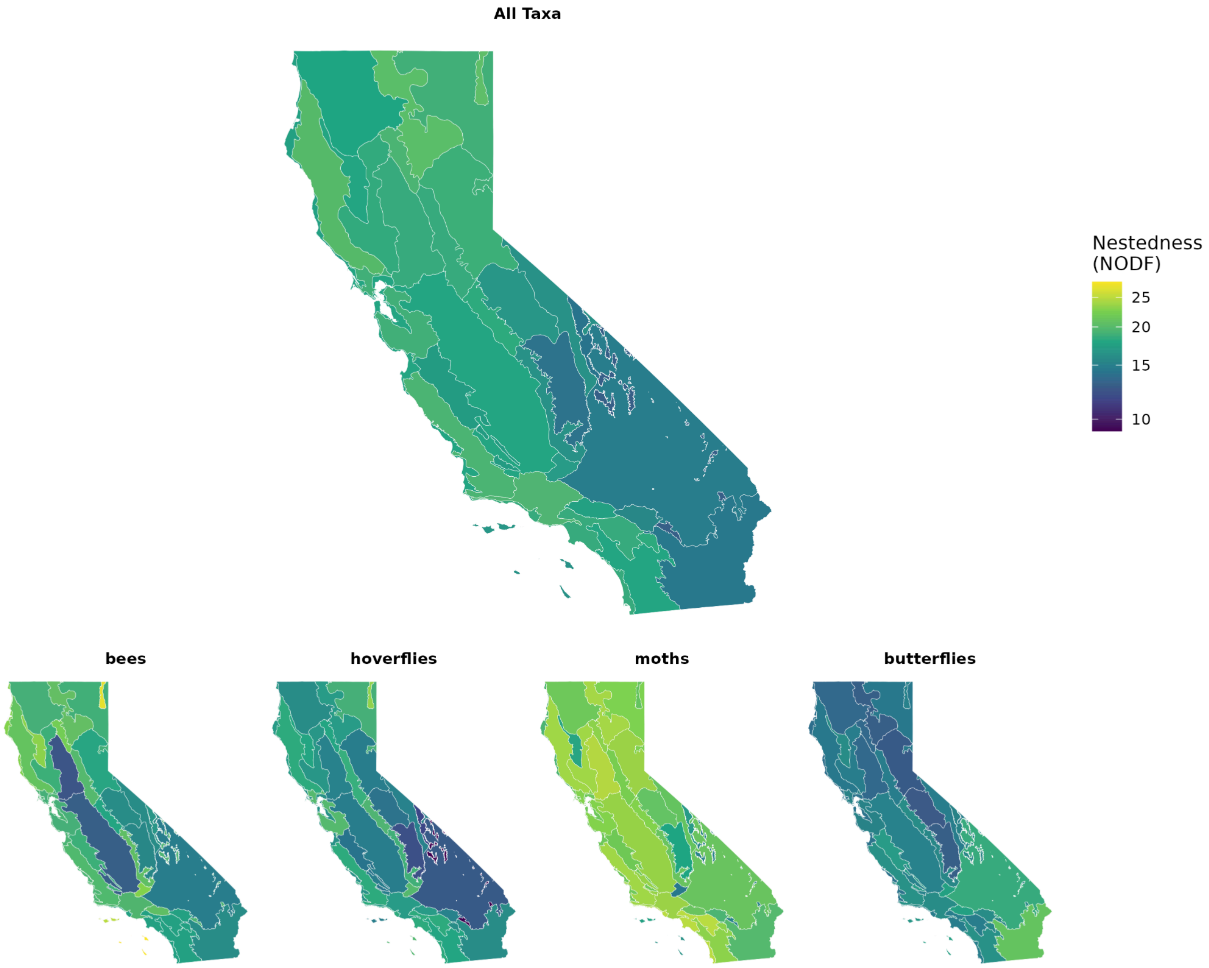


**Figure S30**. Nestedness of our predicted flower visitation interaction networks by ecoregion and pollinator taxonomic group.

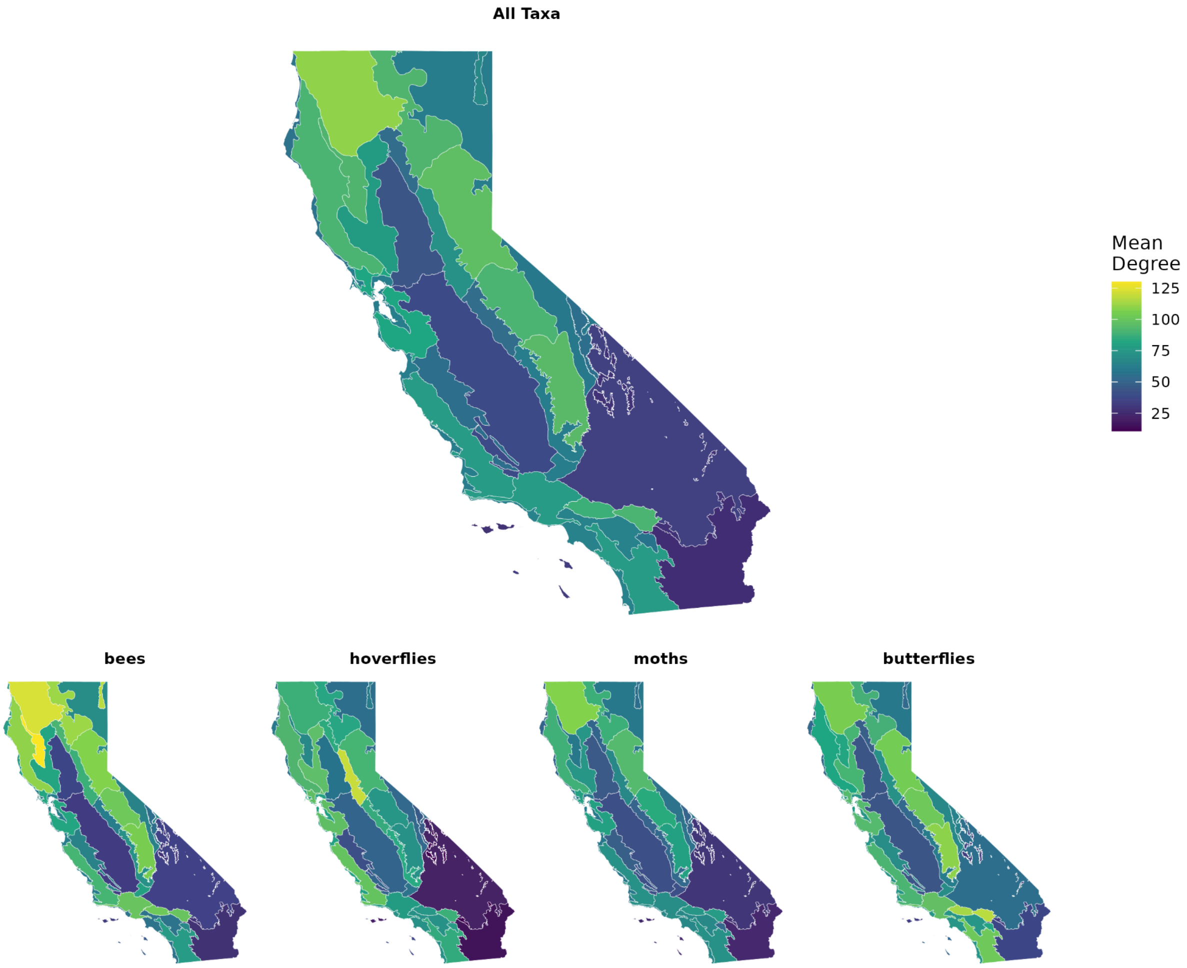


**Figure S31**. Mean number of plant partners (degree) for each ecoregion and taxonomic group from our predicted flower visitation networks.

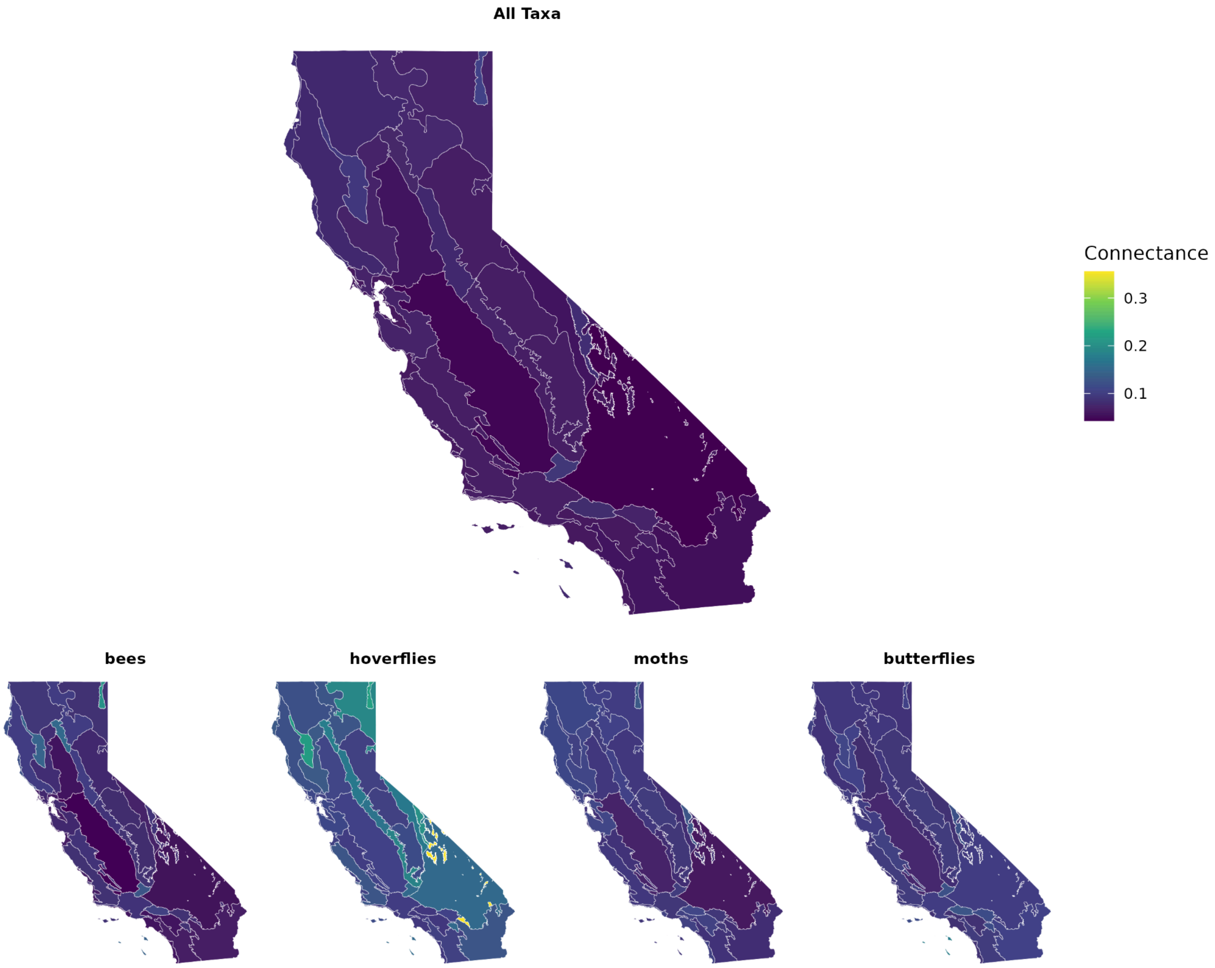


**Figure S32**. Network density of reconstructed bipartite plant-pollinator networks (connectance) for each ecoregion and taxonomic group from our predicted flower visitation networks. The mean connectance across ecoregions of the full interaction network was 0.0696 (min = 0.0428, max = 0.106). When filtering the networks based on each taxonomic group, the mean connectance across ecoregions was 0.0916 (min = 0.0448, max = 0.201) for bee-plant networks; 0.101 (min = 0.0751, max = 0.178) for butterfly-plant networks; 0.144 (min = 0.0953, max = 0.356) for hoverfly-plant networks; and 0.0970 (min = 0.0610, max = 0.134) for moth-plant networks.

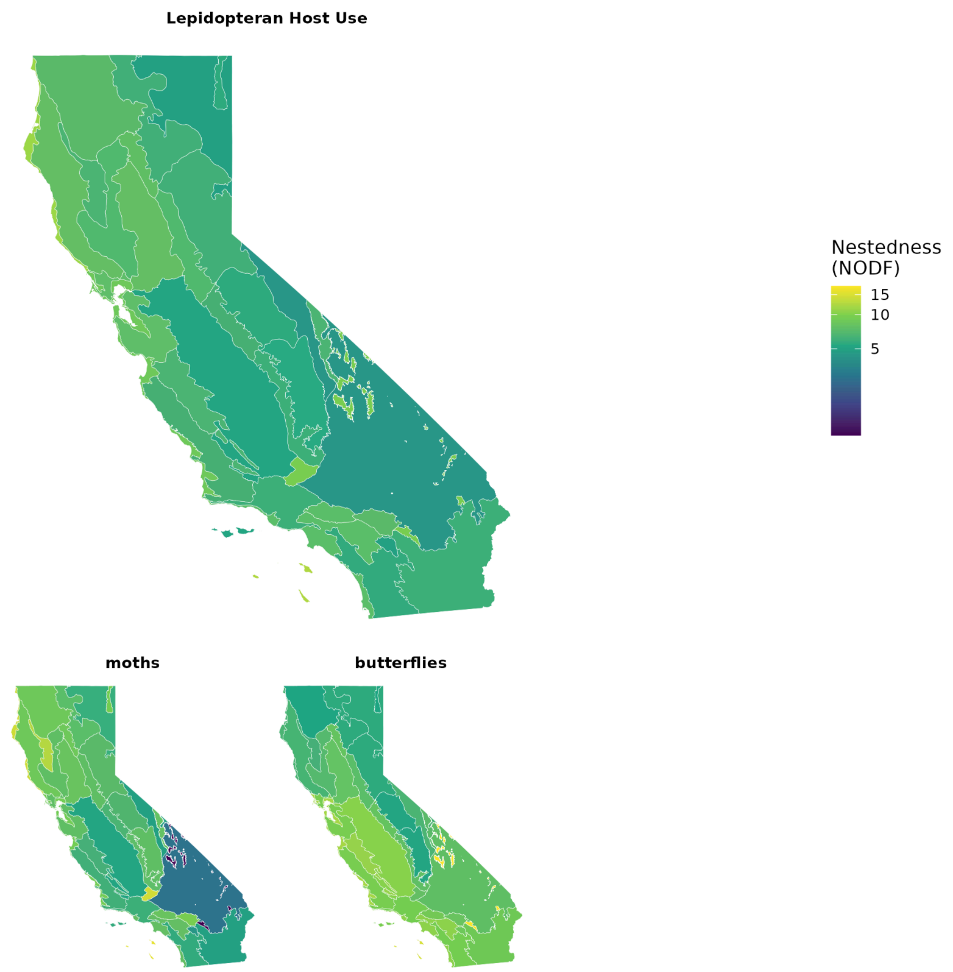


**Figure S33**. Nestedness of our predicted Lepidopteran host use networks by ecoregion and pollinator taxonomic group.

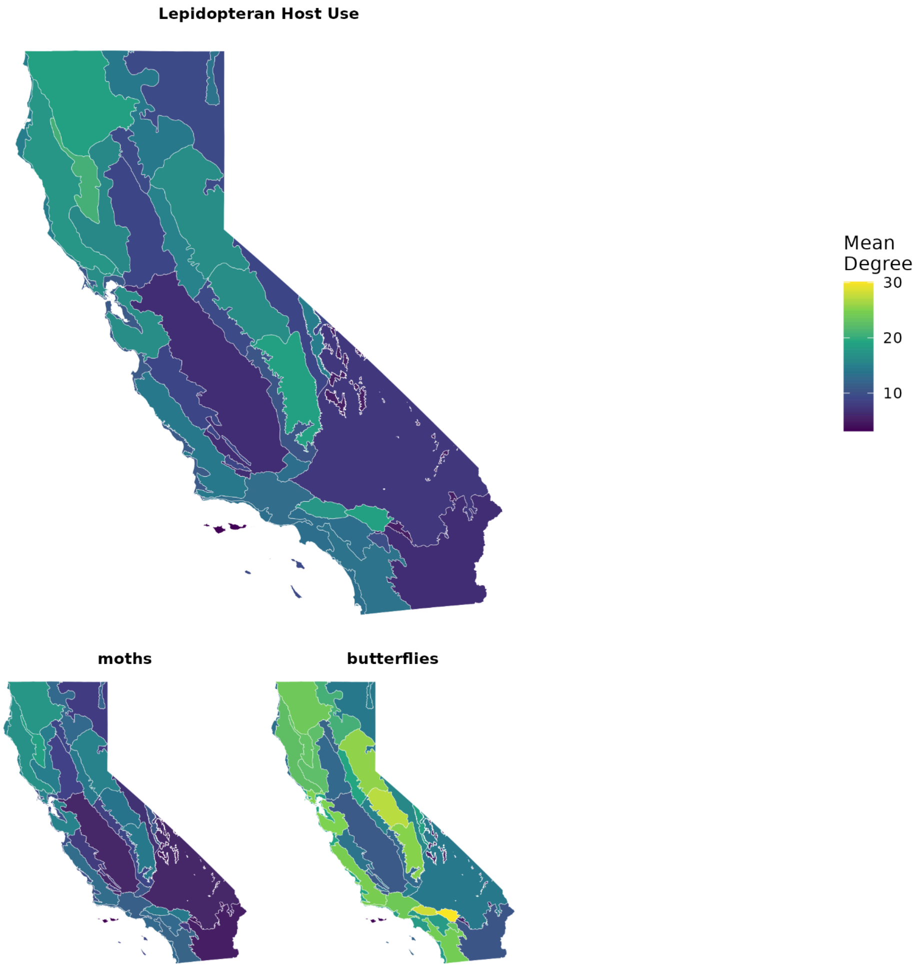


**Figure S34**. Mean number of plant partners (degree) for each ecoregion and taxonomic group from our predicted Lepidopteran host use networks.

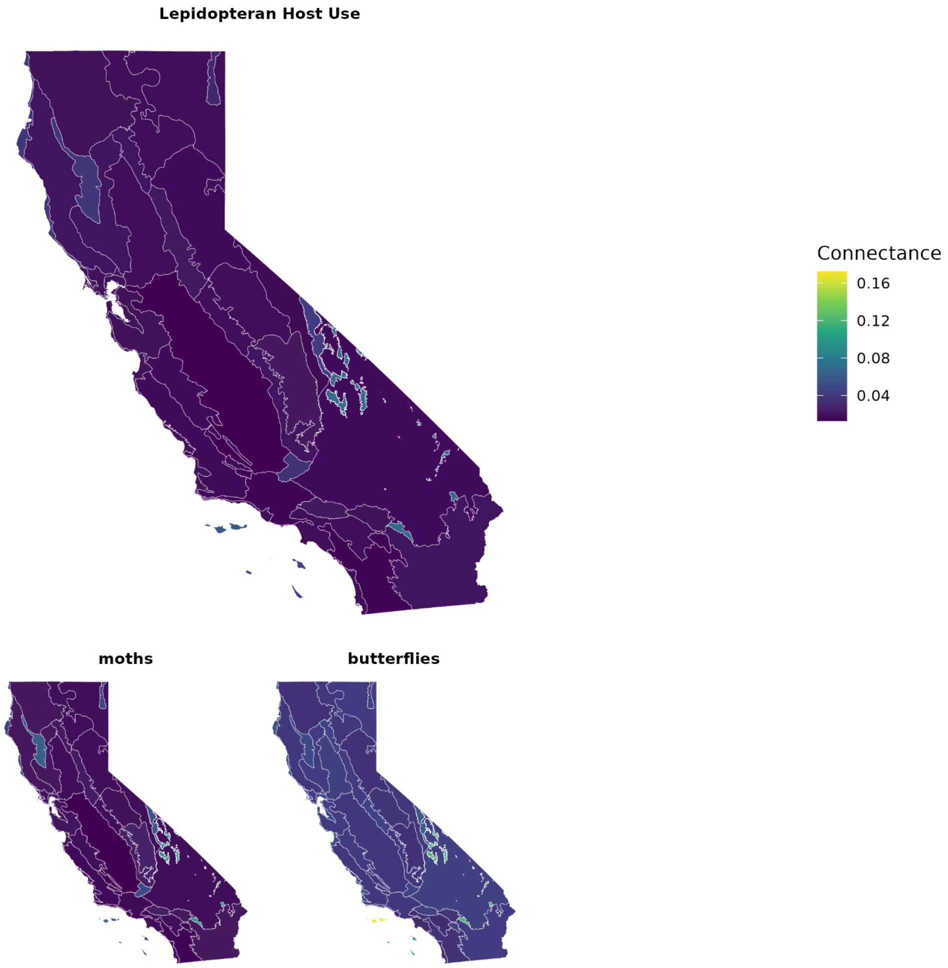


**Figure S35**. Network density of reconstructed bipartite plant-pollinator networks (connectance) for each ecoregion and taxonomic group from our predicted Lepidopteran host use networks.

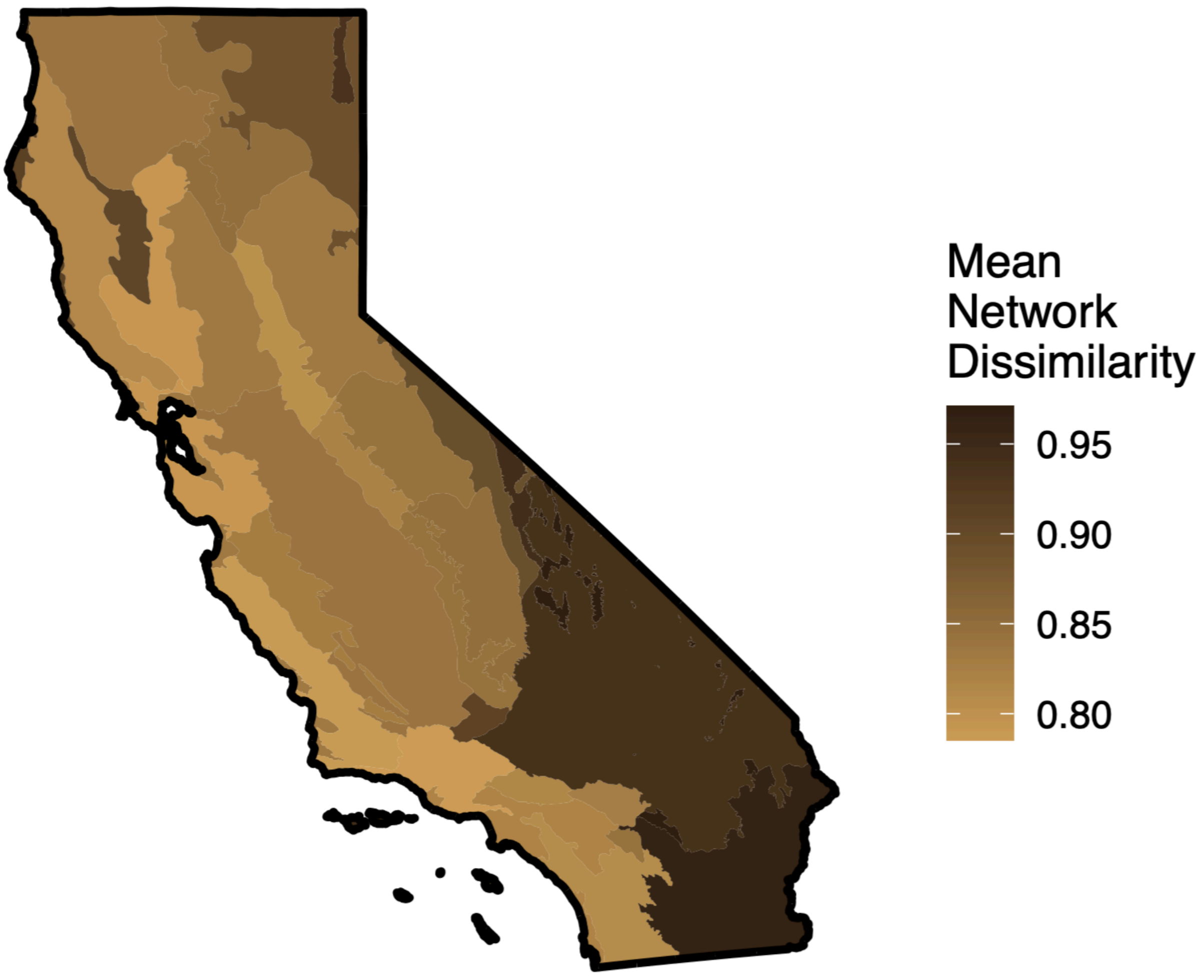


**Figure S36**. Mean dissimilarity of predicted flower visitation interaction networks (average of all pairwise dissimilarities with other regions).

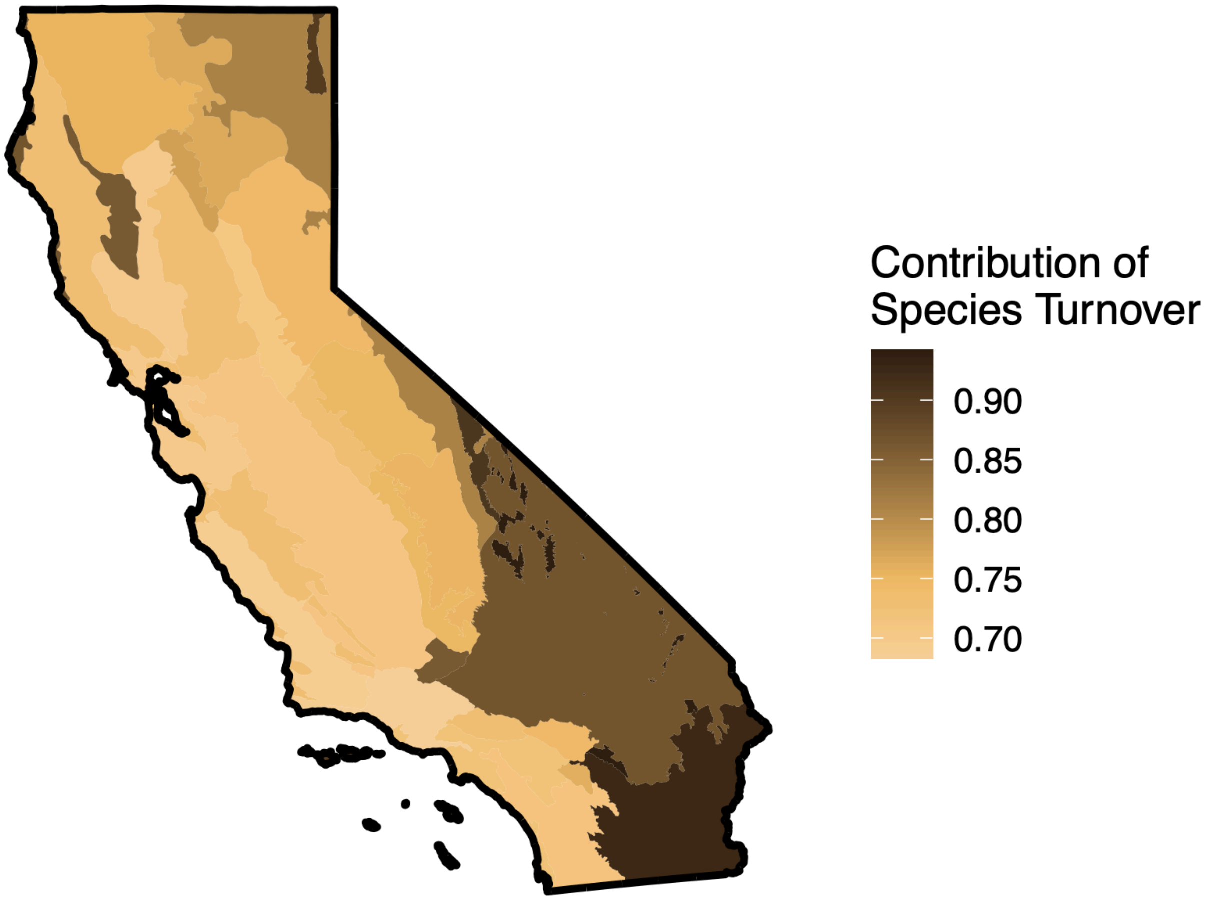


**Figure S37**. Contribution of species turnover to mean dissimilarity of predicted flower visitation interaction networks (average of all pairwise dissimilarities with other regions).

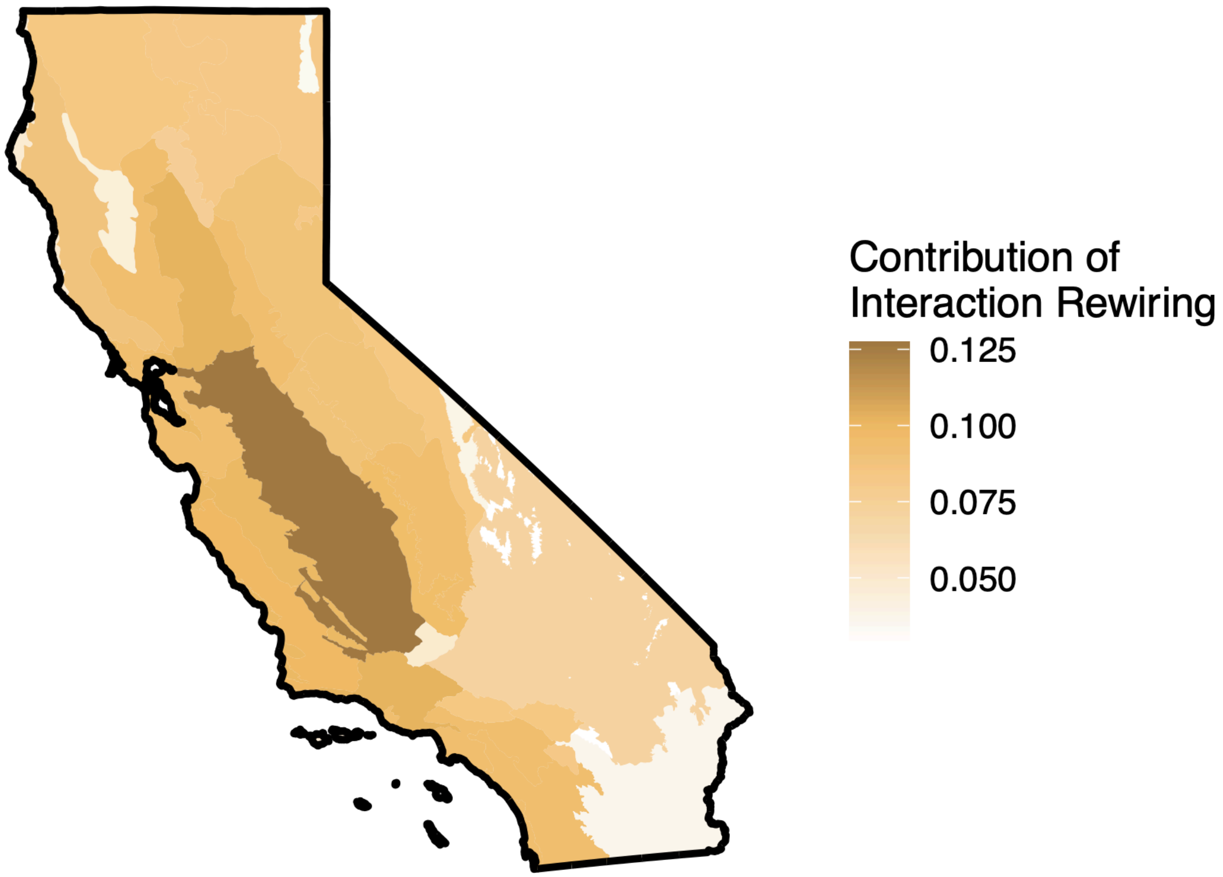


**Figure S38**. Contribution of network rewiring to mean dissimilarity of predicted flower visitation interaction networks (average of all pairwise dissimilarities with other regions).

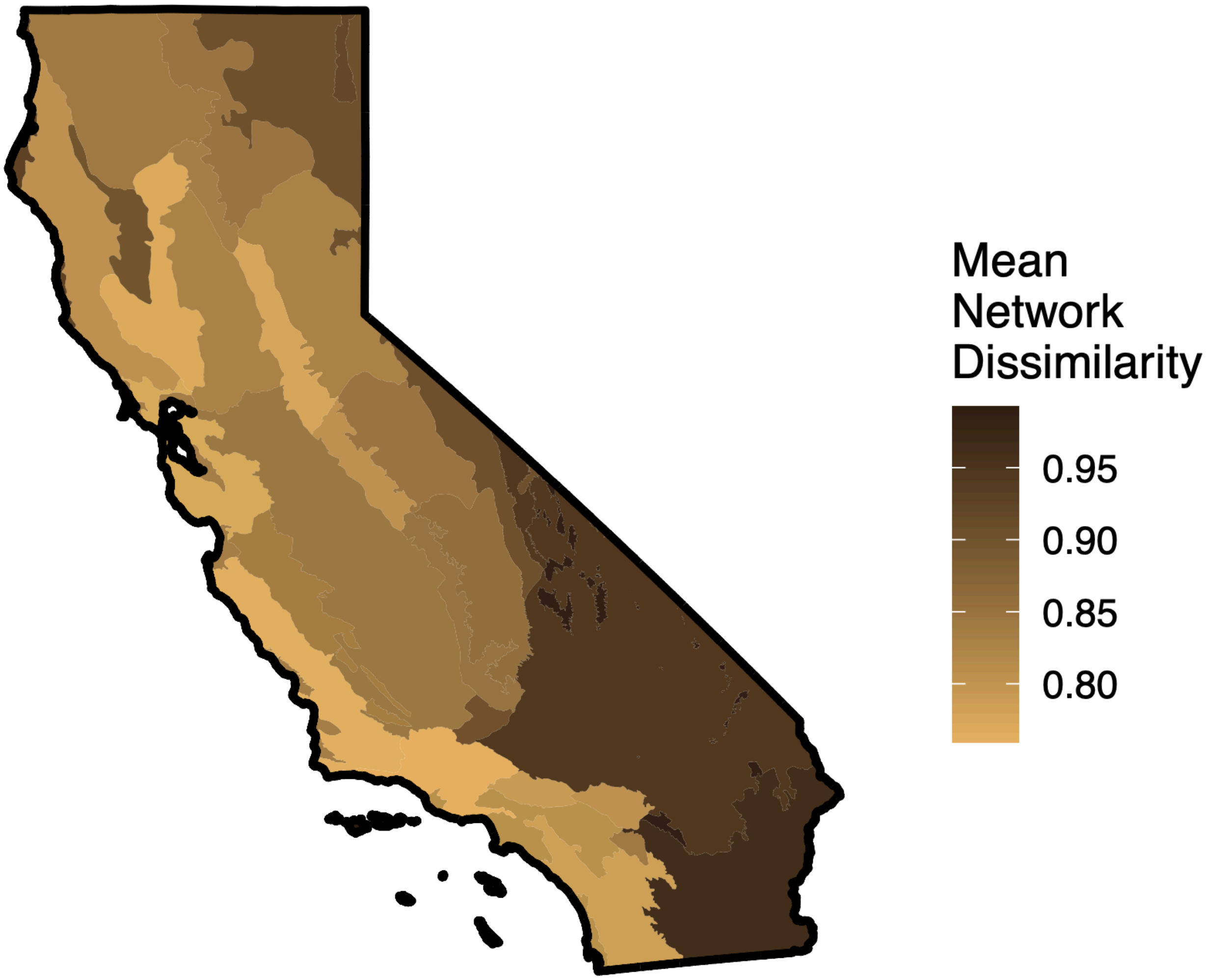


**Figure S39**. Mean dissimilarity of predicted Lepidopteran host use networks (average of all pairwise dissimilarities with other regions).

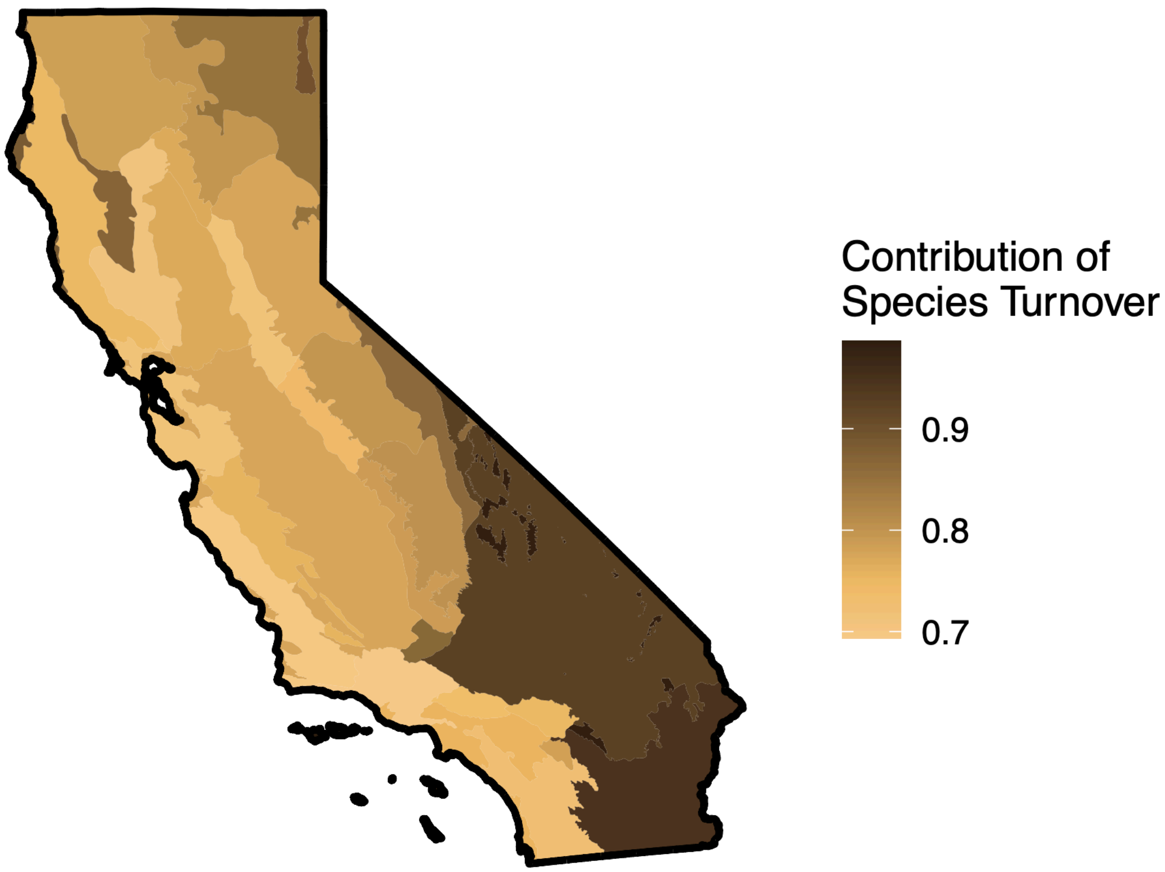


**Figure S40**. Contribution of species turnover to mean dissimilarity of predicted Lepidopteran host use interaction networks (average of all pairwise dissimilarities with other regions).

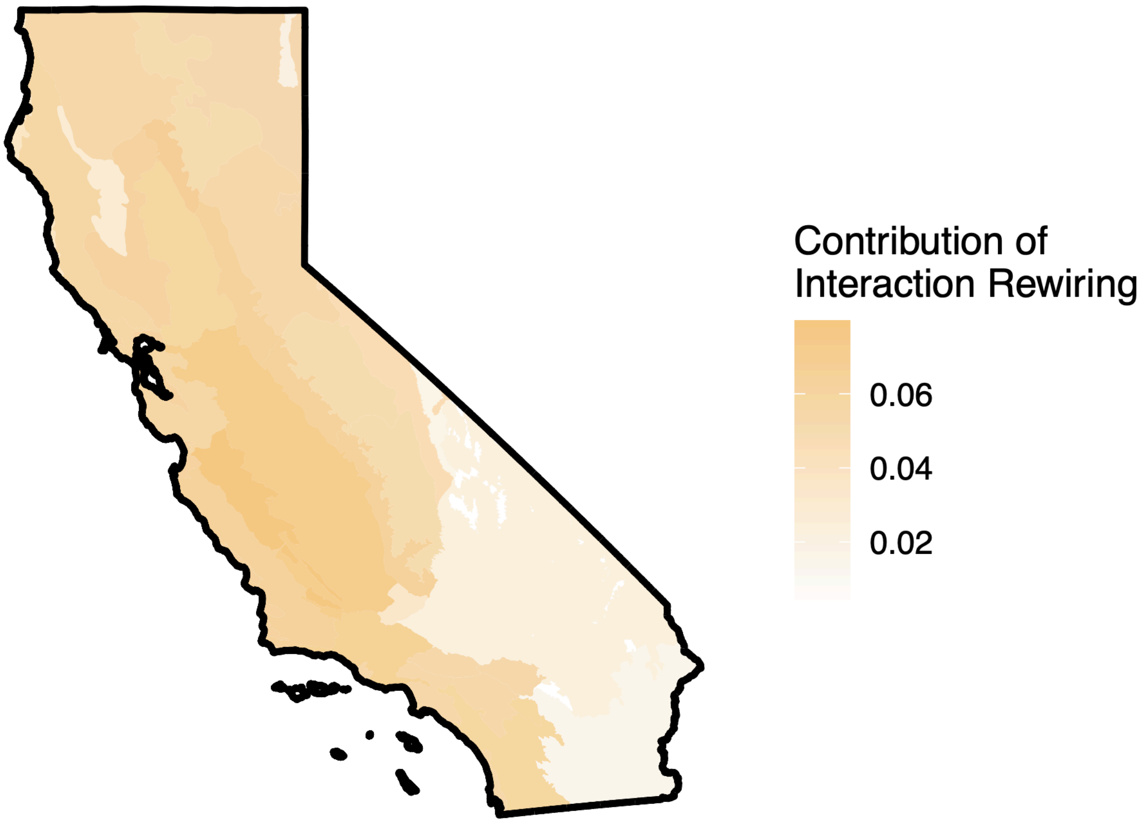


**Figure S41**. Contribution of network rewiring to mean dissimilarity of predicted Lepidopteran host use networks (average of all pairwise dissimilarities with other regions).

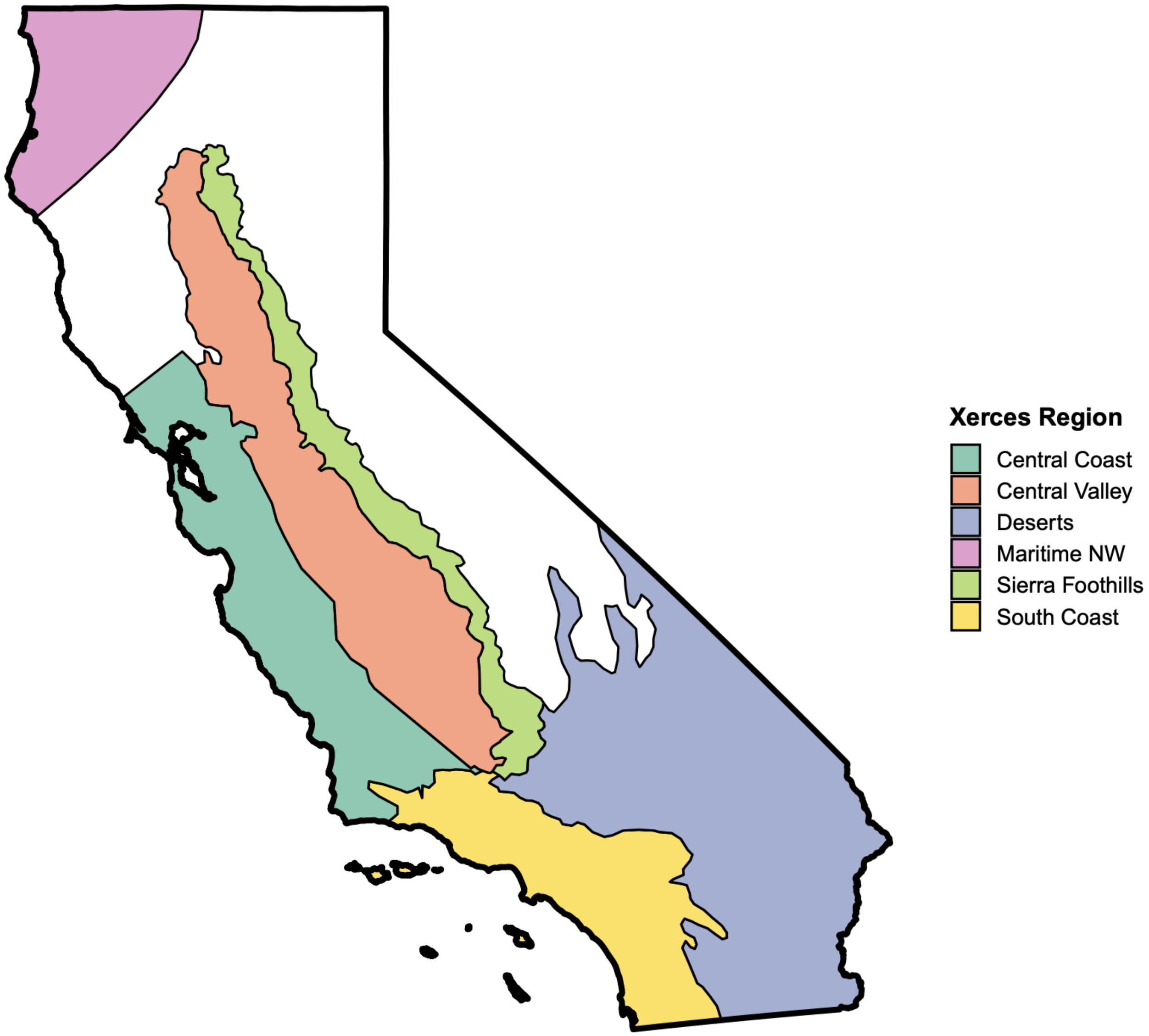


**Figure S42**. Digitized boundaries of Xerces ecoregional planting lists for California.

**Table S1.** Spatial cross-validation performance of plant and insect species distribution models. Values are medians [interquartile ranges] of species-level continuous Boyce index (CBI), area under the receiver operating characteristic curve (AUC), and maximum true skill statistic (TSS). Percentages indicate the proportion of species meeting descriptive reference benchmarks of CBI ≥ 0.25, AUC ≥ 0.70, and TSS ≥ 0.40. ESM = ensemble of small models; SDM = standard ensemble species distribution model.

| Taxon | Model | N | CBI median [IQR] | CBI >=0.25 (%) | AUC median [IQR] | AUC >=0.70 (%) | TSS median [IQR] | TSS >=0.40 (%) |
|---|---|---|---|---|---|---|---|---|
| Plants | ESM | 1363 | 0.798 [0.681-0.881] | 96.6 | 0.873 [0.787-0.929] | 89.7 | 0.681 [0.541-0.785] | 91.5 |
| Plants | SDM | 5283 | 0.772 [0.644-0.882] | 99.4 | 0.901 [0.865-0.931] | 99.7 | 0.723 [0.653-0.798] | 99.8 |
| Bees | ESM | 491 | 0.656 [0.403-0.772] | 83.1 | 0.699 [0.598-0.767] | 49.5 | 0.415 [0.286-0.525] | 53 |
| Bees | SDM | 525 | 0.692 [0.579-0.776] | 97.9 | 0.795 [0.744-0.840] | 89.3 | 0.558 [0.481-0.633] | 91.6 |
| Butterflies | ESM | 139 | 0.650 [0.416-0.798] | 83.5 | 0.711 [0.580-0.819] | 52.5 | 0.435 [0.281-0.587] | 57.6 |
| Butterflies | SDM | 279 | 0.804 [0.657-0.917] | 99.6 | 0.834 [0.790-0.879] | 96.8 | 0.578 [0.509-0.690] | 97.1 |
| Hoverflies | ESM | 33 | 0.672 [0.392-0.752] | 81.8 | 0.725 [0.617-0.806] | 57.6 | 0.467 [0.316-0.559] | 60.6 |
| Hoverflies | SDM | 57 | 0.813 [0.657-0.910] | 98.2 | 0.848 [0.773-0.893] | 93 | 0.611 [0.488-0.701] | 93 |

| | | | | | | | | |
|---|---|---|---|---|---|---|---|---|
| Moths | ESM | 1405 | 0.492 [0.131-0.701] | 67.8 | 0.615 [0.523-0.703] | 25.8 | 0.301 [0.188-0.415] | 27.5 |
| Moths | SDM | 1010 | 0.644 [0.544-0.731] | 98.4 | 0.748 [0.692-0.803] | 71.2 | 0.494 [0.417-0.586] | 81.2 |

**Table S2.** Improvement in expected pollinator community support from plants drawn from top plants identified by NECTAR at random and through complementary selection, using our predicted interaction networks as the baseline for calculating pollinator support.

| Focal Taxa | Plant Selection Method | Mean Proportion Pollinator Community Supported (95% CrI) |
|---|---|---|
| All Pollinators | Random | 0.229 (0.206, 0.254) |
| All Pollinators | External | 0.252 (0.135, 0.439) |
| All Pollinators | Raw metaweb, top draws | 0.390 (0.357, 0.422) |
| All Pollinators | NECTAR, top draws | 0.504 (0.469, 0.537) |
| All Pollinators | NECTAR, complementary selection | 0.599 (0.553, 0.643) |
| Bees | Random | 0.262 (0.232, 0.297) |
| Bees | External | 0.284 (0.160, 0.457) |
| Bees | Raw metaweb, top draws | 0.434 (0.395, 0.477) |
| Bees | NECTAR, top draws | 0.541 (0.501, 0.583) |
| Bees | NECTAR, complementary selection | 0.661 (0.610, 0.709) |
| Butterflies | Random | 0.299 (0.261, 0.341) |
| Butterflies | External | 0.339 (0.210, 0.502) |
| Butterflies | Raw metaweb, top draws | 0.536 (0.489, 0.583) |
| Butterflies | NECTAR, top draws | 0.632 (0.586, 0.675) |
| Butterflies | NECTAR, complementary selection | 0.757 (0.706, 0.803) |

| | | |
|---|---|---|
| Moths | Random | 0.201 (0.177, 0.227) |
| Moths | External | 0.226 (0.119, 0.417) |
| Moths | Raw metaweb, top draws | 0.355 (0.20, 0.392) |
| Moths | NECTAR, top draws | 0.505 (0.465, 0.543) |
| Moths | NECTAR, complementary selection | 0.608 (0.556, 0.659) |
| Hoverflies | Random | 0.223 (0.183, 0.269) |
| Hoverflies | External | 0.267 (0.156, 0.426) |
| Hoverflies | Raw metaweb, top draws | 0.381 (0.323, 0.440) |
| Hoverflies | NECTAR, top draws | 0.553 (0.490, 0612) |
| Hoverflies | NECTAR, complementary selection | 0.702 (0.634, 0.763) |

**Table S3.** Improvement in expected pollinator community support from plants drawn from top plants identified by NECTAR at random and through complementary selection, using a raw metaweb as the baseline for calculating pollinator support.

| Focal Taxa | Plant Selection Method | Mean Proportion Pollinator Community Supported (95% CrI) |
|---|---|---|
| All Pollinators | Random | 0.041 (0.035, 0.048) |
| All Pollinators | External | 0.130 (0.094, 0.184) |
| All Pollinators | Raw metaweb, top draws | 0.211 (0.185, 0.241) |
| All Pollinators | NECTAR, top draws | 0.099 (0.085, 0.115) |
| All Pollinators | NECTAR, complementary selection | 0.133 (0.110, 0.160) |
| Bees | Random | 0.101 (0.092, 0.111) |
| Bees | External | 0.305 (0.210, 0.414) |
| Bees | Raw metaweb, top draws | 0.506 (0.481, 0.533) |
| Bees | NECTAR, top draws | 0.235 (0.217, 0.254) |

| | | |
|---|---|---|
| Bees | NECTAR, complementary selection | 0.260 (0.222, 0.297) |
| Butterflies | Random | 0.107 (0.097, 0.118) |
| Butterflies | External | 0.350 (0.270, 0.447) |
| Butterflies | Raw metaweb, top draws | 0.608 (0.581, 0.633) |
| Butterflies | NECTAR, top draws | 0.393 (0.367, 0.419) |
| Butterflies | NECTAR, complementary selection | 0.325 (0.278, 0.374) |
| Moths | Random | 0.003 (0.002, 0.004) |
| Moths | External | 0.008 (0.007, 0.010) |
| Moths | Raw metaweb, top draws | 0.030 (0.025, 0.038) |
| Moths | NECTAR, top draws | 0.008 (0.007, 0.010) |
| Moths | NECTAR, complementary selection | 0.020 (0.015, 0.027) |
| Hoverflies | Random | 0.047 (0.041, 0.057) |
| Hoverflies | External | 0.149 (0.086, 0.257) |
| Hoverflies | Raw metaweb, top draws | 0.421 (0.384, 0.469) |
| Hoverflies | NECTAR, top draws | 0.182 (0.160, 0.212) |
| Hoverflies | NECTAR, complementary selection | 0.150 (0.113, 0.192) |

**Table S4.** Number of plant-pollinator-ecoregion flower visitation interactions predicted for each threshold used in the interaction plausibility threshold sensitivity analysis.

| Threshold used (percentile of previously observed interactions) | Total flower visitation interactions predicted | Flower visitation interactions predicted where link was identified as potential with phylogenetic constraint | Flower visitation interactions predicted for pollinators with no pre-existing data where link was identified as potential with graph embedding |
|---|---|---|---|
| 10th | 4,154,059 | 1,583,336 | 2,570,723 |
| 15th | 3,358,107 | 1,307,140 | 2,050,967 |
| 20th | 2,778,188 | 1,097,591 | 1,680,597 |

| | | | |
|---|---|---|---|
| 25th (default) | 2,256,597 | 899,129 | 1,357,468 |
| 30th | 1,955,907 | 790,258 | 1,165,649 |
| 35th | 1,664,423 | 678,006 | 986,417 |
| 40th | 1,423,447 | 583,435 | 840,012 |

**Table S5.** The five plant genera most commonly appearing in the top 25 plant species in ranked ecoregion specific plant lists based on predicted flower visitation interactions. This includes both the total number of species by ecoregion (out of 35 ecoregions), or number of distinct species, or number of ecoregions.

| Plant Genus | Number of total appearances in top 25 plant lists (flower visitation) | Number of distinct species appearing in top 25 plant lists (flower visitation) | Number of ecoregions genus appeared in top 25 plant lists (flower visitation) | Supporting References |
|---|---|---|---|---|
| *Eriogonum* (Wild buckwheats) | 99 | 48 | 22 | Meiners, J. M., Griswold, T. L., & Carril, O. M. (2019). Decades of native bee biodiversity surveys at Pinnacles National Park highlight the importance of monitoring natural areas over time. PLoS One, 14(1), e0207566.<br><br>Reyes, E., Smallwood, N. & Wood, E. M. Native-plant landscaping enhances floral visitation by insects and birds in residential yards. *Urban Ecosyst.* **29**, (2026). |

| | | | | |
|---|---|---|---|---|
| *Asclepias* (Milkweed) | 80 | 10 | 29 | Baker, A. M., & Potter, D. A. (2018). Colonization and usage of eight milkweed (Asclepias) species by monarch butterflies and bees in urban garden settings. *Journal of Insect Conservation*, *22*(3), 405-418.<br><br>Jett, I. D., & Yang, L. H. (2026). Trapped honey bees reduce floral visitation on milkweed flowers. Ecology, 107(7), e70452. |
| *Ranunculus* (Buttercups) | 75 | 16 | 27 | Meiners, J. M., Griswold, T. L., & Carril, O. M. (2019). Decades of native bee biodiversity surveys at Pinnacles National Park highlight the importance of monitoring natural areas over time. PLoS One, 14(1), e0207566. |
| *Ceanothus* (California lilac) | 71 | 19 | 25 | Nabors, A., Hung, K. L. J., Corkidi, L., & Bethke, J. A. (2022). California native perennials attract greater native pollinator abundance and diversity than nonnative, commercially available ornamentals in Southern California. *Environmental* |

| | | | | |
|---|---|---|---|---|
| | | | | *entomology*, *51*(4), 836-847.<br><br>Frankie, G., Feng, I., Thorp, R., Pawelek, J., Chase, M. H., Jadallah, C. C., & Rizzardi, M. (2019). Native and non-native plants attract diverse bees to urban gardens in California. Journal of Pollination Ecology, 25. |
| *Trifolium* (Clover) | 61 | 19 | 22 | Cole, J. S., Siegel, R. B., Loffland, H. L., Elsey, E. A., Tingley, M. W., & Johnson, M. (2020). Plant selection by bumble bees (Hymenoptera: Apidae) in montane riparian habitat of California. *Environmental Entomology*, *49*(1), 220-229. |

**Table S6.** The five plant genera most commonly appearing in the top 25 plant species in ranked ecoregion specific plant lists based on predicted Lepidopteran host interactions. This includes both the total number of species by ecoregion (out of 35 ecoregions), or number of distinct species, or number of ecoregions.

| Plant Genus | Number of total appearances in top 25 plant lists (Lepidopteran host use) | Number of distinct species appearing in top 25 plant lists (Lepidopteran host use) | Number of ecoregions genus appeared in top 25 plant lists (Lepidopteran host use) | Supporting References |
|---|---|---|---|---|

| | | | | |
|---|---|---|---|---|
| *Quercus* (Oaks) | 181 | 19 | 28 | Narango, D. L., Tallamy, D. W., & Shropshire, K. J. (2020). Few keystone plant genera support the majority of Lepidoptera species. *Nature communications*, *11*(1), 5751. |
| *Salix* (Willows) | 168 | 20 | 31 | Narango, D. L., Tallamy, D. W., & Shropshire, K. J. (2020). Few keystone plant genera support the majority of Lepidoptera species. *Nature communications*, *11*(1), 5751. |
| *Ceanothus* (California lilac) | 105 | 23 | 24 | Philbin, C. S., Paulsen, M., & Richards, L. A. (2021). Opposing effects of Ceanothus velutinus phytochemistry on herbivore communities at multiple scales. Metabolites, 11(6), 361. |
| *Eriogonum* (Wild Buckwheats) | 77 | 39 | 9 | Graves, S. D., & Shapiro, A. M. (2003). Exotics as host plants of the California butterfly fauna. Biological conservation, 110(3), 413-433. |
| *Prunus* (Stone Fruit) | 62 | 7 | 29 | Narango, D. L., Tallamy, D. W., & Shropshire, K. J. (2020). Few keystone plant genera support the majority of Lepidoptera |

| | | | | |
|---|---|---|---|---|
| | | | | species. *Nature communications*, *11*(1), 5751. |

**Graph embedding**

**Table S7.** Classification thresholds for predicting bee species-plant genus interactions via transfer learning, selected by maximizing Youden's J statistic (i.e., maximizing the sum of true positive and true negative rates) on interactions of held-out pollinator species (90% in training, 10% in validation). Threshold selection treats unobserved interactions as non-interactions, which may include both true non-interactions and undetected interactions. We report the threshold value and recall rate (true positives / (true positives + false negatives)) for each fold of the 10-fold cross-validation.

| Fold | Threshold | Recall Rate |
|---|---|---|
| 1 | 0.00197 | 0.426 |
| 2 | 0.000474 | 0.457 |
| 3 | 0.00207 | 0.291 |
| 4 | 0.000131 | 0.446 |
| 5 | 0.000725 | 0.429 |
| 6 | 0.00134 | 0.408 |
| 7 | 0.000534 | 0.420 |
| 8 | 0.00110 | 0.405 |
| 9 | 0.000105 | 0.550 |
| 10 | 0.00191 | 0.384 |

**Table S8.** Classification thresholds for predicting hoverfly species-plant genus interactions via transfer learning, selected by maximizing Youden's J statistic (i.e., maximizing the sum of true positive and true negative rates) on interactions of held-out pollinator species (90% in training, 10% in validation). Threshold selection treats unobserved interactions as non-interactions, which may include both true non-interactions and undetected interactions. We report the threshold value and recall rate (true positives / (true positives + false negatives)) for each fold of the 10-fold cross-validation.

| Fold | Threshold | Recall Rate |
|---|---|---|
| 1 | 0.00326 | 0.314 |
| 2 | 0.00232 | 0.381 |
| 3 | 0.00213 | 0.381 |
| 4 | 0.00203 | 0.344 |
| 5 | 0.00259 | 0.527 |
| 6 | 0.00528 | 0.359 |
| 7 | 0.000674 | 0.528 |
| 8 | 0.00360 | 0.230 |
| 9 | 0.000436 | 0.369 |
| 10 | 0.00198 | 0.319 |

**Table S9.** Classification thresholds for predicting butterfly species-plant genus interactions via transfer learning, selected by maximizing Youden's J statistic (i.e., maximizing the sum of true positive and true negative rates) on interactions of held-out pollinator species (90% in training, 10% in validation). Threshold selection treats unobserved interactions as non-interactions, which may include both true non-interactions and undetected interactions. We report the threshold value and recall rate (true positives / (true positives + false negatives)) for each fold of the 10-fold cross-validation.

| Fold | Threshold | Recall Rate |
|---|---|---|
| 1 | 0.00135 | 0.714 |
| 2 | 0.00239 | 0.765 |
| 3 | 0.00246 | 0.694 |
| 4 | 0.00381 | 0.635 |
| 5 | 0.00191 | 0.714 |
| 6 | 0.00250 | 0.673 |
| 7 | 0.00306 | 0.755 |
| 8 | 0.00245 | 0.628 |
| 9 | 0.00389 | 0.728 |
| 10 | 0.0498 | 0.331 |

**Table S10.** Classification thresholds for predicting moth species-plant genus interactions via transfer learning, selected by maximizing Youden's J statistic (i.e., maximizing the sum of true positive and true negative rates) on interactions of held-out pollinator species (90% in training, 10% in validation). Threshold selection treats unobserved interactions as non-interactions, which may include both true non-interactions and undetected interactions. We report the threshold value and recall rate (true positives / (true positives + false negatives)) for each fold of the 10-fold cross-validation.

| Fold | Threshold | Recall Rate |
|---|---|---|
| 1 | 0.00204 | 0.418 |
| 2 | 0.00142 | 0.500 |

| | | |
|---|---|---|
| 3 | 0.000459 | 0.660 |
| 4 | 0.00151 | 0.440 |
| 5 | 0.00143 | 0.569 |
| 6 | 0.00142 | 0.446 |
| 7 | 0.000547 | 0.648 |
| 8 | 0.00115 | 0.600 |
| 9 | 0.000912 | 0.605 |
| 10 | 0.000530 | 0.667 |

**Table S11.** Performance of transfer learning predictions for bee species-plant genus interactions using the average classification threshold selected in Table S1. We report recall rate (true positives / (true positives + false negatives)) for each fold of the 3-fold cross-validation on interactions of held-out pollinator species (90% in training, 10% in validation).

| Fold | Threshold | Recall Rate |
|---|---|---|
| 1 | 0.00104 | 0.402 |
| 2 | 0.00104 | 0.406 |
| 3 | 0.00104 | 0.431 |

**Table S12.** Performance of transfer learning predictions for hoverfly species-plant genus interactions using the average classification threshold selected in Table S2. We report recall rate (true positives / (true positives + false negatives)) for each fold of the 3-fold cross-validation on interactions of held-out pollinator species (90% in training, 10% in validation).

| Fold | Threshold | Recall Rate |
|---|---|---|
| 1 | 0.00243 | 0.224 |

| | | |
|---|---|---|
| 2 | 0.00243 | 0.241 |
| 3 | 0.00243 | 0.305 |

**Table S13.** Performance of transfer learning predictions for butterfly species-plant genus interactions using the average classification threshold selected in Table S3. We report recall rate (true positives / (true positives + false negatives)) for each fold of the 3-fold cross-validation on interactions of held-out pollinator species (90% in training, 10% in validation).

| Fold | Threshold | Recall Rate |
|---|---|---|
| 1 | 0.00737 | 0.573 |
| 2 | 0.00737 | 0.559 |
| 3 | 0.00737 | 0.597 |

**Table S14.** Performance of transfer learning predictions for moth species-plant genus interactions using the average classification threshold selected in Table S4. We report recall rate (true positives / (true positives + false negatives)) for each fold of the 3-fold cross-validation on interactions of held-out pollinator species (90% in training, 10% in validation).

| Fold | Threshold | Recall Rate |
|---|---|---|
| 1 | 0.00114 | 0.532 |
| 2 | 0.00114 | 0.461 |
| 3 | 0.00114 | 0.478 |

**Table S15.** A summary table of the graph embedding interaction prediction results. Phylogenetic coverage is calculated based on how many checklist species have phylogenetic information from the Open Tree of Life[1]. The total number of species used in transfer learning is the number of checklist species that have such phylogenetic information.

| Taxon | Type of learning | Total Number of Species in Checklist | Phylogenetic Coverage | Total Number of Species Used in Transfer Learning | Total Number of Interaction Gained | Number of Interaction Gained per Species |
|---|---|---|---|---|---|---|
| Bees | In-sample learning | 1610 | 0.991 | 1596 | 124800 | 78.195 |
| | Out-of-sample learning | | | | 13908 | 8.714 |
| Hoverflies | In-sample learning | 214 | 0.874 | 187 | 4129 | 22.080 |
| | Out-of-sample learning | | | | 2376 | 12.706 |
| Butterflies | In-sample learning | 281 | 0.847 | 238 | 18389 | 77.265 |
| | Out-of-sample learning | | | | 3682 | 15.471 |
| Moths | In-sample learning | 4137 | 0.899 | 3721 | 18526 | 4.979 |
| | Out-of-sample learning | | | | 221098 | 59.419 |
